# De novo lipogenesis controls antigen presentation and metastatic immune escape in triple-negative breast cancer

**DOI:** 10.64898/2026.05.21.726063

**Authors:** Myung Soo Ko, Henry Withers, Divya Ramchandani, Jason McCormick, Sebastian E. Carrasco, Pius Amartey, Arshdeep Singh, Liron Yoffe, Glenn E. Simmons, Guoan Zhang, Wen H. Shen, Vivek Mittal

**Affiliations:** Department of Cardiothoracic Surgery, Weill Cornell Medicine, New York, NY, USA; Neuberger Berman Lung Cancer Center, Weill Cornell Medicine, New York, NY, USA; Department of Cell and Developmental Biology, Weill Cornell Medicine, New York, NY, USA; Department of Biostatistics and Bioinformatics, Roswell Park Comprehensive Cancer Center, Buffalo, New York; Flow Cytometry Core Facility, Weill Cornell Medicine, New York, NY, USA; Laboratory of Comparative Pathology, Memorial Sloan Kettering Cancer Center, Weill Cornell Medicine, Rockefeller University, New York, NY, USA; Department of Biomedical Sciences, College of Veterinary Medicine, Cornell University, Ithaca, USA; Proteomics and Metabolomics Core Facility, Weill Cornell Medicine, New York, NY, USA; Department of Radiation Oncology, Weill Cornell Medicine, New York, NY, USA; Sandra and Edward Meyer Cancer Center, Weill Cornell Medicine, New York, NY, USA; Caryl and Israel Englander Institute for Precision Medicine, New York, NY, USA

## Abstract

Metastatic dissemination is a major cause of mortality in triple-negative breast cancer (TNBC), yet how tumor metabolic state regulates immune surveillance during metastasis remains poorly understood. Here, we identify de novo lipogenesis as a tumor-intrinsic determinant of immune visibility and metastatic progression. Genetic or pharmacologic disruption of fatty acid synthase (FASN) suppresses metastasis with limited effects on primary tumor growth and enhances tumor-cell major histocompatibility complex class I (MHC-I) antigen presentation and CD8^+^ T cell activation. Mechanistically, FASN-dependent lipogenesis sustains epidermal growth factor receptor (EGFR) palmitoylation, selectively maintaining PI3K−AKT−mTOR signaling without altering EGFR abundance, cell-surface localization, or MAPK activation. Disruption of FASN-dependent lipogenesis, EGFR palmitoylation, or mTOR signaling restores peptide–MHC-I presentation and enhances antigen-specific CD8^+^ T cell recognition and cytotoxicity. FASN loss increases the abundance of multiple antigen-processing machinery proteins without corresponding changes in MHC-I transcript abundance, supporting transcription-independent regulation of antigen presentation downstream of PI3K−AKT−mTOR signaling. Tumor-cell B2M depletion substantially restores metastatic outgrowth and circulating tumor cells following FASN loss, identifying MHC-I antigen presentation as a major mediator of lipogenesis-dependent metastatic immune escape. In patient-derived TNBC organoids and independent human breast cancer cohorts, elevated lipogenic gene expression correlated with PI3K−AKT−mTOR-related transcriptional programs and reduced immune-surveillance programs, without significant differences in MHC-I transcriptional signatures. Together, these findings define a tumor-intrinsic immunometabolic circuit in which de novo lipogenesis sustains EGFR palmitoylation and PI3K−AKT−mTOR signaling to regulate antigen presentation, thereby modulating tumor-cell immune visibility during metastatic progression.

## Introduction

Triple-negative breast cancer (TNBC) is an aggressive breast cancer subtype characterized by a high risk of metastatic recurrence and limited therapeutic options^1^. Although advances in systemic therapy have improved outcomes^2^, metastatic disease remains the principal cause of mortality. Successful metastatic progression requires tumor cells to survive and adapt to distinct tissue environments while evading immune surveillance. However, the tumor-intrinsic mechanisms that coordinate metabolic adaptation with immune evasion during metastatic progression remain incompletely understood.

Immune checkpoint blockade (ICB) has improved outcomes for a subset of patients with TNBC, but durable responses remain limited, in part because tumor cells can evade recognition and elimination by cytotoxic CD8^+^ T cells^3^. Effective CD8^+^ T-cell surveillance requires presentation of tumor-derived peptides by major histocompatibility complex class I (MHC-I) molecules. Accordingly, defects in MHC-I antigen presentation represent an important mechanism of tumor immune escape. Although transcriptional, genetic, and signaling mechanisms that regulate MHC-I expression have been described, how tumor cells couple their metabolic state to antigen presentation and immune recognition during metastatic progression remains poorly understood.

Metabolic reprogramming is a hallmark of TNBC and contributes to tumor progression, therapeutic resistance, and adaptation to metastatic environments^4^. Among these adaptations, de novo lipogenesis can promote metastatic fitness, including adaptation of breast cancer cells to lipid-limited metastatic niches such as the brain^5,6^. Fatty acid synthase (FASN), a central enzyme in de novo fatty acid synthesis, is frequently upregulated in aggressive cancers and associated with poor clinical outcomes^7^. Beyond providing lipids for membrane and energy metabolism, FASN generates palmitate that can support protein S-palmitoylation, a reversible lipid modification that regulates the localization, stability, trafficking, and signaling^8,9^. Whether FASN-dependent lipogenesis regulates tumor antigen presentation and immune surveillance through protein S-palmitoylation during metastatic progression remains unknown.

Here, we identify de novo lipogenesis as a tumor-intrinsic regulator of immune visibility during metastatic progression in TNBC. We show that FASN-dependent lipogenesis sustains S-palmitoylation of epidermal growth factor receptor (EGFR), which selectively maintains PI3K−AKT−mTOR signaling without altering EGFR abundance, cell-surface localization, or MAPK signaling. This lipid-dependent signaling state suppresses MHC-I antigen presentation through transcription-independent regulation of the antigen-processing machinery, thereby limiting CD8^+^ T-cell recognition. Disruption of the lipogenesis−EGFR palmitoylation− PI3K−AKT−mTOR axis restores tumor-cell antigen presentation and CD8⁺ T-cell recognition and constrains metastatic progression. Together, these findings reveal a lipid-dependent immunometabolic mechanism that links tumor lipid metabolism to antigen presentation and metastatic immune escape through EGFR palmitoylation and PI3K−AKT−mTOR signaling.

### De novo lipogenesis is required for metastatic progression in TNBC

Using an OCT4/SOX2-GFP reporter system^10^ in orthotopic human MDA-MB-231-LM2 and murine EO771-ML1 TNBC models, we previously identified a rare OCT4/SOX2-GFP^+^ population enriched for stemness-associated factors, including OCT4, SOX2 and NANOG, with markedly enhanced metastatic capacity in vivo. These metastasis-competent cells exhibit aberrant EZH2 catalytic activity^11^, chromosomal instability (CIN)^12^ and dysregulated copper homeostasis^13^. Whether metabolic reprogramming contributes to this state, particularly through mechanisms involving tumor–immune interactions, remains unknown.

RNA-seq analysis of sorted OCT4/SOX2-GFP^+^ metastasis-competent cells isolated from orthotopic primary tumors revealed coordinated enrichment of lipogenic and anabolic metabolic programs, with fatty acid metabolism among the most prominent pathways, alongside reduced expression of immune activation programs (**Fig. 1a and Extended Data Fig. 1a,b**). qPCR confirmed increased expression of the core de novo lipogenesis enzymes ATP citrate lyase (ACLY), acetyl-CoA carboxylase (ACC1; encoded by ACACA) and FASN, accompanied by elevated expression of OCT4, SOX2 and NANOG (**Fig. 1b**).

**Fig. 1.**
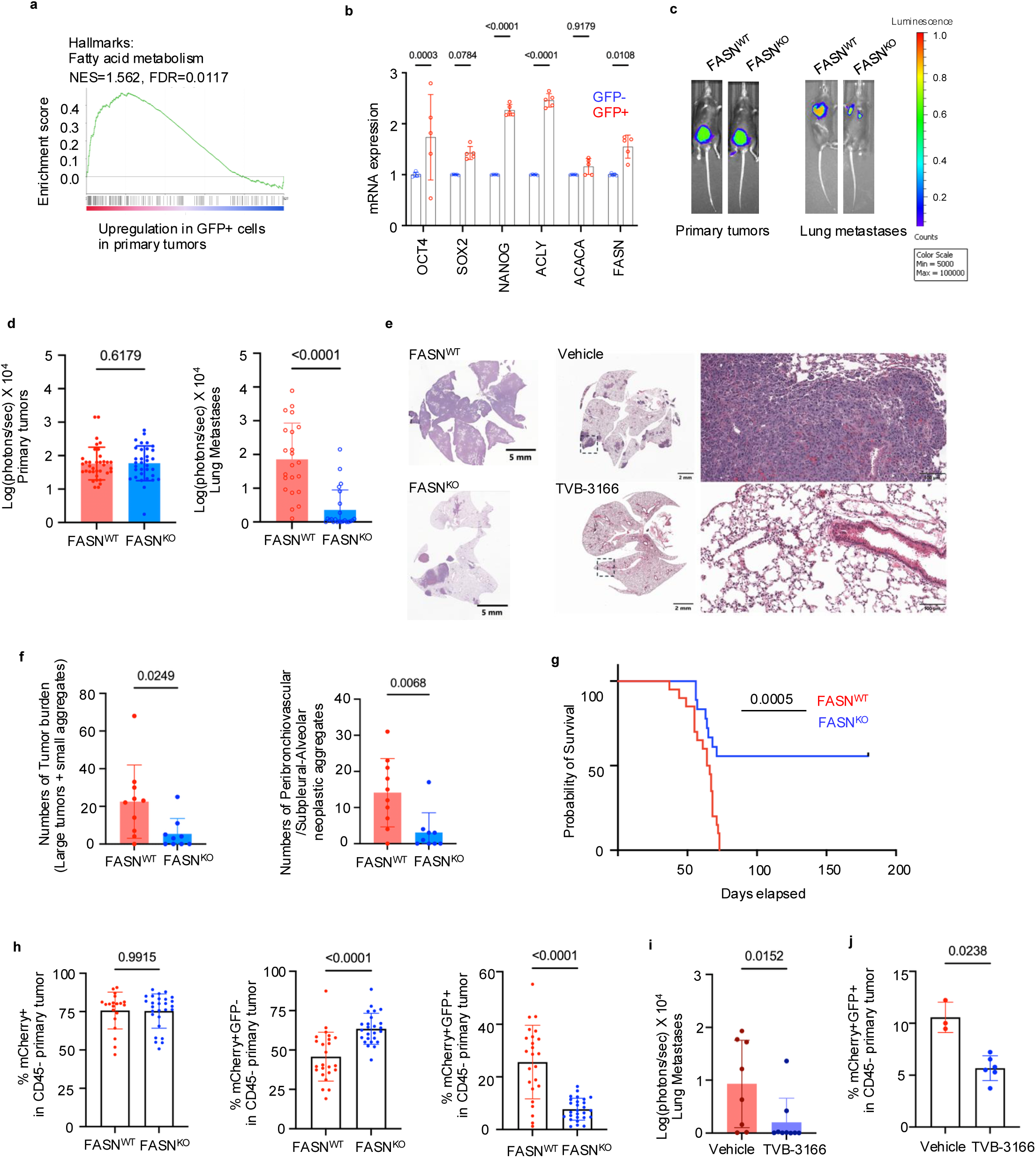
De novo lipogenesis is required for metastatic progression in TNBC. **a,** GSEA showing significant enrichment of the HALLMARK_FATTY_ACID_METABOLISM gene set in OCT4/SOX2-GFP^+^ versus GFP^-^ tumor cells isolated from orthotopic EO771-ML1 primary tumors (n = 3 biological replicates per group). **b,** RT–qPCR analysis of stemness-associated and de novo lipogenesis genes in GFP^+^ versus GFP^-^ tumor cells (n = 5 biological replicates per group). **c,** BLI of orthotopic primary tumors at week 4 and lung metastases at week 3 after primary-tumor resection. **d,** Quantification of BLI signals from primary tumors and lung metastases (n = 20 mice). **e,** Representative H&E-stained whole-lung sections showing metastatic tumor burden in mice bearing FASN^WT^ or FASN^KO^ tumors and in mice treated with vehicle or TVB-3166 at week 3 after primary-tumor resection. Boxed regions are shown at higher magnification. Scale bars, 5 mm, 2 mm and 100 μm, as indicated. **f,** Histological quantification of lung metastatic burden in FASN^WT^ and FASN^KO^ mice, including total tumor burden comprising large lesions and small neoplastic aggregates (left) and peribronchovascular, alveolar and subpleural neoplastic aggregates (right; n = 9 mice). **g,** Kaplan–Meier survival analysis (n = 15 per group). **h,** Flow-cytometric analysis of mCherry^+^ total tumor cells, GFP^-^ cells and GFP^+^ cells in orthotopic primary tumors (n = 20 tumors per group). **i,** BLI quantification of lung metastases following FASN inhibition with TVB-3166 (n = 8 mice). **j,** Flow-cytometric quantification of GFP^+^ tumor cells in primary tumors following vehicle or TVB-3166 treatment (n = 3 vehicle tumors; n = 6 TVB-3166-treated tumors).Data in b, d, f and h–j are presented as mean ± s.e.m. Statistical significance was assessed using a two-sided unpaired Student’s t-test for d, f, and h–j. For a, the normalized enrichment score (NES) and false discovery rate (FDR) are indicated. For b, ordinary two-way ANOVA with Šídák’s multiple-comparisons test was used. Survival in g was compared using a two-sided log-rank (Mantel–Cox) test (P = 0.0005). Exact P values are indicated in the figures.

To assess the relevance of this lipogenic state in human breast cancer, we analyzed integrated single-cell RNA-sequencing datasets from human breast tumors^14^. De novo lipogenesis genes, including FASN, ACACA and ACLY, were expressed across malignant epithelial populations, which also expressed stemness-associated markers POU5F1, SOX2 and NANOG (**Extended Data Fig. 1c**).

These findings identified a lipogenic, stemness-associated tumor-cell state enriched in metastasis-competent populations in experimental TNBC models, with expression of corresponding lipogenic and stemness-associated genes observed in human malignant epithelial populations. We next asked whether this lipogenic state was functionally required for metastatic progression. We focused on FASN, the rate-limiting enzyme that catalyzes de novo synthesis of palmitate.

We used CRISPR to generate FASN knockout (FASN^KO^) cells in murine EO771-ML1 and human MDA-MB-231-LM2 TNBC models (**Extended Data Fig. 1d-g**). FASN deficiency altered cell-cycle distribution but had minimal effects on in vitro cell proliferation, invasion, or growth of 3D-bioprinted spheroids (**Extended Data Fig. 1h-p**). In vivo, FASN deficiency had little effect on orthotopic primary tumor growth in syngeneic immunocompetent mice (**Fig. 1c,d, and Extended Data Fig. 2a**). In contrast, FASN deficiency markedly suppressed metastatic outgrowth. After surgical resection of primary tumors (∼1cm^3^ at week 4), FASN^KO^ tumors showed markedly reduced lung metastasis compared with FASN^WT^ controls, as measured by BLI and lung weight (**Fig. 1c,d, and Extended Data Fig. 2a,b**). Histopathology confirmed a striking reduction in both the number and size of metastatic lesions following FASN deficiency (**Fig.1e, left and f, and Extended Data Fig. 2c-h**). This reduced metastatic burden was associated with significantly improved overall survival (**Fig. 1g**).

We next asked whether disruption of de novo lipogenesis altered the abundance or functional capacity of metastasis-competent cells. FASN^KO^ tumors exhibited a significantly reduced proportion of OCT4/SOX2-GFP^+^ metastasis-competent cells (**Fig. 1h and Extended Data Fig. 2j**), while the remaining GFP⁺ tumor cells lacked detectable FASN protein (**Extended Data Fig. 2i**), supporting a role for FASN-dependent lipogenesis in maintaining the metastasis-competent state. To determine whether increasing the initial abundance of metastasis-competent cells could overcome this defect, we orthotopically implanted cell populations enriched for GFP⁺ cells at a GFP^+^: GFP^-^ ratio of 2:1. GFP^+^ cell enrichment failed to rescue metastatic progression in FASN^KO^ tumors (**Extended Data Fig. 2k,l**), indicating that the reduced metastatic capacity of FASN-deficient tumors could not be overcome by increasing the initial proportion of GFP^+^ cells.

We next examined whether pharmacological FASN inhibition reproduced the effects of genetic FASN loss. We tested two inhibitors, TVB-3166, a preclinical analogue, and TVB-2640, a clinically advanced compound that has undergone clinical evaluation^7^. TVB-3166 reduced FASN protein abundance in a dose-dependent manner in both human and mouse TNBC cells (**Extended Data Fig. 2m**), whereas neither TVB-3166 nor TVB-2640 significantly affected proliferation in vitro (**Extended Data Fig. 2p,q**). In vivo, TVB-3166 treatment had little effect on primary tumor growth (**Extended Data Fig. 2r,s**) but markedly reduced lung metastatic burden (**Fig. 1e, right and i, and Extended Data Fig. 2r**), with flow cytometry independently confirming a reduction in metastasis-competent tumor cells in the lungs (**Fig. 1j and Extended Data Fig. 2t-v**). TVB-3166 was well tolerated, with no significant change in body weight or overt histopathological evidence of treatment-related toxicity (**Extended Data Fig. 2w,x**).

Together, these genetic and pharmacological studies identify FASN-dependent lipogenesis as a critical requirement for metastatic progression in TNBC, with comparatively limited effects on primary tumor growth.

### De novo lipogenesis sustains the metabolic and signaling state of metastasis-competent tumor cells

To determine how de novo lipogenesis supports metastatic competence, we examined metabolic and signaling alterations following disruption of FASN-dependent fatty acid synthesis.

Free fatty acid profiling revealed a broad reduction in intracellular free fatty acid abundance in FASN-deficient (FASN^KO^) tumors, including depletion of palmitate, stearate, and oleate, compared with FASN^WT^ tumors (**Fig. 2a and Extended Data Fig. 3a-d**). OCT4/SOX2-GFP^+^ metastasis-competent cells from FASN^WT^ tumors contained substantially higher levels of free fatty acids than GFP⁻ cells, a metabolic distinction abolished by FASN loss (**Fig. 2b and Extended Data Fig. 3e,f**). Pharmacological inhibition of FASN with TVB-3166 similarly reduced free fatty acid abundance (**Fig. 2c**). FASN^WT^ tumor cells exhibited a lower NADPH/NADP^+^ ratio than FASN^KO^ cells (**Extended Data Fig. 3g**), consistent with increased NADPH utilization for de novo fatty acid synthesis, whereas TCA-cycle metabolite pools were largely preserved following FASN loss (**Extended Data Fig. 3h**). Thus, FASN maintains a distinct free fatty acid profile in metastasis-competent tumor cells.

**Fig. 2.**
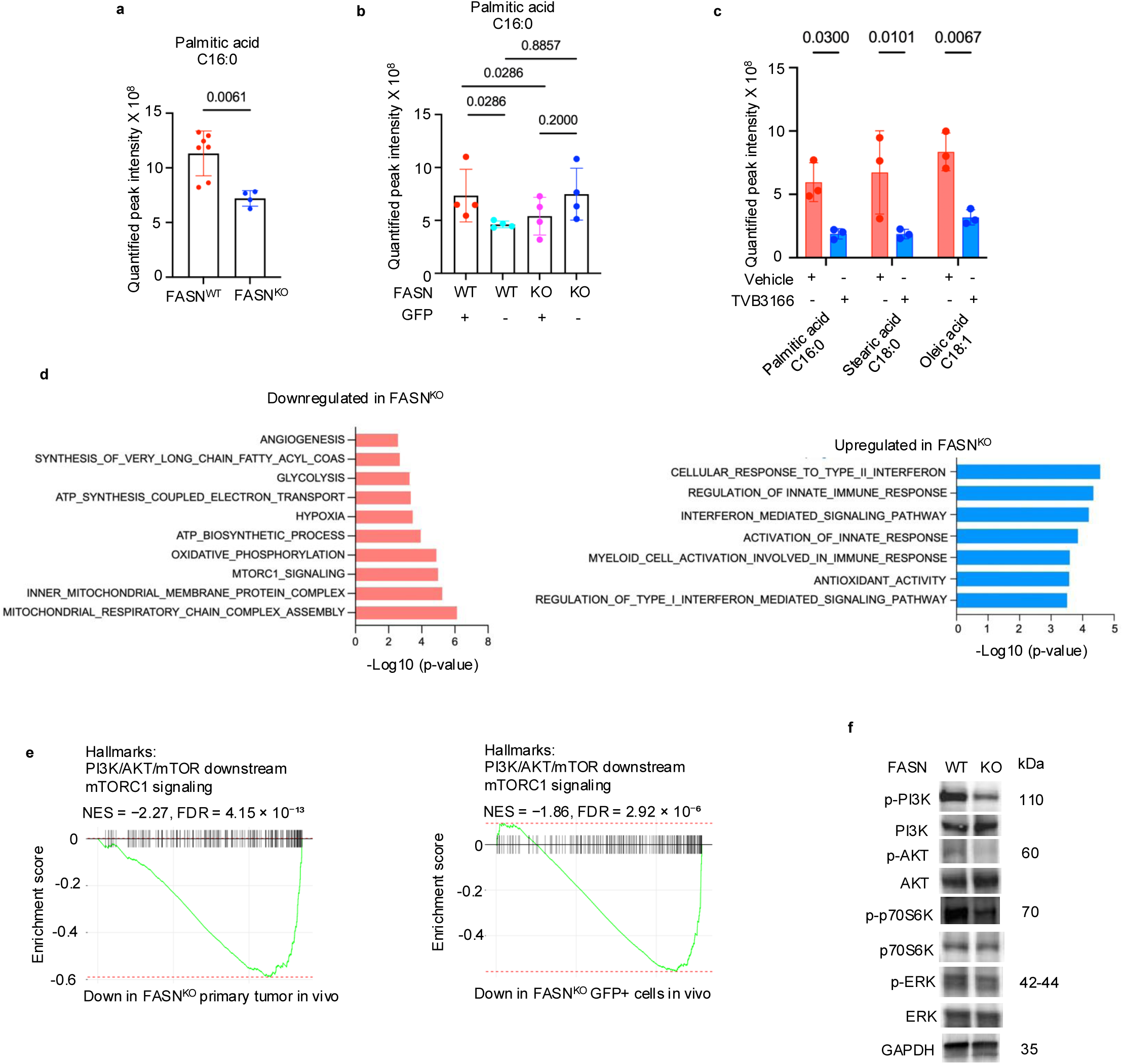
De novo lipogenesis sustains metabolic and signaling state of metastasis-competent tumor cells. a,b, LC–MS quantification of palmitic acid in FASN^WT^ and FASN^KO^ orthotopic EO771-ML1 primary tumors (**a**), and LC–MS quantification of palmitic acid in GFP⁺ and GFP⁻ cells isolated from FASN^WT^ and FASN^KO^ tumors (n = 4 biological replicates per group) (**b**). **c,** LC–MS quantification of free fatty acids (palmitic, stearic and oleic acids) in orthotopic EO771-ML1 tumors treated with vehicle or TVB-3166 (n = 3 biological replicates per group). **d**, Transcriptomic analysis of FASN^WT^ and FASN^KO^ orthotopic tumors (n = 5 biological replicates per group), showing gene set enrichment analysis (GSEA) of pathways downregulated (left) and upregulated (right) following FASN loss. Bars represent −log10(P value). NES, normalized enrichment score; FDR, false discovery rate (q-value); values are provided in the Source Data. **e,** GSEA showing reduced mTORC1 signaling following FASN loss in orthotopic EO771-ML1 primary tumors (left) and GFP^+^ tumor cells isolated from orthotopic tumors (right) (n = 5 biological replicates per group). **f,** Immunoblot analysis of PI3K–AKT–mTOR and MAPK signaling, including phosphorylated and total PI3K, AKT, p70S6K, ERK in FASN^WT^ and FASN^KO^ tumor cells isolated from orthotopic EO771-ML1 tumors. GAPDH served as a loading control. Data in a–c are presented as mean ± s.e.m. Statistical significance was assessed using a two-sided unpaired Student’s t-test for a, ordinary one-way ANOVA with multiple-comparisons tests for b, and ordinary two-way ANOVA with Šídák’s multiple-comparisons test for c. Statistical significance for pathway enrichment in d was determined using the enrichment analysis described in the Methods. GSEA enrichment statistics, including normalized enrichment scores (NES), are indicated in e. Exact P values are indicated in the figures.

We next asked whether this metabolic state was accompanied by broader changes in cellular programs associated with metastatic progression. Transcriptomic profiling of FASN^WT^ and FASN^KO^ tumors revealed reduced enrichment of transcriptional programs associated with lipid elongation, mitochondrial oxidative phosphorylation, ATP synthesis, respiratory chain assembly, and hypoxia following FASN loss (**Fig. 2d, left**). In parallel, immune-associated transcriptional programs, including T cell activation and interferon responses, showed increased enrichment (**Fig. 2d, right**). Expression of the lipogenic regulators SREBP1 and SREBP2 and the fatty acid transporter CD36 remained unchanged (**Extended Data Fig. 3i**), providing no evidence for compensatory transcriptional induction of these pathways following FASN loss^15^.

Given the coordinated metabolic changes induced by FASN loss, we next examined downstream oncogenic signaling. GSEA showed enrichment of PI3K−AKT−mTOR signaling programs, including mTORC1-associated signatures, in FASN^WT^ tumors (**Fig. 2e**). This enrichment was particularly pronounced in OCT4/SOX2-GFP^+^ metastasis-competent cells and was markedly reduced following FASN deletion, whereas these programs remained comparatively preserved in GFP^-^ cells (**Fig. 2e and Extended Data Fig. 3j**). Biochemical analysis confirmed that FASN deletion markedly attenuated PI3K−AKT−mTOR signaling without substantially affecting MAPK signaling (**Fig. 2f**), identifying PI3K−AKT−mTOR as a signaling output preferentially maintained by FASN-dependent lipogenesis. Consistent with this selectivity, MAPK pathway components showed no coordinated transcriptional response to FASN loss (**Extended Data Fig. 3k**).

Together, these findings indicate that de novo lipogenesis sustains a distinct metabolic and signaling state in metastasis-competent tumor cells, characterized by elevated free fatty acid abundance and preferential PI3K−AKT−mTOR signaling. Loss of FASN disrupts this state and is accompanied by increased expression of immune-associated transcriptional programs, prompting us to examine whether FASN-dependent lipogenesis regulates tumor immune surveillance.

### De novo lipogenesis suppresses tumor antigen presentation and CD8^+^ T-cell immune surveillance

Because FASN loss induced immune-related transcriptional programs, we next asked whether disruption of de novo lipogenesis altered the tumor immune microenvironment. Flow cytometric profiling of CD45^+^ immune cells from orthotopic tumors revealed no significant differences in the abundance of total CD8^+^ or CD4^+^ T cells between FASN^WT^ and FASN^KO^ tumors (**Extended Data Fig. 4a,b**). Instead, FASN loss enhanced CD8^+^ T-cell effector phenotypes, with increased frequencies of CD44^+^CD62L^-^ cells and increased IFNγ, TNFα, and granzyme B expression, together with elevated PD-1 expression (**Fig. 3a-d**). FASN loss also reduced polymorphonuclear myeloid-derived suppressor cells (PMN-MDSCs) (**Fig. 3e**), whereas other immune populations, including regulatory T cells, macrophages, conventional dendritic cells, and most NK-cell subsets, remained unchanged (**Extended Data Fig. 4c-l**). Pharmacological inhibition of FASN with TVB-3166 similarly increased CD8^+^ T-cell effector activity and PD-1 expression (**Fig. 3f-h**). Thus, FASN-dependent lipogenesis is associated with reduced CD8^+^

**Fig. 3.**
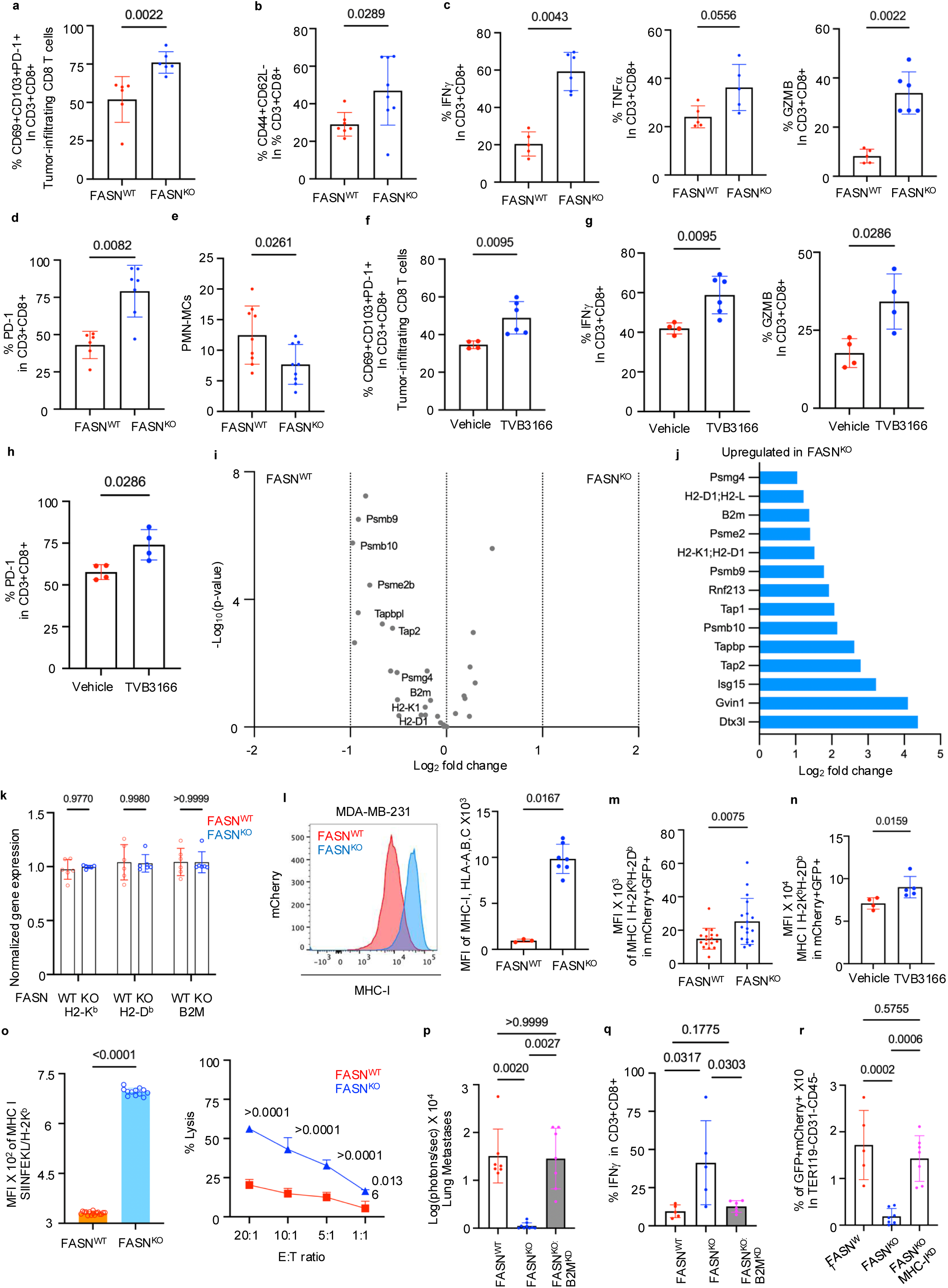
De novo lipogenesis suppresses MHC-I antigen presentation and CD8^+^ T-cell immune surveillance. a–e, Flow-cytometric profiling of immune populations in FASN^WT^ and FASN^KO^ orthotopic EO771-ML1 primary tumors (n = 6 biological replicates per group). **a**, Tumor-infiltrating CD8^+^ T cells. **b,** CD44^+^CD62L^-^ CD8^+^ T cells. **c,** IFNγ^+^, TNFα^+^ and granzyme B (GZMB)^+^ CD8^+^ T cells. **d,** PD-1^+^ CD8^+^ T cells. **e,** PMN-MDSCs. **f–h,** Immune profiling of FASN^WT^ tumors treated with vehicle or TVB-3166 (n = 4 biological replicates per group). **f,** Tumor-infiltrating CD8^+^ T cells. **g,** IFNγ^+^ and GZMB^+^ CD8^+^ T cells. **h,** PD-1^+^ CD8^+^ T cells. **i,j,** Analysis of MHC-I antigen-processing and presentation machinery in FASN^WT^ and FASN^KO^ tumors. **i,** Transcriptomic analyses of MHC-I antigen-processing and presentation genes (n = 5 biological replicates per group), **j,** Proteomic analysis showing changes in MHC-I and antigen-processing machinery protein abundance following FASN loss. **k,** RT–qPCR quantification of H2-K1, H2-D1 and B2M transcripts in GFP^+^ tumor cells isolated from primary tumors (n = 5 biological replicates per group). **l,** Flow-cytometric quantification of cell surface HLA-A, B, C on FASN^WT^ and FASN^KO^ MDA-MB-231 cells (n = 6 biological replicates per group). **m,n,** Flow cytometric quantification of cell surface MHC-I on GFP^+^ tumor cells sorted from **m**, FASN^WT^ or FASN^KO^ tumors (n = 17 FASN^WT^, 18 FASN^KO^). **n**, FASN^WT^ tumors treated with vehicle or TVB-3166 (n = 4 vehicle, 5 TVB-3166). **o,** Cell surface presentation of OVA-derived SIINFEKL–H-2K^b^ on OVA-expressing EO771-ML1 cells (left) and antigen-specific tumor-cell lysis following co-culture with OT-I CD8^+^ T cells at indicated effector-to-target (E:T) ratios (right; n = 3 biological replicates per group). **p–r,** Effects of doxycycline-inducible B2M knockdown in FASN^WT^, FASN^KO^ and FASN^KO^B2M^KD^ models following doxycycline administration. **p**, Lung metastatic burden (n = 7 FASN^WT^, 9 FASN^KO^ and 7 FASN^KO^B2M^KD^ mice). **q**, IFNγ^+^ tumor-infiltrating CD8^+^ T cells (n = 5 FASN^WT^, 5 FASN^KO^ and 6 FASN^KO^B2M^KD^ mice). **r**, Circulating mCherry^+^GFP^+^ tumor cells identified within the TER119^-^CD31^-^CD45^-^ population (n = 5 FASN^WT^, 5 FASN^KO^ and 7 FASN^KO^B2M^KD^ mice). Data in a–h, l–r are presented as mean ± s.e.m. Statistical significance was assessed using a two-sided unpaired Student’s t-test for a–h, l–n, and o (left), ordinary one-way ANOVA with multiple-comparisons tests for p–r, and ordinary two-way ANOVA with Šídák’s multiple-comparisons test for k and o (right). Differentially expressed genes and proteins are shown in I and j. Exact P values are indicated in the figures.

T-cell effector activity despite relatively limited changes in overall immune-cell composition.

To determine whether FASN loss affects the capacity of TNBC cells to colonize the lung, we analyzed metastatic outgrowth following direct vascular delivery of FASN^WT^ and FASN^KO^ tumor cells. Despite comparable tumor burden, FASN-deficient tumors exhibited increased infiltration of CD8^+^ T cells and dendritic cells (**Extended Data Fig. 4p,q**). Although FASN loss did not significantly alter metastatic burden at the examined endpoint, it was associated with increased CD8^+^ T-cell and dendritic-cell infiltration in the lungs.

Because effective CD8^+^ T-cell recognition depends on MHC-I antigen presentation, we next asked whether FASN regulates tumor-cell antigen presentation. Transcriptomic analysis revealed no significant change in the MHC-I antigen-processing and presentation gene signature^16^ following FASN loss (**Fig. 3i**). In contrast, quantitative proteomics revealed increased abundance of MHC-I complex components and multiple antigen-processing machinery proteins in FASN^KO^ tumors (**Fig. 3j**), despite unchanged transcript levels of H2-K1, H2-D1 and B2M, as confirmed by qPCR (**Fig. 3k**). Genetic or pharmacological inhibition of FASN significantly increased surface MHC-I expression in both human and murine TNBC cells in vitro (**Fig. 3l; Extended Data Fig. 2n,o and 4m,n**). OCT4/SOX2-GFP^+^ metastasis-competent cells isolated from orthotopic tumors in immunocompetent mice likewise exhibited increased surface MHC-I following FASN inhibition (**Fig. 3m,n**). Together, these findings support regulation of MHC-I antigen presentation at the protein level, without corresponding changes in the measured MHC-I transcripts.

We next asked whether these changes enhanced functional antigen presentation and antigen-specific CD8^+^ T-cell recognition. We expressed full-length ovalbumin (OVA) in EO771-ML1 cells, enabling intracellular processing and presentation of the OVA-derived SIINFEKL peptide on H-2K^b^. FASN-deficient cells displayed significantly increased surface SIINFEKL–H-2K^b^ complexes and were more efficiently killed by antigen-specific OT-I CD8^+^ T cells (**Fig. 3o**).

Thus, FASN-dependent lipogenesis suppresses peptide–MHC-I presentation and limits antigen-specific CD8^+^ T-cell cytotoxicity.

To determine whether MHC-I antigen presentation mediates the suppression of metastasis following FASN loss, we generated a doxycycline-inducible B2M knockdown model (**Extended Data Fig. 4r,s**). B2M depletion largely restored metastatic outgrowth and attenuated the enhanced CD8^+^ T-cell activation induced by FASN deficiency (**Fig. 3p,q and Extended Data Fig. 4t-x**). Furthermore, FASN loss reduced circulating tumor cells (CTCs) derived from metastasis-competent populations, an effect substantially reversed by B2M depletion (**Fig. 3r**). These findings demonstrate that tumor-cell MHC-I expression is a major contributor to the suppression of metastasis following FASN loss.

Together, these findings demonstrate that FASN-dependent lipogenesis promotes metastatic progression in part by suppressing tumor-cell MHC-I antigen presentation and limiting CD8^+^ T-cell recognition. The restoration of metastatic outgrowth following B2M depletion identifies MHC-I-dependent immune surveillance as a major contributor to the antimetastatic effects of FASN loss. We next investigated how de novo lipogenesis regulates tumor-cell antigen presentation.

### EGFR palmitoylation links de novo lipogenesis to PI3K−AKT−mTOR signaling and antigen presentation

We next sought to identify the lipid-dependent signaling mechanism linking de novo lipogenesis to tumor-cell MHC-I antigen presentation. Because FASN loss increased MHC-I protein abundance without altering MHC-I antigen-processing and presentation transcripts, we first examined alternative mechanisms that could account for this increase. Transcriptomic analysis revealed no significant changes in pathways^16–19^ associated with lysosomal function, ubiquitination, endosomal recycling, or membrane trafficking (**Extended Data Fig. 3l,m**).

Proteins associated with F-actin, which contributes to MHC-I membrane stability, were likewise unchanged (**Extended Data Fig. 3n**). Moreover, pharmacological inhibition of lysosomal function with bafilomycin A1 neither reduced the elevated surface MHC-I expression in FASN-deficient cells nor revealed differences in autophagic flux, as assessed by LC3-II accumulation (**Extended Data Figs. 3o and 4o**). These findings indicated that the increase in surface MHC-I following FASN loss was not explained by detectable changes in lysosomal degradation, membrane trafficking, or autophagic flux.

**Fig. 4.**
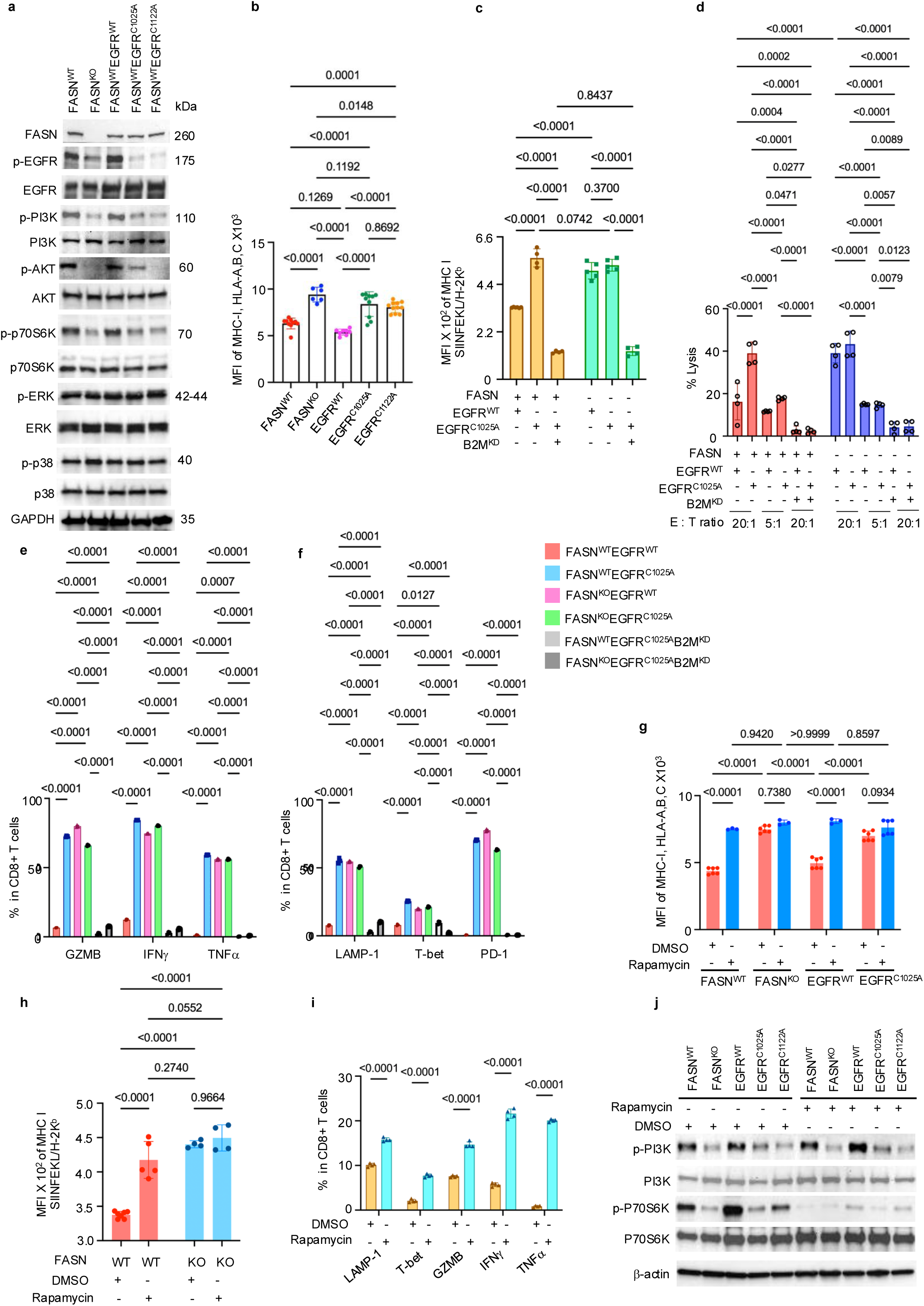
EGFR palmitoylation couples FASN-dependent lipogenesis to PI3K–AKT–mTOR signaling and immune evasion. **a,** Immunoblot analysis of EGFR, PI3K–AKT–mTOR, and MAPK signaling in MDA-MB-231-LM2 cells with the indicated FASN and EGFR genotypes, including EGFR^WT^ and the palmitoylation-deficient EGFR^C1025A^ and EGFR^C1122A^ mutants expressed in EGFR^KO^ cells with intact FASN. Phosphorylated and total signaling proteins are shown as indicated. **b,** Flow-cytometric quantification of cell-surface MHC-I in the same genotypes (n = 6 biological replicates per group). **c,d,** Functional antigen presentation in OVA-expressing EO771-ML1 cells expressing EGFRWT or palmitoylation-deficient EGFR mutants, with or without B2M knockdown. **c,** Cell-surface SIINFEKL–H-2K^b^ complexes (n = 6 EGFR^WT^, 4 EGFR^C1025A^ and 4 EGFR^C1025A^B2M^KD^ in FASN^WT^; n = 5 EGFR^WT^, 5 EGFR^C1025A^ and 4 EGFR^C1025A^B2M^KD^ in FASN^KO^). **d,** Antigen-specific OT-I CD8⁺ T-cell-mediated tumor-cell lysis at the indicated effector: target ratios (n = 4 biological replicates per group). **e,f,** Phenotypic analysis of co-cultured OT-I CD8⁺ T cells. **e,** GZMB, IFNγ and TNFα expression. **f,** LAMP-1, T-bet and PD-1 expression (n = 4 biological replicates per group). **g,** Cell-surface MHC-I on MDA-MB-231 cells with the indicated FASN or EGFR genotypes treated with DMSO or rapamycin (20 nM, 48 h; n = 5 biological replicates per group). **h,i,** Effects of rapamycin on antigen presentation and T-cell activation. **h,** Cell-surface SIINFEKL–H-2K^b^ complexes on OVA-expressing EO771-ML1 cells (n = 7 DMSO and 5 rapamycin in FASN^WT^; n = 4 DMSO and 4 rapamycin in FASN^KO^). **i,** GZMB, IFNγ, TNFα, LAMP-1 and T-bet expression in co-cultured OT-I CD8⁺ T cells (n = 4 biological replicates per group). **j,** Immunoblot analysis of mTORC1 signaling with the indicated FASN and EGFR genotypes following rapamycin treatment (20 nM, 48 h). Data in b–i are presented as mean ± s.e.m. Statistical significance was assessed using ordinary one-way ANOVA with multiple-comparisons tests for b, and ordinary two-way ANOVA with Šídák’s multiple-comparisons test for c–i. Exact P values are indicated in the figures.

We therefore sought to identify the lipid-dependent signaling mechanism connecting FASN to antigen presentation. Given the coordinated activation of FASN-dependent lipogenesis and PI3K−AKT−mTOR signaling in metastasis-competent tumor cells, we focused on signaling nodes capable of coupling lipid metabolism to this pathway. EGFR emerged as a candidate because it is an upstream regulator of PI3K−AKT−mTOR signaling and undergoes S-palmitoylation^20–22^, a reversible lipid modification that regulates receptor signaling^23^. We hypothesized that FASN-dependent lipogenesis sustains EGFR palmitoylation to maintain PI3K−AKT−mTOR signaling and suppress antigen presentation.

We quantified EGFR palmitoylation using an acyl-biotin exchange (ABE) assay^24^. Palmitoylated EGFR was readily detected in FASN^WT^ cells but was markedly reduced following FASN deletion (**Extended Data Fig. 5a, top**). Given that FASN loss selectively attenuated PI3K−AKT−mTOR signaling without substantially affecting MAPK signaling, we next asked whether EGFR palmitoylation could account for this signaling selectivity. Previous studies have shown that palmitoylated EGFR preferentially associates with the PI3K regulatory subunit PIK3R1 (p85), facilitating PI3K recruitment and activation^21^. Together with the reduction in intracellular palmitate following FASN loss, these findings implicated EGFR palmitoylation as a lipid-dependent signaling node linking de novo lipogenesis to PI3K−AKT−mTOR signaling.

EGFR undergoes S-palmitoylation at three conserved C-terminal cysteine residues, C1025, C1034, and C1122, catalyzed by the palmitoyltransferases ZDHHC20 and ZDHHC13^21,22,25^. To directly test the functional requirement for EGFR palmitoylation, we generated EGFR knockout (EGFR^KO^) TNBC cells and reconstituted them with wild-type EGFR or EGFR variants carrying cysteine-to-alanine substitutions at the identified palmitoylation sites. Wild-type and mutant EGFR were expressed at comparable protein levels, and neither FASN depletion nor disruption of EGFR palmitoylation altered total EGFR abundance or cell-surface localization (**Extended Data Fig. 5b,c**). The palmitoylation-deficient EGFR variants markedly reduced EGFR palmitoylation, phenocopying FASN depletion (**Extended Data Fig. 5a, bottom**).

Thus, EGFR palmitoylation can be disrupted without altering receptor abundance or cell-surface localization, allowing its contribution to signaling to be examined independently of receptor expression or trafficking. Importantly, disruption of EGFR palmitoylation selectively attenuated PI3K−AKT−mTOR signaling without substantially affecting MAPK activation, phenocopying FASN depletion (**Fig. 4a**). These findings establish EGFR palmitoylation as a determinant of EGFR signaling output that selectively maintains the PI3K−AKT−mTOR arm without inducing compensatory MAPK activation.

We next asked whether this selective signaling defect was sufficient to restore antigen presentation. Expression of palmitoylation-deficient EGFR increased cell-surface MHC-I expression without altering MHC-I transcript levels (**Fig. 4b and Extended Data Fig. 5i**), phenocopying FASN deficiency and further supporting transcription-independent regulation of MHC-I abundance.

To determine whether increased MHC-I translated into functional antigen presentation, we expressed full-length ovalbumin (OVA) in cells expressing palmitoylation-deficient EGFR. These cells exhibited increased surface presentation of OVA-derived SIINFEKL–H-2K^b^ complexes (**Fig. 4c**) and enhanced killing by antigen-specific OT-I CD8^+^ T cells (**Fig. 4d**). This enhanced antigen-specific recognition was accompanied by increased frequencies of effector CD8^+^ T cells expressing T-bet, LAMP-1, granzyme B, IFNγ, and TNFα, together with increased PD-1 expression (**Fig. 4e,f**), consistent with enhanced antigen-driven T-cell activation.

Importantly, B2M depletion abolished SIINFEKL–H-2K^b^ surface expression (**Fig. 4c**) and the enhanced antigen-specific CD8^+^ T-cell responses induced by disruption of EGFR palmitoylation, including tumor-cell killing, cytokine production, and effector differentiation (**Fig. 4d–f**). These findings establish tumor-cell MHC-I antigen presentation as a functional downstream effector of EGFR palmitoylation-dependent immune regulation.

Having established EGFR palmitoylation as the lipid-dependent signaling node, we next asked whether mTOR functions as the downstream effector linking EGFR signaling to antigen presentation. PI3K−AKT−mTOR signaling has been implicated in MHC-I regulation through transcriptional control, receptor ubiquitination, lysosomal degradation, and membrane trafficking^26–28^, but our findings indicated that FASN-dependent regulation of MHC-I was independent of transcription and not explained by altered lysosomal degradation or autophagic flux. We therefore directly tested the contribution of mTOR signaling to tumor-cell antigen presentation. Pharmacological inhibition of mTOR with Torin1, a catalytic inhibitor of mTORC1 and mTORC2, suppressed phosphorylation of p70 S6 kinase (p70 S6K) and significantly increased cell-surface MHC-I expression (**Extended Data Fig. 6a,b**). Rapamycin similarly restored cell-surface MHC-I expression (**Fig. 4g**) and enhanced functional presentation of full-length OVA, increasing surface SIINFEKL−H-2K^b^ complexes and antigen-specific CD8^+^ T-cell activation (**Fig. 4h,i**), while suppressing downstream mTOR signaling (**Fig. 4j**). Thus, pharmacological disruption of mTOR signaling phenocopied the effects of FASN depletion and palmitoylation-deficient EGFR. mTOR inhibition did not alter MHC-I transcript abundance (**Extended Data Fig. 6c–f**), further supporting transcription-independent regulation of tumor antigen presentation by mTOR signaling. These findings identify mTOR as a downstream effector through which the FASN−EGFR palmitoylation pathway controls tumor-cell antigen presentation.

We next asked whether extracellular lipid availability could compensate for loss of de novo lipogenesis. FASN-deficient cells remained depleted of intracellular palmitate and total free fatty acids despite culture in regular FBS (**Extended Data Figs. 5g,h and 6g**). Reduced EGFR signaling and proliferation likewise persisted across regular and lipid-depleted conditions (**Extended Data Fig. 6h-l**). Thus, regular serum did not compensate for FASN loss, supporting a specific requirement for de novo lipogenesis in maintaining EGFR signaling.

Together, these findings establish a tumor-intrinsic immunometabolic circuit in which FASN-dependent lipogenesis sustains EGFR palmitoylation and selective PI3K−AKT−mTOR signaling, thereby suppressing MHC-I antigen presentation and antigen-specific CD8⁺ T-cell recognition. Disruption of this axis at the level of FASN, EGFR palmitoylation, or mTOR restores tumor-cell immune visibility, defining a lipid-dependent signaling mechanism that links tumor metabolic state to antigen presentation during metastatic progression.

### A lipogenic−PI3K−AKT−mTOR state is associated with reduced immune surveillance in human TNBC and basal-like breast cancer

To examine the clinical relevance of the lipogenesis−PI3K−AKT−mTOR relationship identified experimentally, we analyzed RNA-seq profiles from patient-derived TNBC organoids and normal breast organoids. FASN and mTOR expression were significantly higher in tumor-derived than normal-derived organoids. FASN expression positively correlated with RPTOR, a component of mTORC1, in TNBC organoids, whereas this association was not significant in normal organoids (**Fig. 5a**). In contrast, FASN expression showed no significant correlation with MAPK1, MAPK3, or MAPK14 in either group (**Extended Data Fig. 7a,b**), supporting an association between FASN expression and an mTORC1 component in human TNBC organoids.

**Figure 5.**
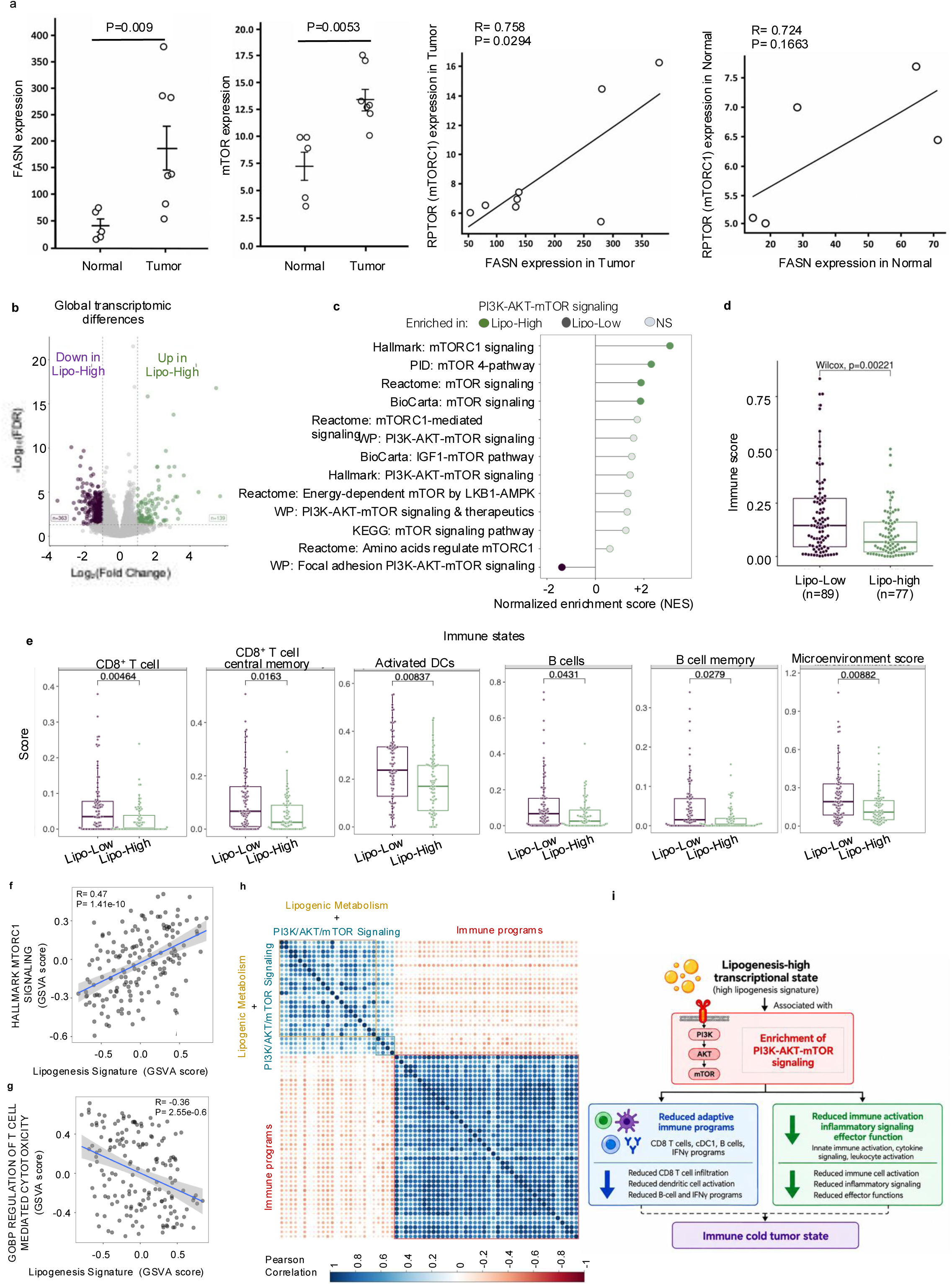
Elevated lipogenesis is associated with enrichment of PI3K–AKT–mTOR transcriptional programs and reduced immune-response signatures in human basal-like breast cancer. **a,** RNA-seq analysis of patient-derived TNBC and normal breast organoids showing correlations between FASN and RPTOR expression. FASN expression positively correlated with RPTOR expression in tumor-derived organoids (Pearson r = 0.758, P = 0.0294; n = 8), whereas the correlation was not statistically significant in normal-derived organoids (r = 0.724, P = 0.1663; n = 5). **b,** Differential gene-expression analysis comparing lipogenesis-high (Lipo-high) and lipogenesis-low (Lipo-low) basal-like breast tumors. Differentially expressed genes were defined by a false discovery rate (FDR) < 0.05 and absolute log_2_ fold change > 1. **c,** Gene set enrichment analysis (GSEA) showing enrichment of PI3K–AKT–mTOR-gene set collections, including Hallmark, Reactome, PID, BioCarta and WikiPathways gene sets. Normalized enrichment scores (NES) are shown for representative gene sets. Filled circles indicate significant enrichment (Benjamini–Hochberg-adjusted P < 0.05); whereas open circles indicate nonsignificant enrichment (NS). **d,** Comparison of xCell immune scores between Lipo-low and Lipo-high tumors. **e,** Comparison of inferred immune-cell abundance and immune-microenvironment scores using TIMER and xCell in Lipo-low and Lipo-high tumors. Group comparisons were performed using two-sided Wilcoxon rank-sum tests with Benjamini–Hochberg correction for multiple testing. **f,g,** Pearson correlations between the continuous lipogenesis GSVA score and the GSVA scores for Hallmark mTORC1 signaling (f) and T-cell cytotoxicity (g). Pearson correlation coefficients (r) and P values are indicated. **h,** Pairwise Pearson correlation matrix of GSVA scores for lipogenesis, PI3K–AKT–mTOR signaling and immune-surveillance programs. Significant correlations were defined by Benjamini–Hochberg-adjusted P < 0.05. **i,** Proposed model illustrating the association of elevated lipogenesis-related gene expression with enrichment of PI3K–AKT–mTOR transcriptional programs and reduced immune-response signatures in human basal-like breast cancer. Correlations in a, f and g were assessed using Pearson correlation. Group comparisons in d and e were performed using two-sided Wilcoxon rank-sum tests, with Benjamini–Hochberg adjustment for multiple comparisons where applicable. Adjusted P values for the pathway-enrichment and immune-cell comparisons are provided in the Supplementary Data Tables 1-5. Group comparisons in a, d, and e were performed using two-sided Wilcoxon rank-sum tests, with Benjamini–Hochberg adjustment for multiple comparisons where applicable. Correlations in a, f, and g were assessed using Pearson correlation. Pathway associations in h were considered significant at a Benjamini–Hochberg-adjusted P < 0.05. Exact P values are shown in the figure.

We next asked whether this relationship extended across human basal-like breast cancer. We analyzed transcriptomic profiles from 166 primary basal-like breast tumors in The Cancer Genome Atlas (TCGA) and stratified tumors according to a focused lipogenesis score comprising ACLY, ACACA, FASN, SCD, ELOVL6, GPAM, DGAT1, and DGAT2, representing key steps in de novo fatty acid synthesis, fatty acid elongation and desaturation, and triglyceride synthesis^29^. Lipogenesis-high tumors exhibited broad transcriptional remodeling relative to lipogenesis-low tumors (**Fig. 5b**). Gene set enrichment analysis of receptor tyrosine kinase (RTK) downstream signaling programs revealed consistent enrichment of PI3K−AKT−mTOR-associated pathways, including mTORC1, mTOR, and PI3K−AKT signatures (**Fig. 5c**). Other major RTK downstream signaling programs showed substantially less enrichment (**Extended Data Fig. 7c**). Thus, across human basal-like breast cancers, elevated lipogenesis-associated gene expression was preferentially associated with a PI3K−AKT−mTOR transcriptional state.

We next examined whether this lipogenic–signaling state was associated with the immune landscape. Lipogenesis-high tumors exhibited significantly lower immune and microenvironment scores (**Fig. 5d**), accompanied by reduced inferred CD8⁺ T-cell, activated dendritic-cell, and memory-lymphocyte abundance (**Fig. 5e**). In contrast, MHC-I signature scores were not significantly different between lipogenesis-high and lipogenesis-low tumors (**Extended Data Fig. 7d**), a finding compatible with the absence of major changes in MHC-I transcript expression observed experimentally. Independent immune-deconvolution approaches similarly identified broad differences in inferred immune-cell composition between lipogenesis-high and lipogenesis-low tumors (**Extended Data Fig. 8**). Importantly, these relationships were also evident across the continuous range of lipogenesis scores.

Increasing lipogenesis scores were positively associated with PI3K−AKT−mTOR pathway signatures and inversely associated with CD8^+^ T-cell cytotoxicity signatures (**Fig. 5f,g**), indicating that the relationship between tumor lipogenic state and immune-associated transcriptional programs was not dependent on categorical stratification.

To examine the relationship among these transcriptional programs, we performed GSVA-based pairwise correlation analysis followed by unsupervised hierarchical clustering. This analysis identified two inversely correlated groups of transcriptional signatures: a lipogenic– signaling module comprising lipid metabolic and PI3K−AKT−mTOR programs, and an immune-surveillance module comprising immune-response signatures (**Fig. 5h**). The inverse relationship between these modules provides transcriptomic correlate of the inverse relationship between lipogenic signaling and immune-associated programs observed experimentally. The complete pathway architecture and hierarchical relationships among the analyzed signatures are shown in Extended Data Fig. 9.

We next tested whether this relationship was reproducible in an independent breast cancer cohort. Analysis of basal-like tumors from METABRIC demonstrated positive correlations between lipogenesis scores and multiple independently defined PI3K−AKT−mTOR-related gene sets, including PI3K/AKT/mTOR signaling, mTOR signaling, and TOR/rapamycin-target pathways (**Extended Data Fig. 10a-c**). Lipogenesis-high tumors also exhibited reduced immune-response programs together with increased lipid and sterol metabolic programs (**Extended Data Fig. 10d**), and lipogenesis scores were inversely associated with immune– cytokine-response signatures (**Extended Data Fig. 10e**). Thus, an independent human cohort confirmed the association of elevated lipogenesis scores with PI3K−AKT−mTOR-associated transcriptional programs and reduced immune response signatures (**Fig. 5i**). Together, these analyses identify a reproducible association between lipogenesis-related gene expression, PI3K−AKT−mTOR transcriptional programs, and reduced immune-response signatures in human basal-like breast cancer. These human associations are consistent with the FASN-dependent signaling and immune-evasion mechanism identified experimentally and support its relevance to human TNBC.

## Discussion

Despite advances in immune checkpoint blockade, TNBC remains difficult to treat with immunotherapy, in part because tumor-intrinsic mechanisms restrict antigen presentation and enable escape from CD8^+^ T-cell surveillance^3^. Here, we identify de novo lipogenesis as a regulator of tumor immune visibility and metastatic competence. FASN-dependent lipogenesis sustains EGFR palmitoylation, thereby maintaining PI3K−AKT−mTOR signaling and limiting MHC-I antigen presentation through changes in antigen-processing machinery protein abundance without corresponding changes in the measured MHC-I transcripts. Disruption of this metabolic–signaling axis restores tumor-cell peptide–MHC-I presentation and enhances antigen-specific CD8^+^ T-cell recognition. These findings establish a mechanistic link between tumor lipid metabolism, receptor lipidation, oncogenic signaling, and immune surveillance during metastatic progression.

Previous work established de novo fatty acid synthesis as a metabolic requirement for breast cancer brain metastasis, where tumor cells adapt to limited extracellular lipid availability^6^.

Our findings extend this concept by demonstrating that de novo lipogenesis also regulates metastatic progression through tumor immune visibility. Although FASN loss had limited effects on primary tumor growth, it markedly suppressed spontaneous metastasis. Importantly, direct vascular delivery of FASN-deficient cells resulted in comparable lung tumor burden at the examined endpoint, indicating that FASN loss did not significantly reduce metastatic burden in this experimental setting. Nevertheless, FASN-deficient metastatic tumors exhibited increased infiltration of CD8^+^ T cells and dendritic cells, suggesting altered immune interactions within the metastatic lung microenvironment.

The substantial restoration of metastatic outgrowth and circulating tumor cells following B2M depletion supports tumor-cell MHC-I expression as a major mediator of the antimetastatic effects of FASN loss. Together, these findings suggest that FASN-dependent lipogenesis supports metastatic progression, at least in part, by limiting tumor-cell recognition by CD8^+^ T cells.

A central mechanistic feature of this pathway is the selective requirement for EGFR palmitoylation in maintaining PI3K−AKT−mTOR signaling. Disruption of EGFR palmitoylation attenuated PI3K−AKT−mTOR signaling without substantially affecting MAPK signaling, EGFR abundance, or cell-surface localization. EGFR lipidation therefore appears to regulate signaling output rather than simply receptor expression or membrane localization. Although EGFR palmitoylation has previously been implicated in receptor signaling, our findings identify FASN-dependent lipogenesis as an upstream regulator of this modification and demonstrate that disrupting this pathway restores MHC-I antigen presentation and antigen-specific CD8^+^ T-cell recognition, linking receptor lipidation to metastatic immune escape.

An important question raised by these findings is how de novo lipogenesis supports EGFR palmitoylation. FASN loss markedly depleted intracellular palmitate and reduced EGFR palmitoylation, consistent with a model in which de novo lipogenesis maintains a lipid pool that supports receptor S-palmitoylation. FASN-deficient cells remained depleted of intracellular palmitate despite culture in complete medium containing FBS, suggesting that lipids available under standard culture conditions do not compensate for FASN loss in maintaining intracellular palmitate abundance. This raises the possibility that lipid source, compartmentalization, or intracellular trafficking influences substrate availability for signaling-associated lipidation reactions. However, because we did not directly trace newly synthesized palmitate into palmitoylated EGFR, whether newly synthesized palmitate directly supplies the substrate for EGFR palmitoylation or instead maintains a functionally distinct intracellular lipid pool remains unresolved. Defining how cellular lipid flux is coupled to receptor lipidation will be an important direction for future investigation.

A corresponding lipogenic–signaling state was evident in human breast cancer. In patient-derived TNBC organoids, FASN expression was associated with components of the mTORC1 pathway, whereas correlations with the examined MAPK pathway genes were not statistically significant. Across independent TCGA and METABRIC cohorts, elevated lipogenesis-associated gene expression correlated with PI3K−AKT−mTOR transcriptional programs and reduced immune-response signatures. Although these human datasets do not establish EGFR palmitoylation or post-transcriptional regulation of MHC-I, they demonstrate that the relationship between lipogenesis-associated transcriptional programs, PI3K−AKT−mTOR pathway signatures, and reduced immune-response signatures is reproducible in human basal-like breast cancer. These findings support the clinical relevance of the immunometabolic relationship identified experimentally.

An important unresolved question is how PI3K−AKT−mTOR signaling controls antigen-processing machinery abundance. FASN loss increased MHC-I and multiple antigen-processing machinery proteins without corresponding changes in the measured MHC-I transcripts, and our findings did not support altered autophagic flux as the primary explanation. These observations suggest post-transcriptional regulation of antigen-processing machinery abundance, potentially involving altered protein synthesis or degradation, while changes in trafficking may additionally contribute to increased surface MHC-I expression. Defining how mTOR signaling controls antigen-processing machinery proteostasis will be important for understanding how oncogenic signaling pathways regulate tumor antigen presentation. Previous work in hepatocellular carcinoma linked FASN inhibition to reduced MHC-I palmitoylation and lysosomal degradation^17^. Our findings reveal a distinct mechanism in which de novo lipogenesis regulates antigen presentation indirectly through EGFR palmitoylation and selective PI3K−AKT−mTOR signaling, highlighting multiple routes through which lipid metabolism can control tumor antigen presentation.

The pharmacological tractability of FASN and mTOR raises the possibility of targeting this metabolic–signaling circuit to enhance tumor immune visibility. Our findings provide a rationale for evaluating FASN inhibition in combination with immune checkpoint blockade, particularly in tumors with elevated lipogenesis-associated gene expression and reduced antigen presentation. However, whether FASN inhibition improves checkpoint blockade efficacy in vivo, and how its systemic metabolic and immune effects influence therapeutic responses, remain to be determined.

Collectively, our findings identify de novo lipogenesis as a regulator of metastatic immune escape in TNBC. Beyond its biosynthetic role, FASN-dependent lipogenesis sustains EGFR palmitoylation and PI3K−AKT−mTOR signaling to suppress tumor-cell MHC-I antigen presentation and CD8^+^ T-cell recognition. This mechanism links tumor lipid metabolism to immune surveillance and metastatic progression, highlighting how metabolic adaptation can promote tumor immune escape.

## Methods

### Cell lines and culture conditions

Human MDA-MB-231-LM2 and murine EO771-ML1 breast cancer cells expressing mCherry and firefly luciferase were cultured in complete medium containing DMEM supplemented with 10% FBS, 1% L-glutamine, 1% penicillin–streptomycin (PS) and 2 mM HEPES. Cells expressing the OCT4/SOX2-GFP reporter system were used to identify and isolate metastasis-competent GFP^+^ and metastasis-deficient GFP^-^ tumor-cell populations by flow cytometry, as described previously^11–13^. All cell lines were periodically tested for mycoplasma. A list of reagents is included in **Supplementary Table S4**.

### CRISPR/Cas9-mediated gene knockout

FASN knockout in EO771-ML1 cells was generated using CRISPR/Cas9 ribonucleoprotein (RNP) electroporation (Neon Transfection System, Thermo Fisher Scientific) following the manufacturer’s instructions^30^. The crRNA and tracrRNA were obtained from Integrated DNA Technologies (IDT). Briefly, sgRNA duplexes were formed by annealing ATTO-647–tracrRNA with target-specific crRNAs, followed by assembly with Cas9 protein at a 1:1.2 molar ratio.

Cells (1 × 10^6^) were resuspended in Neon Buffer R, mixed with RNP complexes and electroporation enhancer, and electroporated (1,400 V, 10 ms, 4 pulses). Cells were recovered in complete medium, and knockout efficiency was assessed 18 h post-transfection. Single cells were sorted by flow cytometry, clonally expanded, and validated by Western blot. For stable FASN or EGFR knockout in MDA-MB-231-LM2 cells, gene-targeting gRNAs were cloned into the Lenti-Cas9-gRNA-TagBFP2 vector (Addgene #124774). Lentiviral particles were produced in HEK293 cells and used to transduce target cells. Knockout clones were isolated by flow cytometry or validated by Western blot. CRISPR guides, reagents, and antibodies are listed in **Supplementary Table S3, S4 and S6.**

### Generation and expression of palmitoylation-deficient EGFR constructs

Palmitoylation-deficient human and mouse EGFR mutants (C1025A, C1122A) were generated from wild-type constructs using the Q5 Site-Directed Mutagenesis Kit (New England Biolabs). Lentiviral particles were produced in HEK293T cells and used to transduce EGFR knockout cells, followed by antibiotic selection. Expression of wild-type and mutant EGFR was confirmed by Western blotting, and cell-surface EGFR abundance was assessed by flow cytometry. DNA cloning reagents are listed in **Supplementary Table S7**.

### Generation of cells with inducible B2M knock-down

For doxycycline-inducible B2M knockdown, B2M shRNA was cloned into the pLV[miR30]-TRE3G>EBFP-miR30-shRNA vector (VectorBuilder) using XbaI/SpeI. FASN^WT^ or FASN^KO^ EO771-ML1 cells expressing EGFR^WT^ or the indicated EGFR mutants were transduced and selected with 10 μg/ml blasticidin in the presence of 1 μg/ml doxycycline. Cells were enriched by FACS based on EBFP expression. B2M knockdown was confirmed by immunoblotting and by reduced cell-surface MHC-I expression. The B2M shRNA and control shRNA sequences are provided in **Supplementary Table S2**, and DNA cloning and molecular biology reagents are listed in **Supplementary Table S7**.

### Cell proliferation assays

Cell proliferation was measured using the CellTiter 96 AQueous One Solution MTS assay (Promega, G3582). Cells were seeded at 5,000 cells per well and cultured under the indicated conditions. Absorbance at 490 nm was measured using a SpectraMax M5 multimode microplate reader (Molecular Devices). Lipid-dependent proliferation was assessed after 72 h. For pharmacological studies, cells were treated with TVB-3166 or TVB-2640 (10 μM) or rapamycin (20 nM) for 72 h. DMSO was used as the vehicle control. Experiments included at least three biological replicates. Reagents used in these experiments are listed in **Supplementary Table S4**.

### Signaling assays

For signaling studies, cells were treated with TVB-3166 or TVB-2640 (10 μM), rapamycin (20 nM), or Torin1 (100 nM) for 48 h. Parallel cultures were analyzed by immunoblotting.

Experiments included at least three biological replicates. Reagents used in these experiments are listed in **Supplementary Table S4**.

### Lipid depletion experiments

FASN^WT^ and FASN^KO^ cells were cultured in complete medium containing either regular FBS or lipid-depleted FBS for 72 h. Cells were subsequently collected for free fatty acid profiling, immunoblotting and proliferation assays, as indicated. For free fatty acid profiling, 1 × 10^6^ cells were collected for intracellular free fatty acid analysis. Intracellular free fatty acid abundance was normalized to cell number. Reagents used in these experiments are listed in **Supplementary Table S4**.

### In vitro invasion assay

Cell invasion was assessed using Matrigel-coated invasion inserts (BD BioCoat) with 8 μm pore size. Cells (2.5 × 10^4^ per insert) were seeded in serum-free DMEM, with DMEM containing 10% FBS in the lower chamber as the chemoattractant. After 24 h, non-invasive cells were removed from the upper surface of the inserts. Invaded cells were fixed and stained with Kwik-Diff (Thermo Fisher Scientific), images in six randomly selected fields per insert and quantified by phase-contrast microscopy. Experiments included at least three independent biological replicates. Reagents are listed in **Supplementary Table S4**.

### Cell cycle analysis

Cells were fixed in ice-cold 70% ethanol, washed, and stained with propidium iodide (PI) and RNase A for 30 min at room temperature in the dark (Abcam kit). DNA content was analyzed by flow cytometry, and G0/G1, S, and G2/M phases were quantified based on PI fluorescence intensity. Reagents are listed in **Supplementary Table S4**.

### 3D-bioprinting and spheroid growth analysis

EO771-ML1 FASN^WT^ and FASN^KO^ cells were seeded in AggreWell 800 plates at 7.5 × 10^5^ cells per well to generate spheroids^31^. Mature spheroids were resuspended in 10% methacrylated gelatin (95% degree of methacrylation) at 3 ml per well and loaded into amber cartridges compatible with a Cellink BioX 3D bioprinter. Spheroid-containing gels were distributed into 24-well plates using a droplet-bioprinting protocol and cultured under the indicated conditions. Bright-field images acquired on days 0, 6 and 10 were analyzed using CellProfiler v4.2.7. Images were converted to grayscale, smoothed using a Gaussian filter (σ = 2) and segmented by Sobel edge detection to define spheroid boundaries. Spheroid area and mean image intensity were quantified for individual spheroids. Opacity-adjusted area (OAA) was calculated as spheroid area × (1−mean intensity)^2^/ minimum spheroid area in the experimental group, where the minimum spheroid area was 23,000 square pixels. OAA values are reported in arbitrary units and were normalized to the respective control group. Four independent biological replicates were analyzed per condition.

### Animal experiments

All animal procedures were performed in compliance with protocols approved by the Institutional Animal Care and Use Committee (IACUC) at Weill Cornell Medical College (WCM protocol no. 0806-763A), adhering to established ethical guidelines. Female C57BL/6J mice (The Jackson Laboratory) were used unless otherwise specified. OT-I (C57BL/6-Tg (TcraTcrb)1100Mjb/J) and Thy1.1 mice (B6.PL-Thy1a/CyJ) were obtained from The Jackson Laboratory and bred in-house. Mice were housed under standard conditions (12 h light/dark cycle, 65-75°F, 40–60% humidity) and used for in vivo experiments at 8 weeks of age.

Reagents and laboratory supplies used in these experiments are listed in **Supplementary Table S4**.

### Orthotopic tumor model

Female C57BL/6J mice (n = 10/group) were orthotopically injected with 1 × 10^5^ FASN^WT^ or FASN^KO^ EO771-ML1 cells in 50 μl HBSS into the fourth mammary fat pad. In pharmacologic studies, mice received FASN^WT^ EO771-ML1 cells. Primary tumors were allowed to grow for approximately 4 weeks. During the treatment period, tumor growth was monitored by bioluminescence imaging (BLI) using a Xenogen imaging system and by caliper measurements. For BLI, mice were administered D-luciferin (150 mg/kg, intraperitoneally) and imaged 10 min after injection using an IVIS imaging system. Bioluminescence was quantified using Living Image software by measuring total photon flux within defined regions of interest. When primary tumors reached 1 cm^3^, they were surgically resected and mice were then monitored weekly for lung metastasis by BLI. Around 3 weeks post-surgical resection, mice were euthanized and lungs were collected for histological and downstream analyses. Survival cohorts were monitored until predefined humane endpoints.

### Orthotopic model with inducible B2M knockdown

Mice (n = 10 per group) were orthotopically injected with 1 × 10^5^ EO771-ML1 cells expressing FASN^WT^, FASN^KO^ or FASN^KO^/B2M^KD^ into the fourth mammary fat pad. Doxycycline-containing chow (200 mg/kg doxycycline; Envigo) was provided ad libitum starting on day 5 after tumor implantation and maintained throughout the experiment. Tumors were allowed to grow to approximately 1 cm^3^ and were then surgically resected. Mice were followed longitudinally by BLI for lung metastasis and survival. Reagents and laboratory supplies used in these experiments are listed in **Supplementary Table S4**.

### In vivo FASN inhibition

Mice bearing orthotopic FASN^WT^ EO771-ML1 tumors received daily intraperitoneal injections of TVB-3166 (5 mg/kg) starting on day 5 after tumor implantation. TVB-3166 was formulated in 10% DMSO, 40% PEG300, 5% Tween-80 and 45% saline, as recommended for in vivo formulation. Primary tumor growth and lung metastasis were monitored by BLI and caliper measurements. Toxicity was assessed by body weight, clinical status and H&E histological analysis. Treatment was continued for 25 days. Pharmacological inhibitors are listed in **Supplementary Table S4**.

### Histology and quantification of lung metastases

Lungs were fixed in 4% paraformaldehyde for at least 48 h, processed through ethanol and xylene, and embedded in paraffin using a Leica ASP6025 tissue processor (Leica Biosystems, Deer Park, IL, USA). Paraffin blocks were sectioned at 5 μm and stained with hematoxylin and eosin (H&E). Sections were evaluated by a board-certified veterinary pathologist (SEC), who documented microscopic changes in tumors and adjacent pulmonary tissues. Pulmonary tumor burden was quantified for each mouse by counting large expansile tumor masses, perivascular/peribronchial and alveolar neoplastic aggregates, and subpleural neoplastic aggregates. The composite tumor burden was calculated as the sum of the numbers of large tumors, perivascular/peribronchial and alveolar aggregates, and subpleural aggregates.

Intravascular tumor rafts and individual neoplastic cells were also recorded separately but were not included in the composite tumor-burden calculation. H&E-stained sections were scanned using an Olympus VS200 slide scanner (Evident Scientific, Hamburg, Germany), and images were analyzed using OlyVIA software v3.4.1 (Olympus).

### Immunological assays

#### i. OT-I CD8⁺ T cell cytotoxicity assay

In vitro BLI-based cytotoxicity assays were performed as described previously^32,33^. The full-length OVA expression plasmid was obtained from Addgene (plasmid #135073). Stable OVA-expressing EO771-ML1 cells were generated by lentiviral transduction followed by puromycin selection. OVA expression was confirmed by flow cytometry using parental, non-transduced EO771-ML1 cells as a negative control. Surface presentation of the OVA-derived SIINFEKL peptide in complex with H-2K^b^ was assessed by flow cytometry using an antibody specific for the SIINFEKL–H-2K^b^ complex. Live, single tumor cells were identified as DAPI-negative and mCherry-positive using standard viability and singlet gating. Surface SIINFEKL–H-2K^b^ presentation was quantified by mean fluorescence intensity (MFI). Mouse splenic OT-I CD8^+^ T cells were isolated by magnetic-activated cell sorting (MACS) using CD8α MicroBeads. T cells were activated with anti-CD3ε and anti-CD28 antibodies (2 μg/ml each) in complete T-cell medium consisting of RPMI-1640 supplemented with 10% FBS, 1% PS, 2 mM L-glutamine, 1 mM sodium pyruvate and 50 μM β-mercaptoethanol, with IL-2 (50 U/ml). Luciferase-expressing EO771-ML1-OVA cells were co-cultured with activated OT-I CD8^+^ T cells for 4 h at the indicated effector-to-target (E:T) ratios. Tumor-cell viability was quantified by BLI, and percentage tumor-cell killing relative to tumor-only controls was calculated as described previously^32,33^. T-cell activation and cytokine production were assessed by flow cytometry.

Reagents and antibodies used in these experiments are listed in **Supplementary Tables S4 and S5**.

#### ii. Flow cytometry analysis

Cells were stained in FACS buffer and analyzed by flow cytometry. For in vitro assays, dead cells were excluded using DAPI. For in vivo analyses, tumors were dissociated using collagenase IV (1 mg/ml), hyaluronidase (50 U/ml) and DNase I (0.1 mg/ml), followed by TrypLE digestion and ACK lysis. Single-cell suspensions were stained with surface markers in the presence of Fc receptor blocking reagent. Tumor cells were identified as CD45^-^ mCherry^+^GFP^+^ or CD45^-^mCherry^+^GFP^-^ cells, whereas immune cells were identified as CD45^+^ cells. Cell-surface MHC-I was measured on viable, non-permeabilized cells using antibodies against mouse H-2K^b^/H-2D^b^ or human HLA-ABC as appropriate. Surface EGFR was measured similarly using an EGFR-specific antibody. Surface fluorescence was quantified as median fluorescence intensity (MFI).

For immune profiling, CD45^+^ cells were stimulated with PMA and ionomycin in the presence of brefeldin A and monensin for 6 h, followed by surface staining, fixation and permeabilization, and intracellular staining. Viable cells were identified using Zombie Aqua. Major immune populations were identified by sequential gating of singlets, viable cells and CD45^+^ cells, followed by lineage-specific markers. Detailed gating strategies are provided in Supplementary Information. Fluorescence-minus-one (FMO) controls were used to define positive populations where appropriate. CTCs were identified in peripheral blood following ACK lysis as DAPI^-^ CD45^-^CD31^-^Ter119^-^mCherry^+^GFP^+^ cells.

shRNAs, antibodies and other reagents used in these experiments are listed in **Supplementary Tables S2, S4 and S5**. Flow cytometry was performed using a BD FACSymphony S6 and a BD LSRFortessa at the Weill Cornell Medicine CLC Flow Cytometry Core Facility. Compensation was performed using single-stained compensation beads, and data were analyzed using FlowJo v10 and v11.

### Biochemical analyses

#### i. Acyl-biotin exchange assay for EGFR palmitoylation

Protein S-palmitoylation was assessed by acyl-biotin exchange (ABE) as described previously^22^. Cells were lysed in buffer containing 20 mM N-ethylmaleimide (NEM) and incubated overnight at 4 °C with gentle rotation to block free cysteine residues. Proteins were precipitated by chloroform–methanol precipitation, resuspended in 4% SDS buffer, and divided equally into hydroxylamine-negative (HAM−) and hydroxylamine-positive (HAM+) samples.

HAM+ samples were treated with 0.7 M hydroxylamine for 1 h at room temperature to cleave thioester-linked S-acyl groups, whereas HAM− samples were treated identically without hydroxylamine. Following re-precipitation, newly exposed cysteine residues were labeled with 1 mM biotin-HPDP for 1 h at room temperature. Biotinylated proteins were enriched using streptavidin–agarose beads, washed extensively and eluted in Laemmli sample buffer.

Samples were analyzed by immunoblotting for EGFR, with hydroxylamine-dependent biotinylation used as the measure of EGFR palmitoylation. Reagents used in these experiments are listed in **Supplementary Table S4**.

#### ii. NADPH/NADP^+^ measurements

The NADPH/NADP^+^ ratio was measured using a NADPH/NADP^+^ Fluorometric Assay Kit (Abcam) according to the manufacturer’s instructions. Equal numbers of FASN^WT^ and FASN^KO^ EO771-ML1 tumor cells isolated from primary tumors were lysed and analyzed in biological triplicates. Fluorescence was measured using a SpectraMax M5 multimode microplate reader (Molecular Devices) at 560 nm excitation and 590 nm emission. Reagents used in these experiments are listed in **Supplementary Tables S4**.

### Western blot analyses

Cell lysates were prepared in RIPA buffer (Millipore) and protein concentrations were determined using the Pierce BCA Protein Assay Kit (Thermo Scientific) or normalized by cell number, as indicated. Equal amounts of protein were resolved by SDS-PAGE on 4-20% gels and transferred to PVDF membranes. Membranes were blocked with 5% non-fat milk and incubated with primary antibodies overnight at 4 °C with gentle rotation, followed by incubation with HRP-conjugated secondary antibodies. Signals were detected using ECL Prime (Amersham). Where indicated, immunoblot band intensities were quantified using ImageJ/Fiji and normalized to the corresponding loading control. Reagents and antibodies are listed in **Supplementary Tables S4 and S6.**

### Patient-derived TNBC tumor and normal breast organoids

Patient-derived fresh TNBC tumor tissue and matched normal-appearing breast tissue adjacent to the tumor were collected with written informed consent under protocols approved by the Institutional Review Board at Weill Cornell Medicine (IRB #21-06023682). A live biobank of patient-derived TNBC tumor organoids is supported by the DoD Transformative Breast Cancer Consortium and the clinical trial “De-convoluting interactions between genes, the cancer environment, and the immune system to develop therapies that work for you” (DIGNITY). Patient-derived TNBC tumor and normal breast organoids were established as previously described, with minor modifications^34–37^. Fresh tissue specimens were collected in CO_2_-independent medium (Gibco) supplemented with GlutaMAX (1×; Invitrogen), Antibiotic-Antimycotic (100 U/mL; Thermo Fisher Scientific), and Primocin (100 μg/mL; InvivoGen).

Samples were washed twice with culture medium, transferred to a 10 cm Petri dish, and mechanically dissociated. Tissue fragments were then enzymatically digested in DMEM (Gibco) supplemented with penicillin (100 U/mL), streptomycin (100 μg/mL; Gibco), collagenase IV (250 U/mL; Life Technologies), Primocin (100 μg/mL), and ROCK inhibitor Y-27632 (10 μmol/L; Selleck Chemicals). Digestion was performed at 37°C with shaking at 200 rpm in a volume at least 20-fold greater than the tissue volume. Incubation time ranged from 20 to 60 min, depending on tissue amount, and was continued until the majority of cell clusters were in suspension. Advanced DMEM/F12 medium (Invitrogen) supplemented with GlutaMAX (1×), penicillin-streptomycin (100 U/mL), and HEPES (10 mM; Gibco) was added to the digested tissue suspension, followed by centrifugation at 300 × *g* for 3 min. The pellet was washed with the same medium and resuspended in the appropriate TNBC tumor or normal breast organoid culture medium. The final cell suspension was mixed with Matrigel (Corning) at a 1:2 ratio (medium: Matrigel), and 80 μL droplets (five per well) were plated in six-well suspension culture plates (Greiner Bio-One). Plates were incubated at 37°C with 5% CO_2_ for 30 min to allow Matrigel polymerization, after which 3 mL of the corresponding organoid culture medium was added to each well. Cultures were maintained with medium changes twice weekly. Organoids reaching 200–500 μm in diameter were passaged every 10–30 days, depending on the organoid line. For passaging, Matrigel droplets were dissociated with TrypLE Express (Gibco) for up to 7 min at 37°C. Cell clusters and single cells were washed and replated as described above. Organoids were cryopreserved in Recovery Cell Culture Freezing Medium (Gibco) and stored in liquid nitrogen. Cultures were routinely screened for mycoplasma contamination using a PCR-based detection kit (ABM) and confirmed negative before experimental use. Whole-exome sequencing (WES) and/or Oncomine targeted sequencing was performed on passage 5 organoid pellets and matched tumor specimens to confirm sample identity and mutational concordance. Reagents used in these experiments are listed in **Supplementary Table S8**.

### RNA sequencing and transcriptomic analysis

#### Mouse RNA sequencing and transcriptomic analysis

Libraries were prepared from total RNA using the NEBNext Ultra II Directional RNA Library Prep Kit following poly(A) enrichment with the NEBNext Poly(A) mRNA Magnetic Isolation Module (New England Biolabs). Libraries were sequenced on the Illumina NovaSeq X Plus platform to generate paired-end 2 × 51-bp reads. Raw base call (BCL) files were demultiplexed and converted to FASTQ format using Illumina bcl2fastq v2.20. Adapter sequences and low-quality bases were trimmed using Cutadapt v3.5^38^. Reads were aligned to the mouse GRCm39 reference genome using STAR v2.7.9a^39^, and gene-level read counts were generated using HTSeq-count v0.13.5^40^.

Differential expression analysis was performed using DESeq2 v1.38.3^41^. Gene set enrichment analysis (GSEA) was performed using clusterProfiler v4.6.2^42^, with genes ranked by shrunken log2 fold change, against KEGG, Reactome and MSigDB mouse Hallmark (M), curated (M2) and ontology (M5) gene sets. Enrichment results were visualized using enrichplot v1.27.5 and ggplot2 v3.4.1.

#### Patient-derived organoid RNA sequencing

Total RNA was extracted from patient-derived organoids using TRIzol (Invitrogen) and treated with DNase I using the Direct-zol RNA Miniprep Kit (Zymo Research) according to the manufacturer’s instructions. Libraries were prepared from polyadenylated RNA using the TruSeq RNA Library Prep Kit v2 or TruSeq Stranded mRNA Library Prep Kit (Illumina), according to the manufacturer’s instructions.

Libraries were sequenced on an Illumina NovaSeq 6000 platform to generate single-end 50-bp reads. Raw BCL files were demultiplexed and converted to FASTQ format using bcl2fastq v2.20 (Illumina). Adapter sequences were removed using Cutadapt v1.18. Bioinformatic processing followed the same general workflow described above, with reads aligned to the human GRCh37 reference genome using STAR v2.5.2b and gene-level counts generated using HTSeq-count v0.11.2. Normalized expression values were generated using a regularized log transformation implemented in DESeq2 v1.26.0.

### Analysis of publicly available RNA-sequencing datasets

#### Single-cell RNA-seq dataset

Publicly available single-cell RNA-sequencing data from 26 primary human breast tumors (130,246 cells passing quality control) were accessed through the Broad Institute Single Cell Portal (SCP1039; GEO accession GSE176078; EGA accession EGAS00001005173)^14^. The original study processed the raw sequencing data using Cell Ranger and identified cell-containing droplets using EmptyDrops, retaining cells with >200 detected genes, >250 unique molecular identifiers and <20% mitochondrial transcripts. Gene expression data were normalized and scaled using Seurat, followed by dimensionality reduction and unsupervised clustering. Neoplastic epithelial cells were distinguished from normal epithelial cells using inferCNV-based copy-number variation profiles. For the analysis shown in Extended Data Fig. 1, the processed Entire Breast Cancer Atlas – All cells dataset and predefined cell-type annotations were accessed through the Single Cell Portal. Expression of ACLY, ACACA, FASN, POU5F1, SOX2 and NANOG was visualized using the portal’s dot-plot function. Dot size represents the percentage of cells expressing each gene, and color intensity represents scaled mean expression.

#### TCGA and METABRIC analysis

TCGA (https://www.cancer.gov/tcga) gene expression data and associated clinical data for primary breast tumors and adjacent normal tissue were retrieved from the National Cancer Institute (NCI) Genomic Data Commons (GDC) using the TCGAbiolinks Bioconductor R package (version 2.40.0)^43–45^. Specifically, the harmonized data for gene expression quantification from the “STAR - Counts” standardized workflow were retrieved. Poor quality samples were removed, and remaining samples were filtered to include only primary breast tumor samples (n=166). Counts were normalized and differential expression was modeled with the DESeq2 pipeline (version 1.52.0)^41^. Multiple test correction was completed with the Benjamini-Hochberg method, and all differentially expressed genes reported herein meet a false discovery rate adjusted p-value of less than 0.05 (FDR < 0.05) and absolute log_2_ fold change greater than 1.0 (Ilog_2_FC| > 1.0). Differential expression rank order for Gene Set Enrichment Analysis (GSEA) was performed using the Bioconductor package fgsea (version 1.38.0) and referencing the Hallmark (H), Canonical Pathways (C2), and Ontology: GO Bioprocesses (C5) gene sets from the Molecular Signatures Database (MSigDB)^46,47^. Estimations of pathway activity in each sample were obtained by gene set variation analysis Bioconductor package (GSVA; version 2.6.6) referencing the same gene sets utilized for GSEA^48^.

A lipogenesis signature comprising FASN, ACLY, ACACA, SCD, ELOVL6, GPAM, DGAT1, and DGAT2 was scored using GSVA. Tumors with scores >0 were classified as Lipo-High and those with scores <0 as Lipo-Low. Pre-computed immune deconvolution scores for all TCGA samples were retrieved from the TIMER2.0 (https://compbio.cn/timer2/) and restricted to only the basal-like tumors analyzed above^49,50^.

For independent validation, METABRIC expression and clinical data were obtained using MetaGxBreast v1.32.0. Samples were restricted to PAM50 basal-like tumors (n = 329), and differential expression between Lipo-High and Lipo-Low tumors was analyzed using limma package (version 3.68.5)^51,52^. GSEA and GSVA were performed as described above for TCGA. Immune deconvolution of METABRIC tumors was performed using xCell implemented through immunedeconv v2.1.4. (https://github.com/omnideconv/immunedeconv) because transcripts per kilobase million (TPM) values were not available for use with other immune deconvolution methods^49,50^.

Correlations between continuous variables were assessed using Pearson correlation. Lipo-High and Lipo-Low groups were compared using Wilcoxon tests implemented in rstatix v1.1.0. P values from multiple comparisons were adjusted using the Benjamini–Hochberg procedure.

### RNA extraction and quantitative RT–PCR

Total RNA was isolated from cell lines, bulk tissues and sorted tumor cells using the RNeasy Plus Mini Kit (Qiagen) according to the manufacturer’s instructions. Approximately 250 ng RNA was reverse transcribed using qScript cDNA SuperMix (Quanta Biosciences). Quantitative RT– PCR was performed using SsoAdvanced SYBR Green Supermix (Bio-Rad) on a CFX96 Real-Time PCR System. All reactions were performed using at least three biological replicates, and specificity was confirmed by melt-curve analysis. Gene expression was normalized to GAPDH and calculated using the 2^−ΔΔC^_T_ method. Primer sequences are listed in **Supplementary Table S1.**

### LC-MS–based lipid and metabolite profiling

Free fatty acids, lipids and short-chain acyl-CoAs were analyzed by liquid chromatography– mass spectrometry (LC–MS) at the Weill Cornell Medicine Proteomics and Metabolomics Core Facility. At least three biological replicates were analyzed per condition. For tumor samples, metabolite and lipid abundance was normalized to tumor weight, whereas measurements from cultured cells were normalized to cell number. Stable-isotope-labeled lipid standards (SPLASH LIPIDOMIX, Avanti Polar Lipids) were used as internal standards for lipidomics. Free fatty acids were extracted using 90% methanol, clarified by centrifugation and dried using a SpeedVac, followed by reconstitution in 50% methanol. Chromatographic separation was performed on a Vanquish UHPLC system (Thermo Scientific) equipped with a Cadenza CD-C18 column (3 μm, 2.1 mm i.d. × 150 mm; Imtakt) coupled to a Q Exactive Orbitrap mass spectrometer (Thermo Scientific) via an Ion Max ion source with a HESI II probe. Data were acquired in negative ion mode over m/z 140–1,000 at a resolution of 70,000. Metabolites were identified based on accurate mass within 5 ppm and standard retention times, and relative abundance was determined from MS signal intensity using X Calibur v4.1. Lipids were extracted using isopropanol according to a published method^53^ and reconstituted in acetonitrile/isopropanol/water (65:30:5). Lipid profiling was performed using the same Vanquish UHPLC–Q Exactive Orbitrap platform and Cadenza CD-C18 column. Data-dependent MS/MS acquisition was performed in positive and negative ion modes, with MS1 and MS2 resolutions of 70,000 and 17,500, respectively. MS/MS data were acquired using a 1 m/z isolation window and stepped normalized collision energies of 20, 30 and 40. Lipid identification and relative quantification were performed using MS-DIAL v4.9. Short-chain acyl-CoAs were extracted using pre-cooled 90% methanol, centrifuged at 20,000g for 20 min at 4 °C, cleaned using a Sep-Pak column and dried. Targeted LC–MS analysis was performed on a Q Exactive Orbitrap mass spectrometer (Thermo Scientific) coupled to a Vanquish UPLC system using a Cadenza CD-C18 column (2.0 mm i.d. × 150 mm; Imtakt) in positive ion mode. Metabolites were identified based on accurate mass within 5 ppm and standard retention times, and relative abundance was determined from MS signal intensity using XCalibur v4.1.

### Global proteomic analysis

Mouse tumor samples were analyzed by Creative Proteomics. Proteins were extracted and digested using a filter-aided sample preparation (FASP) workflow. Briefly, tumor lysates were subjected to ultrasonic lysis and centrifugation, and proteins were processed using Microcon YM-10 centrifugal filters. Proteins were reduced with 10 mM dithiothreitol at 56 °C for 1 h and alkylated with 20 mM iodoacetamide at room temperature in the dark for 1 h. Proteins were digested overnight at 37 °C with trypsin at an enzyme-to-protein ratio of 1:50. Peptides were recovered, lyophilized and resuspended in 0.1% formic acid before LC–MS/MS analysis. Peptides were analyzed using an Easy-nLC 1000 system (Thermo Fisher Scientific) equipped with a C18 column (1.8 μm, 0.15 × 100 mm) at a flow rate of 600 nL/min. Mobile phase A consisted of 0.1% formic acid in water and mobile phase B consisted of 90% acetonitrile. The gradient was 4–7% B for 2 min, 7–25% B for 78 min, 25–35% B for 4 min and 35–90% B for 2 min. MS data were acquired using an LTQ Orbitrap ETD mass spectrometer (Thermo Fisher Scientific) over m/z 350–1,600 at a resolution of 30,000. Higher-energy collisional dissociation (HCD) was performed with an automatic gain control target of 1 × 10⁵, normalized collision energy of 35%, an isolation window of 1.0 Th and a dynamic exclusion time of 30 s. Unassigned, singly charged and >6 precursor ions were excluded. Raw mass spectrometry files were analyzed using MaxQuant v2.1 against a *Mus musculus* protein database. Trypsin specificity was used with a maximum of two missed cleavages.

Carbamidomethylation of cysteine was specified as a fixed modification, whereas methionine oxidation and N-terminal acetylation were specified as variable modifications. Precursor and MS/MS mass tolerances were set to 20 ppm and 0.05 Da, respectively. A total of 3,516 proteins were identified. Protein abundance was quantified using label-free quantification (LFQ) intensity. Protein-level q-values, peptide counts and LFQ intensities were obtained from the MaxQuant output. Differential abundance was assessed based on fold change, with proteins showing a fold change >2 or <0.5 considered upregulated or downregulated, respectively.

### Statistical analysis

Unless otherwise stated, experiments were performed using independent biological replicates. Statistical analyses were performed using GraphPad Prism v11.0.1. Comparisons between two groups were performed using two-sided unpaired Student’s t-test or Mann–

Whitney *U* tests, as appropriate. Comparisons among three or more groups were performed using ordinary one-way analysis of variance (ANOVA), with multiple-comparison tests as specified in the corresponding figure legends. Comparisons involving two independent variables were performed using ordinary two-way ANOVA with Šídák’s multiple-comparisons test. BLI signal was quantified as total photon flux (photons s⁻¹) within the defined region of interest. Statistical comparisons of BLI values were performed using GraphPad Prism. Overall survival was analyzed using the log-rank (Mantel–Cox) test. **For patient-derived organoid analyses,** comparisons between tumor-derived and normal organoids were performed using two-sided Mann–Whitney U tests, and correlations between gene-expression values were assessed using Pearson correlation. Unless otherwise stated, data are presented as mean ± s.e.m. from independent biological replicates. All statistical tests were two-sided, and P < 0.05 was considered statistically significant. **Lipidomic analysis.** Lipid intensity values were analyzed in Excel to calculate fold changes and P values. Volcano plots were generated using log₂-transformed fold changes and −log₁₀-transformed P values. **Transcriptomic analysis.** Fold changes and P values were obtained from the transcriptomic dataset. Volcano plots were generated using log₂-transformed fold changes and −log₁₀-transformed P values.

**Proteomics analysis.** Protein intensity values were obtained from the proteomics dataset and analyzed in Excel to calculate fold changes between groups. Bar graphs were generated using log_2_-transformed fold changes.

## Data Availability

RNA-sequencing data generated in this study have been submitted to the NCBI Gene Expression Omnibus (GEO) under accession numbers GSE347931(mouse) and GSE349342 (human). Proteomics data have been deposited with the ProteomeXchange Consortium via the PRIDE repository under accession number PXD084210. TCGA data are available through the NCI Genomic Data Commons, and METABRIC data were accessed as described in the Methods. Source Data are provided with this paper. All other data supporting the findings of this study are available within the Article, Extended Data and Supplementary Information or from the corresponding author upon reasonable request.

## Code availability

No custom codes were developed for this study.

## Acknowledgements

We thank J. Xiang and T. Zhang of the Genomics Resources Core Facility; J. Lively of Creative Proteomics; T. Baumgartner, P. Byrne, and M. Wallach of the Flow Cytometry Core Facility; and L. Lei, Z. Li, X. Yang, and M. Zhu of the Proteomics and Metabolomics Core Facility for technical expertise and assistance in RNA sequencing, proteomics, flow cytometry, and metabolomics. We thank S. B. Miller for animal colony management, and G. Markowitz, Y. He, and M. Martin for experimental guidance and insightful discussions. We are grateful to L. Vahdat (Dartmouth Cancer Center), E. Witze (University of Pennsylvania), E. Caron (Yale University), M. Loda (Pathology and Laboratory Medicine), M. Plasilova (NYP, Surgery), C. Vanpouille-Box and M. Martino (Radiation Oncology), and D. Gao (Cardiothoracic Surgery) for their scientific insights and discussions. We also thank members of the Cardiothoracic Surgery team, the Mittal laboratory members, and colleagues at the Meyer Cancer Center for discussions around this work.

This research was supported in part by the National Institutes of Health (R01 CA257254-01A1 to V.M.). H.W. is supported by the DRCC U24 award (U24CA274159). W.H.S. is supported by the United States Department of Defense Pilot Project Award (W81XWH-21-2-0031). Research reported in this publication was also supported by the Office of the Director of the National Institutes of Health under Award Number 1S10OD032397-01A1 to Weill Cornell Medicine. We thank the Laboratory of Comparative Pathology for histopathology support with funding from NIH Core Grant P30 CA008748-59. We are grateful to Rory and David Jones for their generous support of this study. The funders had no role in study design, data analysis, or preparation of the manuscript.

## Author contributions

M.S.K. and V.M. conceived the study. V.M. supervised the study and contributed to study design, data interpretation, and scientific direction. M.S.K. designed the study and experiments, performed the experiments, generated the experimental models, analyzed and interpreted the data, and drafted the manuscript. M.S.K. and V.M. wrote and revised the manuscript. M.S.K. collected lung tissues with assistance from A.S. P.A. and G.E.S., Jr. conducted the 3D spheroid experiments, and M.S.K. and G.E.S., Jr. analyzed the data. G.Z. conducted the lipidomics experiments, and M.S.K. and G.Z. analyzed the data. M.S.K. designed, performed and analyzed the flow-cytometry experiments; J.M. assisted with flow-cytometry acquisition. M.S.K. and L.Y. performed bioinformatics analyses of the *in vivo* datasets. M.S.K. and V.M. designed the TCGA and METABRIC analyses; H.W. performed the analyses, and V.M., M.S.K. and H.W. interpreted the results. M.S.K. and S.E.C. performed histopathological evaluations. W.H.S. provided patient-derived TNBC tumor and normal breast organoids and performed RNA sequencing. M.S.K. generated and analyzed the associated sequencing data. The Genomics Core at Weill Cornell generated next-generation sequencing libraries and performed sequencing. Creative Proteomics performed mass-spectrometry-based proteomics. H.W., J.M., S.E.C., A.S., G.E.S., Jr., G.Z. and W.H.S. contributed to manuscript writing and organization.

---

**Extended Data Fig. 1.**
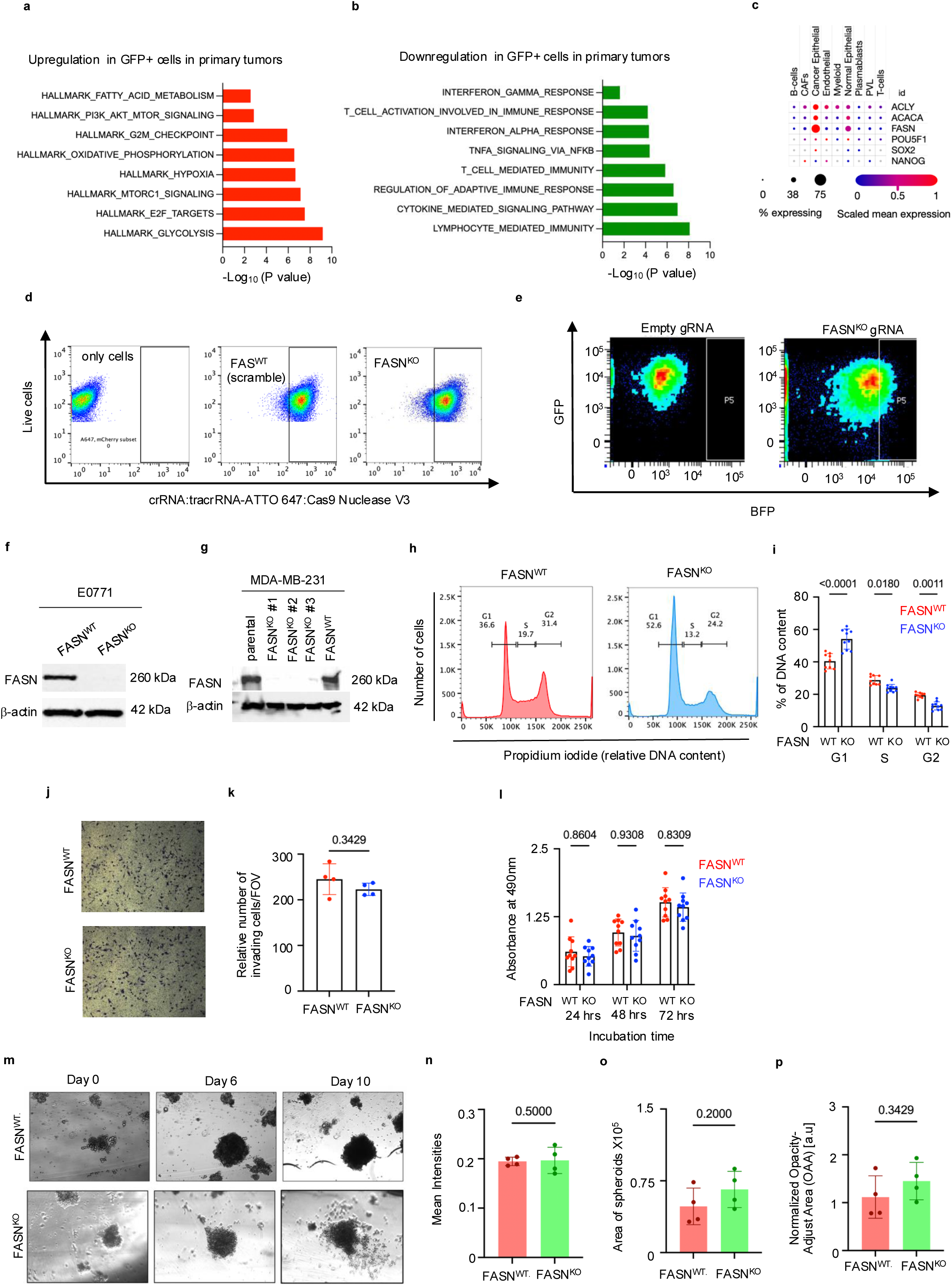
Lipogenic features of metastasis-competent tumor cells and characterization of FASN-deficient tumor models. a,b, GSEA of GFP^+^ versus GFP^-^ tumor cells isolated from orthotopic EO771-ML1 primary tumors (n = 5 biological replicates per group), showing pathways upregulated (**a**) or downregulated (**b**) in GFP^+^ cells relative to GFP^-^ cells. **c,** Single-cell RNA-sequencing analysis of human breast tumors showing expression of FASN and selected lipogenic and stemness-associated genes across major cell populations. Bubble size indicates the percentage of cells expressing each gene and color indicates scaled mean expression. **d,e,** CRISPR-Cas9-mediated FASN knockout in EO771-ML1(**d**) and MDA-MB-231-LM2 cells (**e**), with non-targeting gRNA as control. **f,g,** Immunoblot validation of FASN knockout in (**f**) EO771-ML1 and (**g**) MDA-MB-231-LM2 cells. **h,i,** Cell-cycle analysis of FASN^WT^ and FASN^KO^ cells, showing (**h**) representative DNA-content profiles and (**i**) quantification. **j,k,** Cell invasion assays of FASN^WT^ and FASN^KO^ cells, showing representative images (**j**) and quantification (**k**). **l,** Cell proliferation assay of FASN^WT^ and FASN^KO^ cells. **m–p**, 3D-bioprinted EO771-ML1 FASN^WT^ and FASN^KO^ spheroids cultured for 10 days, showing representative bright-field images at days 0, 6 and 10 (m), mean intensity (n), spheroid area (o) and normalized opacity-adjusted area (OAA) at day 10 (p) (n = 4 independent biological replicates per group). Data in panels i, k, l and n–p are presented as mean ± s.e.m. Individual data points represent biological replicates. Statistical significance for pathway enrichment in panels a and b was determined using the enrichment analysis described in the Methods. Statistical significance was assessed using ordinary two-way ANOVA with Šídák’s multiple-comparisons test for panels i and l and a two-tailed unpaired Student’s *t*-test for panel k and n-p. Exact P values are indicated in the figures.

**Extended Data Fig. 2.**
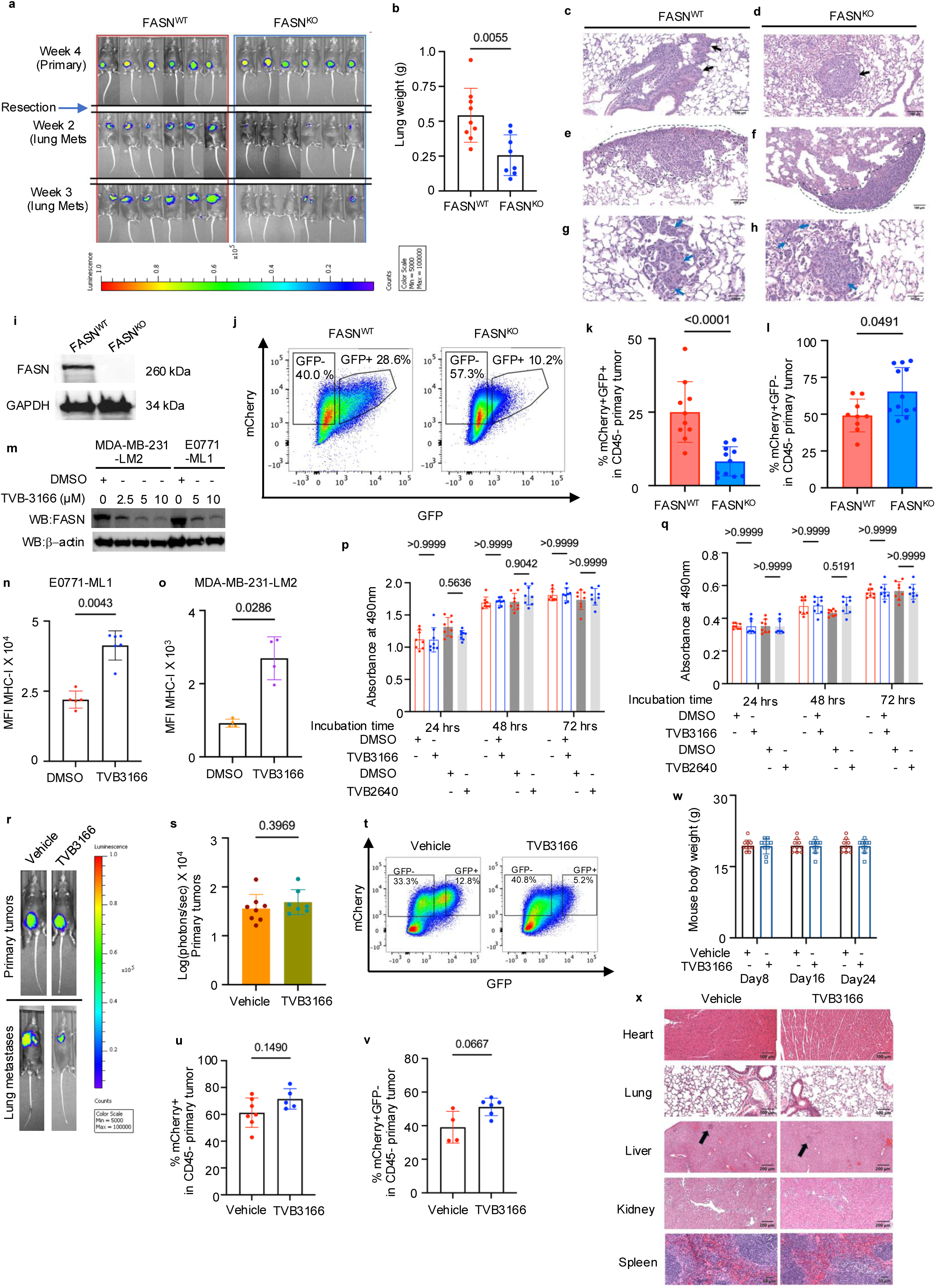
Genetic and pharmacological inhibition of FASN suppresses metastatic progression in TNBC. a-i, Characterization of primary tumor growth and metastatic progression in mice bearing FASN^WT^ or FASN^KO^ EO771-ML1 tumors. **a,** Representative bioluminescence imaging (BLI) of primary tumors and lung metastases. **b,** Lung weight. **c-h**, Representative H&E-stained lung sections showing metastatic lesions in peribronchiolar and perivascular spaces (black arrows), as well as subpleural and alveolar compartments (blue arrows). Scale bars, 50 and 100μm. **i,** Immunoblot of FASN in sorted tumor cells. **j,** Representative flow-cytometry analysis of GFP^+^ and GFP^−^ tumor cell populations isolated from E0771-ML1 primary tumors. **k,l,** Quantification of tumor-cell populations following orthotopic transplantation at a GFP^+^:GFP^-^ ratio of 2:1 (n = 10 mice per group). **m,** Immunoblot analysis of FASN in EO771-ML1 and MDA-MB-231-LM2 cells treated with TVB-3166 (0–10 μM, 72 h). **n,o,** Cell-surface MHC-I expression in EO771-ML1 (**n**) and MDA-MB-231-LM2 (**o**) following TVB-3166 treatment. **p,q,** Cell viability following treatment with TVB-3166 or TVB-2640 (10 μM, 0–72 h; n = 8 biological replicates per group). **r,s,** Representative BLI (**r**) and quantification (**s**) of primary-tumor burden in mice bearing FASN^WT^ tumors treated with vehicle or TVB-3166 (n = 7 mice per group). **t-v,** Flow-cytometric analysis of tumor-cell populations following vehicle or TVB-3166 treatment, showing representative GFP^+^ and GFP^-^ populations (**t**), total mCherry^+^ tumor cells (**u**) and (**v**) GFP^-^ tumor cells. **w,x,** Assessment of systemic tolerability following TVB-3166 treatment showing body weight over time after tumor implantation (**w**), and representative H&E-stained sections of heart, lung, liver, kidney and spleen (**x**). Arrows indicate focal single-cell hepatocellular necrosis with associated mononuclear leukocytic infiltrates in the livers of mice from both groups, consistent with background findings. The remaining tissues are histologically unremarkable. Scale bars, 100 μm (heart and lung), 200 μm (liver and kidney), and 50 μm (spleen). Data in panels b, k, l, n–q, s and u–w are presented as mean ± s.e.m. Individual data points represent biological replicates. Statistical significance was assessed using a two-tailed unpaired Student’s *t*-test for panels b, k, l, n, o, s, u and v, and ordinary two-way ANOVA with Šídák’s multiple-comparisons test for panels p, q and w. Exact P values are indicated in the figures.

**Extended Data Fig. 3.**
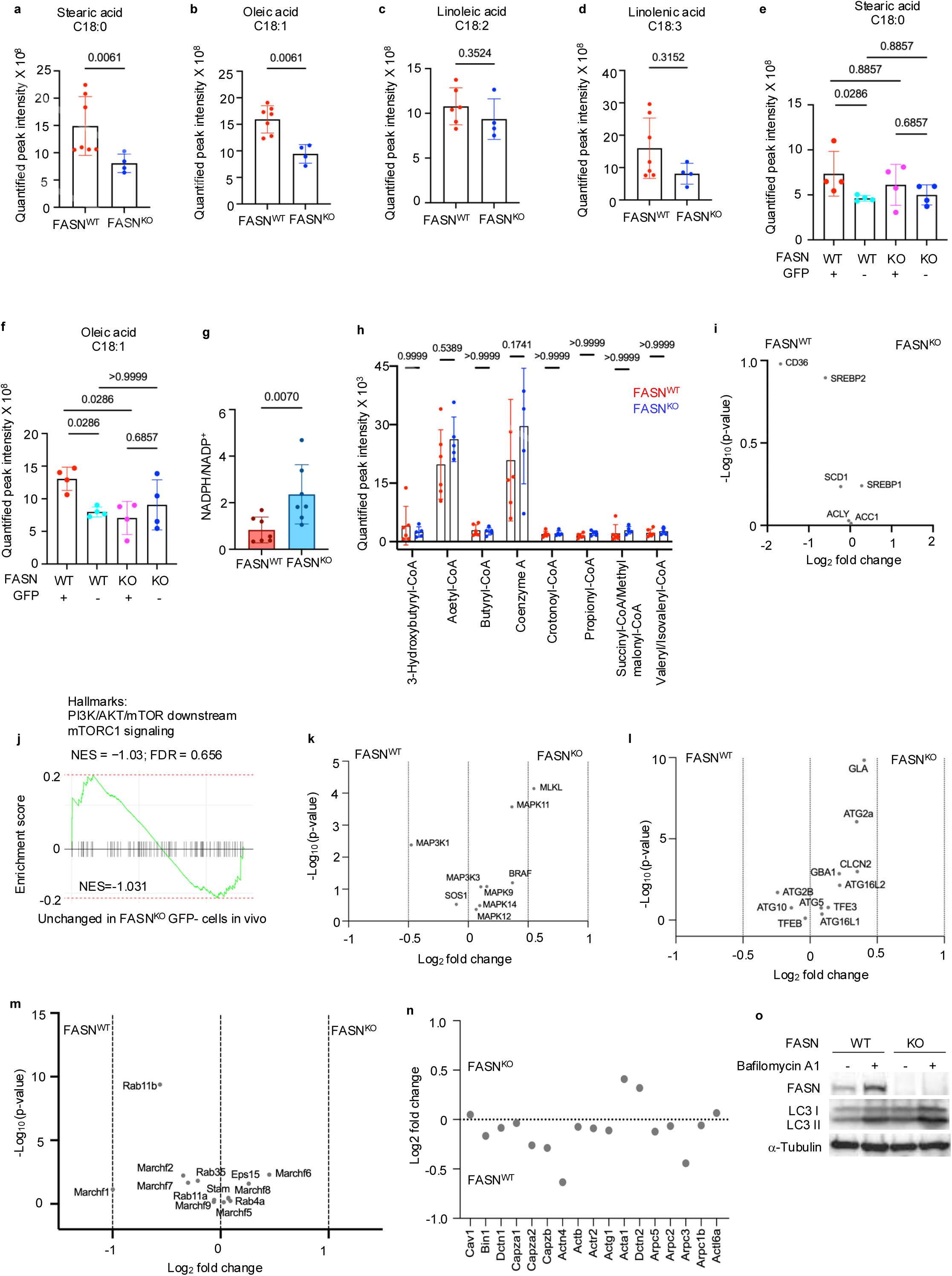
FASN-dependent lipogenesis sustains lipid abundance and PI3K–AKT– mTOR signaling. a-d, LC-MS analysis of free fatty acid abundance in FASN^WT^ and FASN^KO^ EO771-ML1 primary tumors, including (**a**) stearic acid (C18:0), (**b**) oleic acid (C18:1), (**c**) linoleic acid (C18:2) and (**d**) linolenic acid (C18:3) (n = 6 biological replicates per group). **e,f,** LC-MS analysis of (**e**) stearic acid and (**f**) oleic acid in GFP^+^ and GFP^-^ tumor cells isolated from orthotopic EO771-ML1 primary tumors (n = 4 biological replicates per group). **g,** NADPH/NADP^+^ ratio in FASN^WT^ and FASN^KO^ EO771-ML1 tumor cells. **h,** LC-MS analysis of TCA-cycle metabolites in FASN^WT^ and FASN^KO^ primary tumors (n = 6 biological replicates per group). **i-n,** Transcriptomic and proteomic analyses of FASN^WT^ and FASN^KO^ EO771-ML1 primary tumors (n = 5 biological replicates per group), showing differentially expressed genes associated with fatty acid synthesis (**i**); GSEA showing enrichment of PI3K–AKT– mTOR downstream mTORC1 signaling in GFP^-^ tumor cells from FASN^WT^ relative to FASN^KO^ tumors (**j**); differentially expressed genes associated with MAPK signaling (**k**); differentially expressed genes involved in autophagy and lysosomal function (**l**); differentially expressed genes associated with endocytic recycling and intracellular membrane trafficking (**m**); and proteomic changes in proteins associated with F-actin-related pathways (**n**). **o,** Immunoblot analysis of LC3-I and LC3-II in FASN^WT^ and FASN^KO^ MDA-MB-231-LM2 cells treated with vehicle or bafilomycin A1 (50 nM) for 16 h. Data in panels a-g are presented as mean ± s.e.m. Individual data points represent biological replicates. Statistical significance was assessed using a two-tailed Student’s *t*-test for a,b,c,d,g, ordinary one-way ANOVA with multiple-comparisons tests for e,f, and ordinary two-way ANOVA with Šídák’s multiple-comparisons test for h. GSEA enrichment statistics, including normalized enrichment scores (NES), are indicated in j. k–m, Volcano plots showing differentially expressed genes between the indicated groups. The x-axis shows log₂ fold change and the y-axis shows −log₁₀(P value). Differential expression analysis was performed using DESeq2. Exact P values are indicated in the figures.

**Extended Data Fig. 4.**
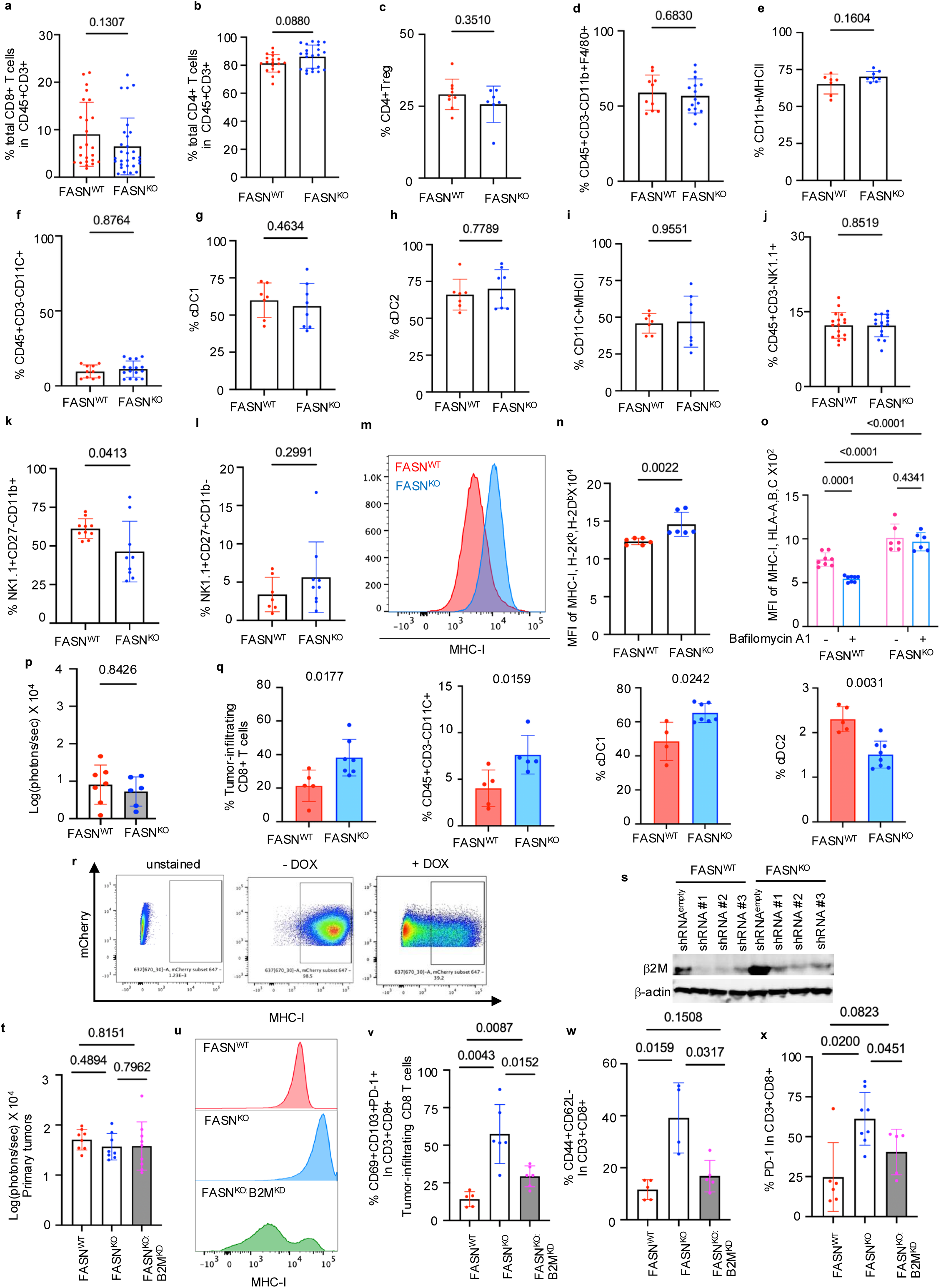
FASN loss enhances MHC-I antigen presentation without broadly altering tumor immune-cell composition. a-l, Flow-cytometric profiling of immune populations in FASN^WT^ and FASN^KO^ EO771-ML1 primary tumors (n = 8 mice per group), including (**a**) total CD8^+^ T cells; (**b**) total CD4^+^ T cells; (**c**) regulatory T (Treg) cells; (**d**) macrophages; (**e**) MHC-II^+^ macrophages; (**f**) dendritic cells; (**g**) cDC1 cells; (**h**) cDC2 cells; (**i**) MHC-II^+^ dendritic cells; (**j**) NK cells; (**k**) mature NK cells; and (**l**) immature NK cells. **m,n,** Flow-cytometric analysis of cell-surface MHC-I expression in FASN^WT^ and FASN^KO^ EO771-ML1 cells, showing (**m**) representative histograms and (**n**) quantification (n = 6 biological replicates per group). **o,** Flow-cytometric analysis of cell-surface MHC-I expression in FASN^WT^ and FASN^KO^ MDA-MB-231-LM2 cells treated with DMSO or bafilomycin A1 (50 nM) (n = 6 biological replicates per group). **p,q,** Analysis of experimental lung metastasis following intravenous injection of FASN^WT^ and FASN^KO^ EO771-ML1 cells, showing **(p)** lung metastatic burden measured by bioluminescence imaging (BLI) and **(q)** flow-cytometric profiling of tumor-infiltrating CD8^+^ T cells, dendritic cells, cDC1, and cDC2 populations. **r,s,** Generation and validation of doxycycline (DOX)-inducible B2M-knockdown EO771-ML1 cells, showing representative flow-cytometric analysis of cell-surface MHC-I following DOX treatment (**r**) and immunoblot validation of B2M knockdown (**s**). **t– x,** Analysis of DOX-inducible B2M knockdown in FASN^WT^, FASN^KO^ and FASN^KO^ B2M^KD^ orthotopic EO771-ML1 primary tumors. Mice received DOX chow following primary-tumor palpation (n = 8 mice per group). **t,** Quantification of primary-tumor BLI signal. **u**, Representative histograms of cell-surface MHC-I expression in tumor cells. **v-x**, Flow-cytometric profiling of CD8^+^ T cell populations in FASN^WT^, FASN^KO^ and FASN^KO^ B2M^KD^ orthotopic EO771-ML1 primary tumors, showing tumor-infiltrating CD8⁺ T cells (**v**), effector-memory CD8^+^ T cells (**w**) ; and PD-1^+^ CD8^+^ T cells (**x**). Data in panels a–l, n–q, t and v-x are presented as mean ± s.e.m. Individual data points represent biological replicates. Statistical significance was assessed using a two-tailed Student’s t-test for panels a–l, n, p and q, ordinary two-way ANOVA with Šídák’s multiple-comparisons test for panel o, and ordinary one-way ANOVA with multiple-comparisons tests for panels t and v-x. Exact P values are indicated in the figures.

**Extended Data Fig. 5.**
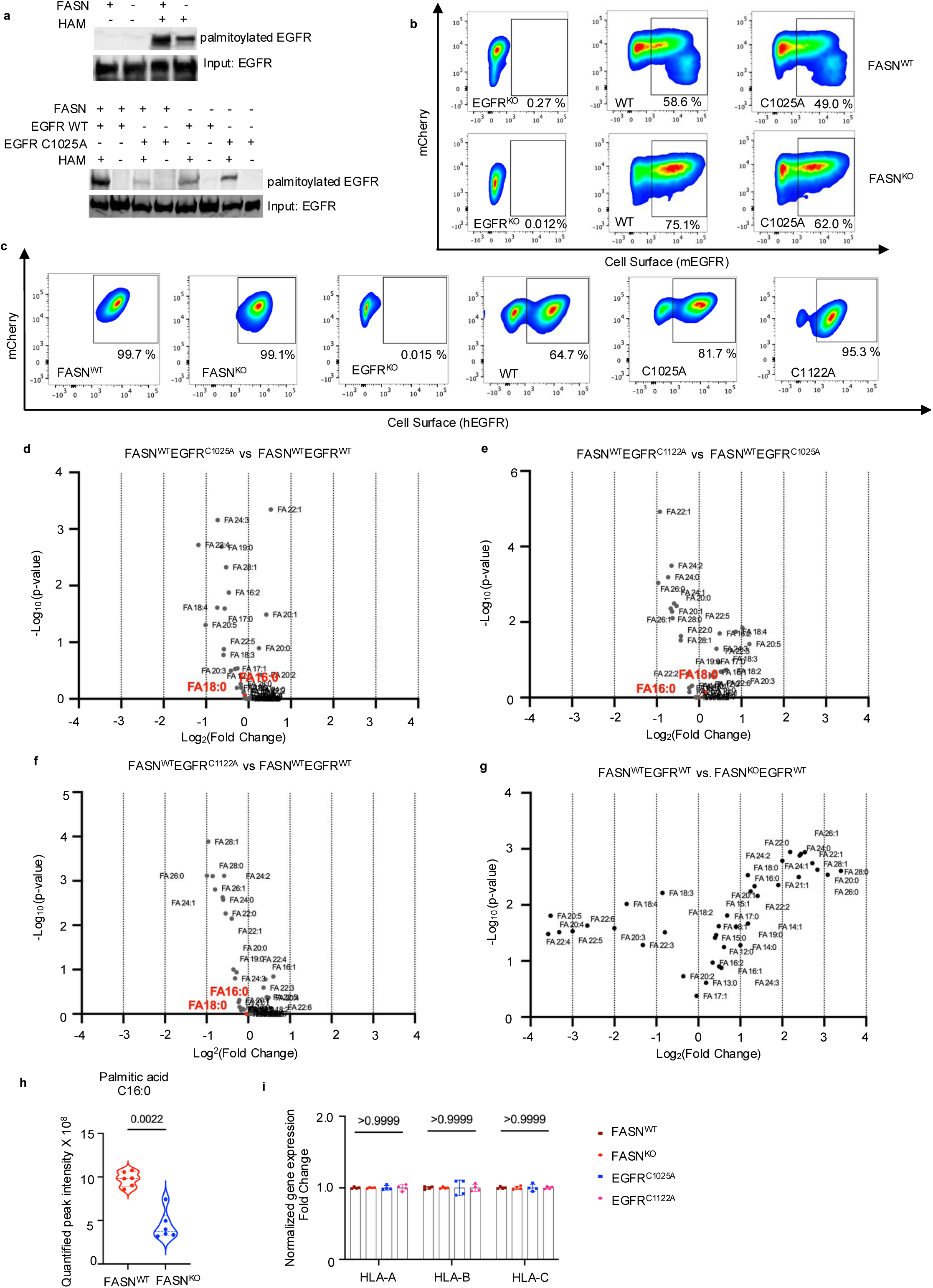
EGFR palmitoylation acts downstream of de novo lipogenesis without altering EGFR cell-surface abundance or MHC-I transcript levels. **a,** Acyl-biotin exchange (ABE) analysis of EGFR palmitoylation in EO771-ML1 cells expressing wild-type EGFR (EGFR^WT^) in the presence or absence of FASN (top) or wild-type or palmitoylation-deficient EGFR (bottom). Hydroxylamine (HAM) dependent biotinylation was used to detect palmitoylated EGFR. **b,** Representative flow-cytometry plots of cell-surface EGFR in FASN^WT^ and FASN^KO^ EO771-ML1 cells expressing EGFR^KO^, EGFR^WT^ or EGFR^C1025A^. **c,** Representative flow-cytometry plots of cell-surface EGFR in MDA-MB-231-LM2 cells expressing FASN^WT^, FASN^KO^, EGFR^KO^, EGFR^WT^, EGFR^C1025A^ or EGFR^C1122A^. **d-g,** LC-MS-based profiling of intracellular fatty acid species in MDA-MB-231-LM2 cells comparing FASN^WT^EGFR^C1025A^ versus FASN^WT^EGFR^WT^ (**d**) FASN^WT^EGFR^C1122A^ versus FASN^WT^EGFR^C1025A^(**e**) FASN^WT^EGFR^C1122A^ versus FASN^WT^EGFR^WT^(**f**) and FASN^WT^EGFR^WT^ versus FASN^KO^EGFR^WT^ (**g**) (n = 4 biological replicates per group). Cells were cultured in complete medium. **h,** LC-MS quantification of intracellular palmitic acid in FASN^WT^ and FASN^KO^ EO771-ML1 cells (n = 4 biological replicates per group). **i,** Quantitative PCR analysis of MHC-I transcripts (HLA-A, HLA-B and HLA-C) in MDA-MB-231-LM2 cells expressing FASN^WT^ or FASN^KO^ together with EGFR^WT^, EGFR^C1025A^ or EGFR^C1122A^. Data in panels h and i are presented as mean ± s.e.m. Individual data points represent biological replicates. Volcano plots showing differential lipid abundance between the two groups in panels d–g. The x-axis shows log_2_ fold change and the y-axis shows −log_10_(P value). P values were calculated using a two-tailed unpaired Student’s t-test. Statistical significance was assessed using a two-tailed Student’s t-test for h and ordinary two-way ANOVA with Šídák’s multiple-comparisons test for i. Exact P values are indicated in the figures.

**Extended Data Fig. 6.**
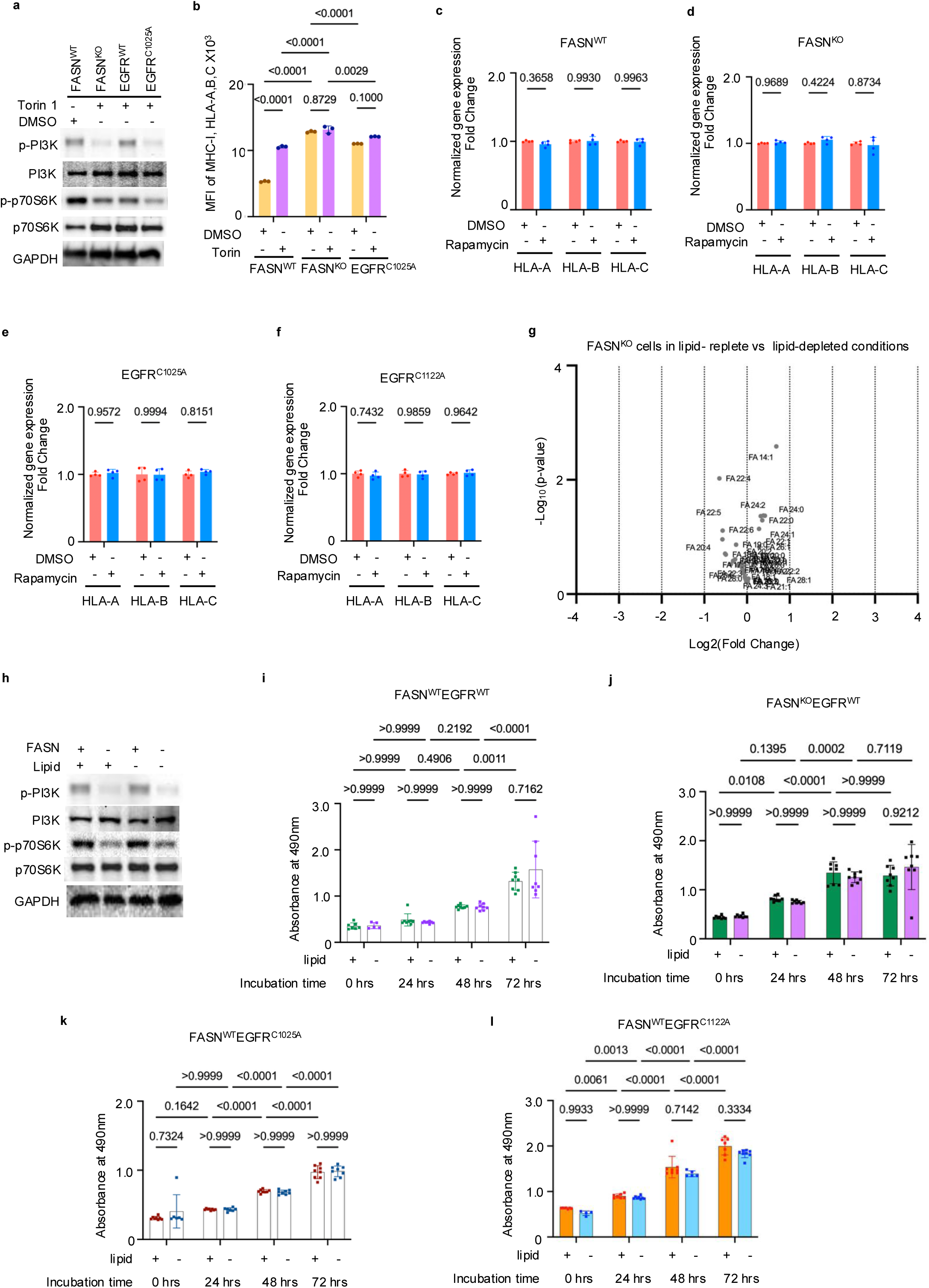
mTOR suppresses MHC-I antigen presentation through a transcription-independent mechanism downstream of EGFR palmitoylation. a,b, Effect of mTOR inhibition with Torin 1 on PI3K–mTOR signaling (**a**) and cell-surface MHC-I expression (**b**) in MDA-MB-231-LM2 cells expressing FASN^WT^ EGFR^WT^, FASN^KO^ EGFR^WT^ or FASN^WT^ EGFR^C1025A^ (n = 3 biological replicates per group). Cells were treated with DMSO or Torin (100 nM) for 48 h. **c-f,** Quantitative PCR analysis of MHC-I transcripts (HLA-A, HLA-B and HLA-C) in MDA-MB-231 cells expressing FASN^WT^ EGFR^WT^ (**c**), FASN^KO^ EGFR^WT^ (**d**), FASN^WT^ EGFR^C1025A^(**e**), and FASN^WT^ EGFR^C1122A^ (**f**) following treatment with DMSO or rapamycin (20 nM) for 48 h (n = 4 biological replicates per group). **g,** LC–MS-based profiling of free fatty acid abundance in FASN^KO^ MDA-MB-231-LM2 cells cultured in complete medium containing regular or lipid-depleted FBS (n = 4 biological replicates per group). **h,** Immunoblot analysis of PI3K-mTOR signaling in FASN^WT^ and FASN^KO^ MDA-MB-231-LM2 cells cultured in complete medium containing regular or lipid-depleted FBS. **i-l,** Cell proliferation assays of MDA-MB-231 cells cultured in complete medium containing regular or lipid-depleted FBS for 0, 24, 48 and 72 h, expressing FASN^WT^EGFR^WT^(**i**), FASN^KO^EGFR^WT^(**j**), FASN^WT^EGFR^C1025A^ (**k**) or FASN^WT^EGFR^C1122A^ (**l**) (n = 8 biological replicates per group). Data in panels b–f and i–l are presented as mean ± s.e.m. Individual data points represent biological replicates. Statistical significance was assessed using ordinary two-way ANOVA with Šídák’s multiple-comparisons test for panels b–f and i–l. Volcano plot showing differential lipid abundance between the two groups in g. The x-axis shows log_2_ fold change and the y-axis shows −log_10_(P value). P values were calculated using a two-tailed unpaired Student’s t-test. Exact P values are indicated in the figures.

**Extended Data Fig. 7.**
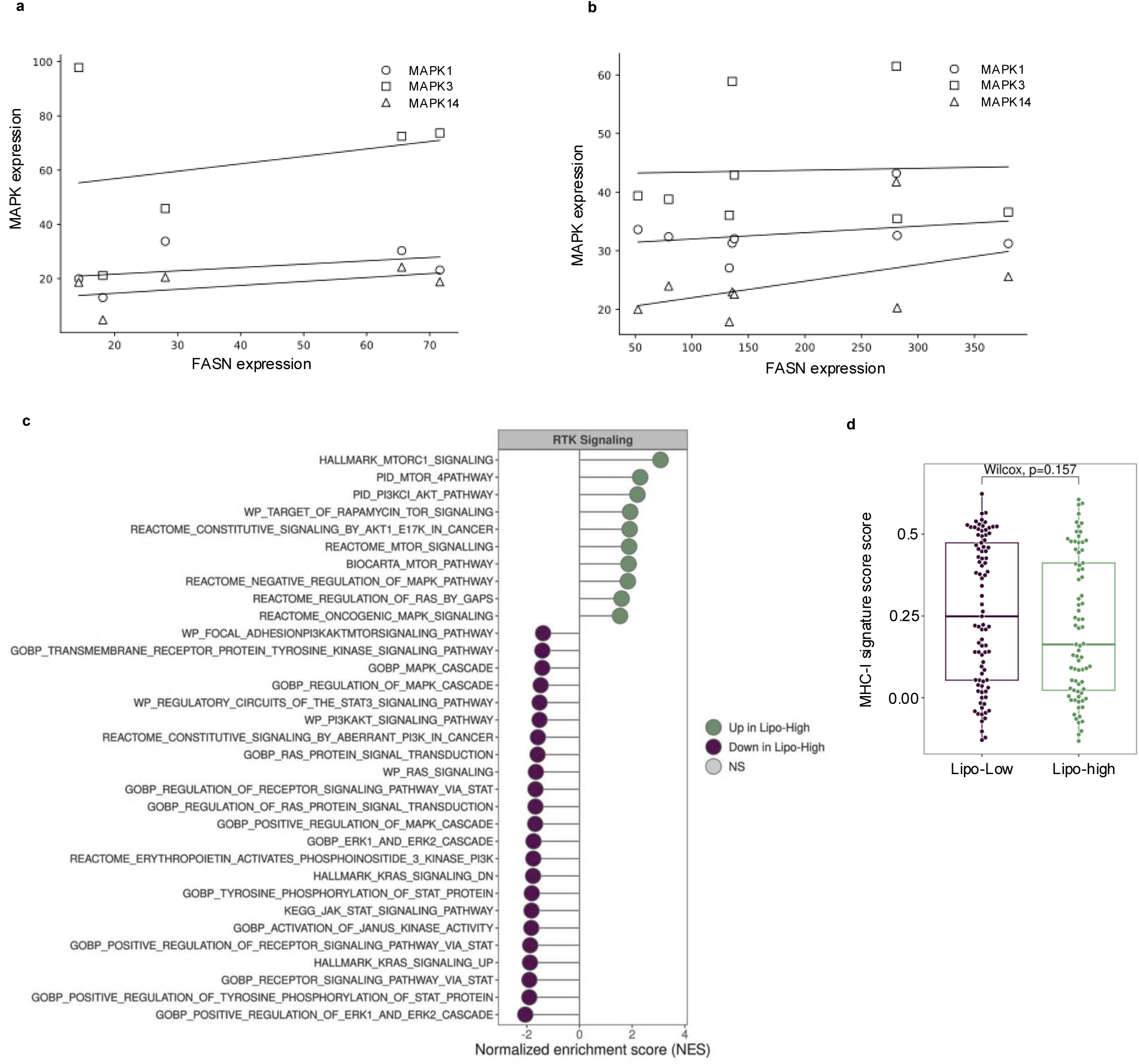
Association of lipogenesis with RTK signaling and MHC-I transcriptional signatures in human breast cancer. a,b, Correlation of FASN expression with MAPK1, MAPK3, and MAPK14 expression in normal breast organoids (a) and tumor-derived TNBC organoids (b). Lines indicate linear regression. Pearson correlation coefficients and P values are shown for MAPK1, MAPK3, and MAPK14: a, MAPK1, r = 0.404, P = 0.500; MAPK3, r = 0.252, P = 0.682; MAPK14, r = 0.531, P = 0.357; b, MAPK1, r = 0.275, P = 0.510; MAPK3, r = 0.036, P = 0.933; MAPK14, r = 0.437, P = 0.279. **c,** Gene set enrichment analysis (GSEA) of RTK downstream signaling pathways comparing lipogenesis-high and lipogenesis-low basal-like breast tumors. Normalized enrichment scores (NES) are shown for representative gene sets; filled circles indicate significantly enriched pathways (Benjamini–Hochberg-adjusted P < 0.05). **d,** MHC-I antigen-presentation transcriptional signature GSVA scores in lipogenesis-high (n = 77) and lipogenesis-low (n = 89) tumors. Scores did not differ significantly between groups (two-sided Wilcoxon rank-sum test, P = 0.157). Correlations in a,b were assessed using Pearson correlation, with r and P values shown in the figure. GSEA in c was evaluated using Benjamini–Hochberg-adjusted P < 0.05. Group comparison in d was performed using a two-sided Wilcoxon rank-sum test (P = 0.157). The corresponding NES and Benjamini–Hochberg-adjusted P values for all gene sets in c are provided in the Supplemental Data.

**Extended Data Fig. 8.**
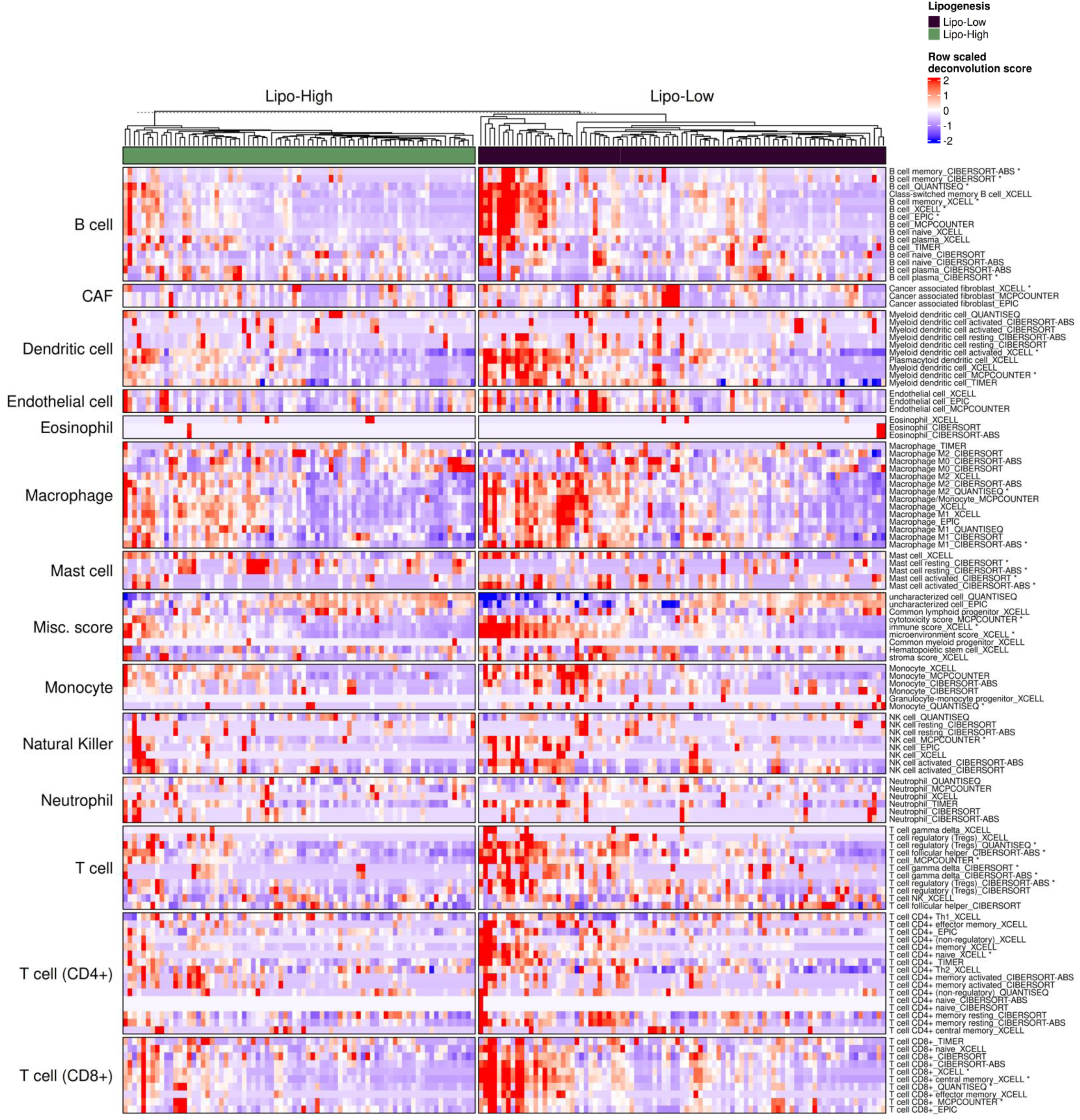
Differences in inferred immune-cell composition between lipogenesis-high and lipogenesis-low basal-like breast tumors. Heatmap of immune-cell abundance and immune-microenvironment scores inferred from multiple deconvolution algorithms, including TIMER, xCell, CIBERSORT/CIBERSORT-ABS, EPIC, MCP-counter and quanTIseq, across lipogenesis-high (n = 77) and lipogenesis-low (n = 89) basal-like breast tumors. Columns represent individual tumors and rows represent immune-cell populations or immune-related scores generated by the indicated algorithms. Values are displayed as row-scaled Z-scores. Asterisks indicate estimates that differ significantly between lipogenesis-low and lipogenesis-high tumors (two-sided Wilcoxon rank-sum test with Benjamini–Hochberg correction for multiple testing**; adjusted P < 0.05)**. Adjusted P values for all comparisons are provided in Supplementary Data Tables 3 and 6.

**Extended Data Fig. 9.**
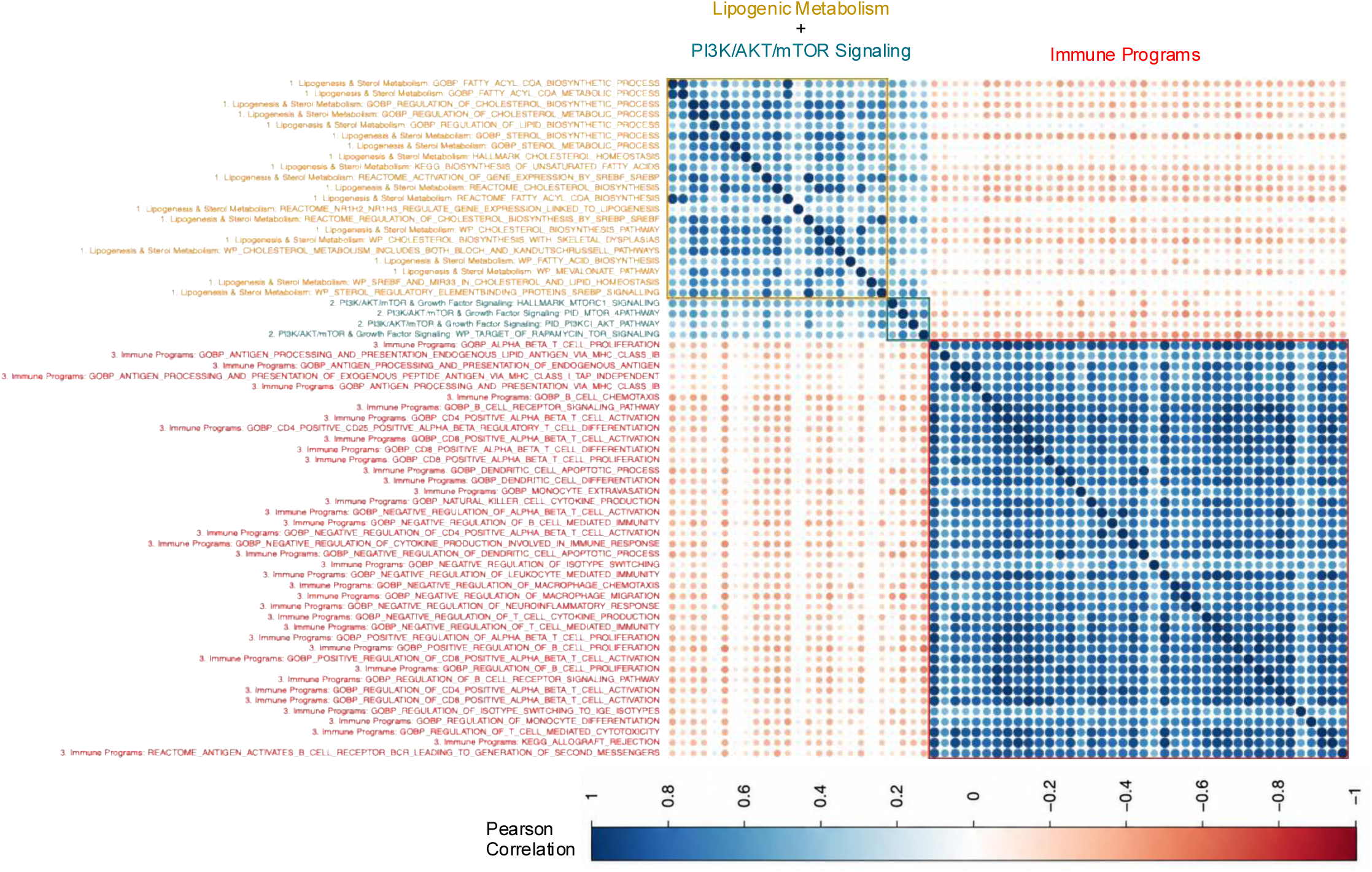
Systems-level organization of lipogenic, signaling and immune-surveillance programs in human basal-like breast cancer. Pairwise Pearson correlations among GSVA pathway signatures across basal-like breast tumors (n = 166; lipogenesis-high, n = 77; lipogenesis-low, n = 89) reveal associations among lipogenic metabolism, PI3K–AKT–mTOR signaling and immune-surveillance programs. Lipogenic and PI3K– AKT–mTOR signatures show positive correlations, whereas these programs display predominantly inverse associations with immune-surveillance programs. Boxes highlight groups of positively correlated pathway signatures and their inverse relationships with immune-surveillance signatures. Pairwise correlations were assessed using Pearson correlation, with Benjamini–Hochberg correction for multiple testing. Significant correlations were defined by adjusted P < 0.05.

**Extended Data Fig. 10.**
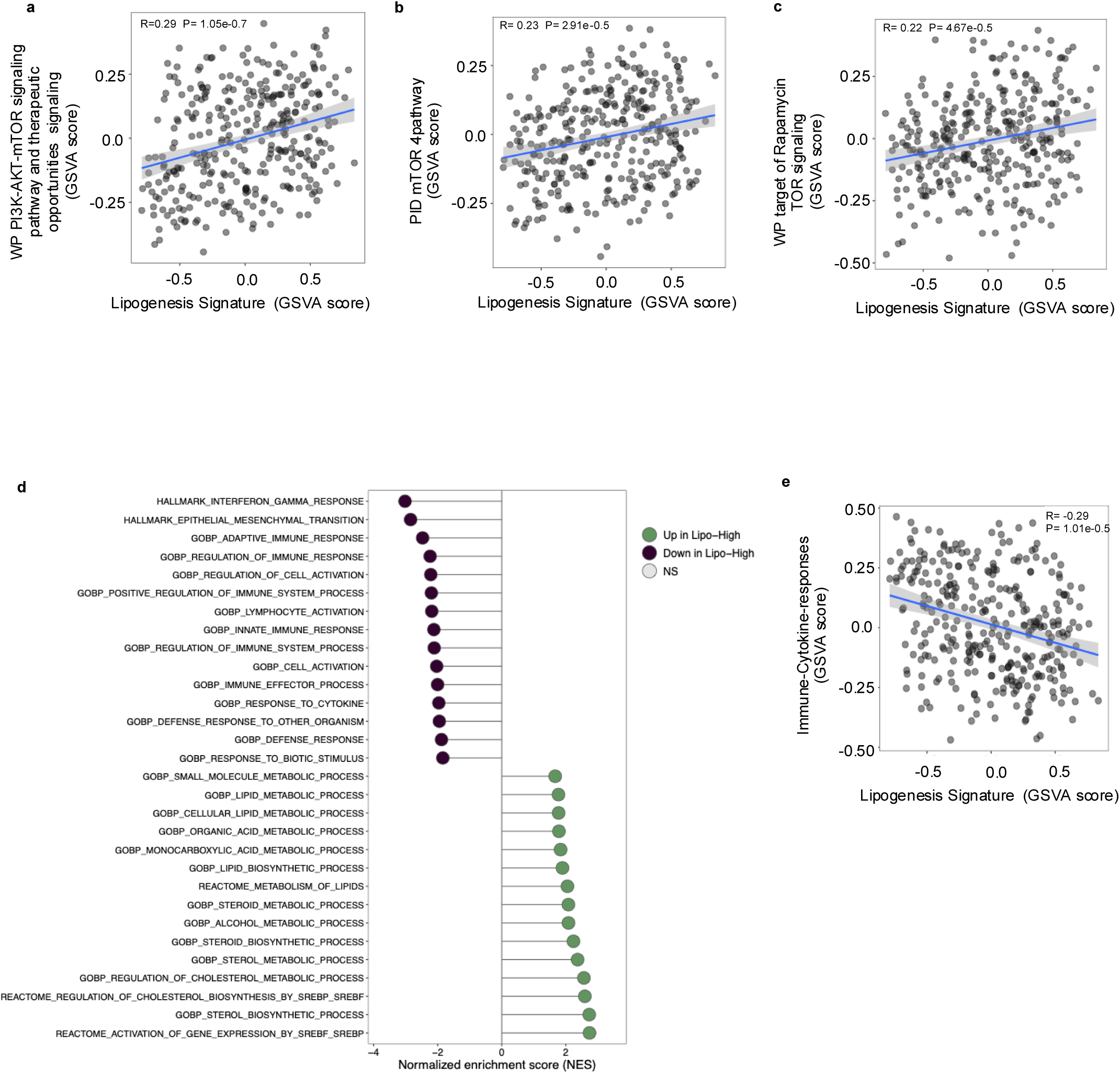
Independent validation of lipogenesis-associated signaling and immune-response signatures in METABRIC basal-like breast tumors. a–c, Correlations between the lipogenesis signature and PI3K–AKT–mTOR-related pathway GSVA scores across individual METABRIC basal-like breast tumors. Lines indicate linear regression fits with 95% confidence intervals; Pearson correlation coefficients (**r**) and P values are indicated. **d,** GSEA of immune and metabolic programs comparing lipogenesis-high (n = 175) and lipogenesis-low tumors (n = 154). Normalized enrichment scores (NES) are shown for representative gene sets; green indicates enrichment in lipogenesis-high tumors and purple indicates enrichment in lipogenesis-low tumors. Filled circles indicate significantly enriched gene sets (Benjamini–Hochberg-adjusted P < 0.05). **e,** Correlation between the lipogenesis signature and immune-cytokine-response GSVA score across individual tumors. Lines indicate linear regression fits with 95% confidence intervals; Pearson correlation coefficient (r) and P value are indicated. Exact P values for the GSEA results in d are provided in Supplementary Data Tables 7-12.

## Supplementary Tables

**Table S1.**
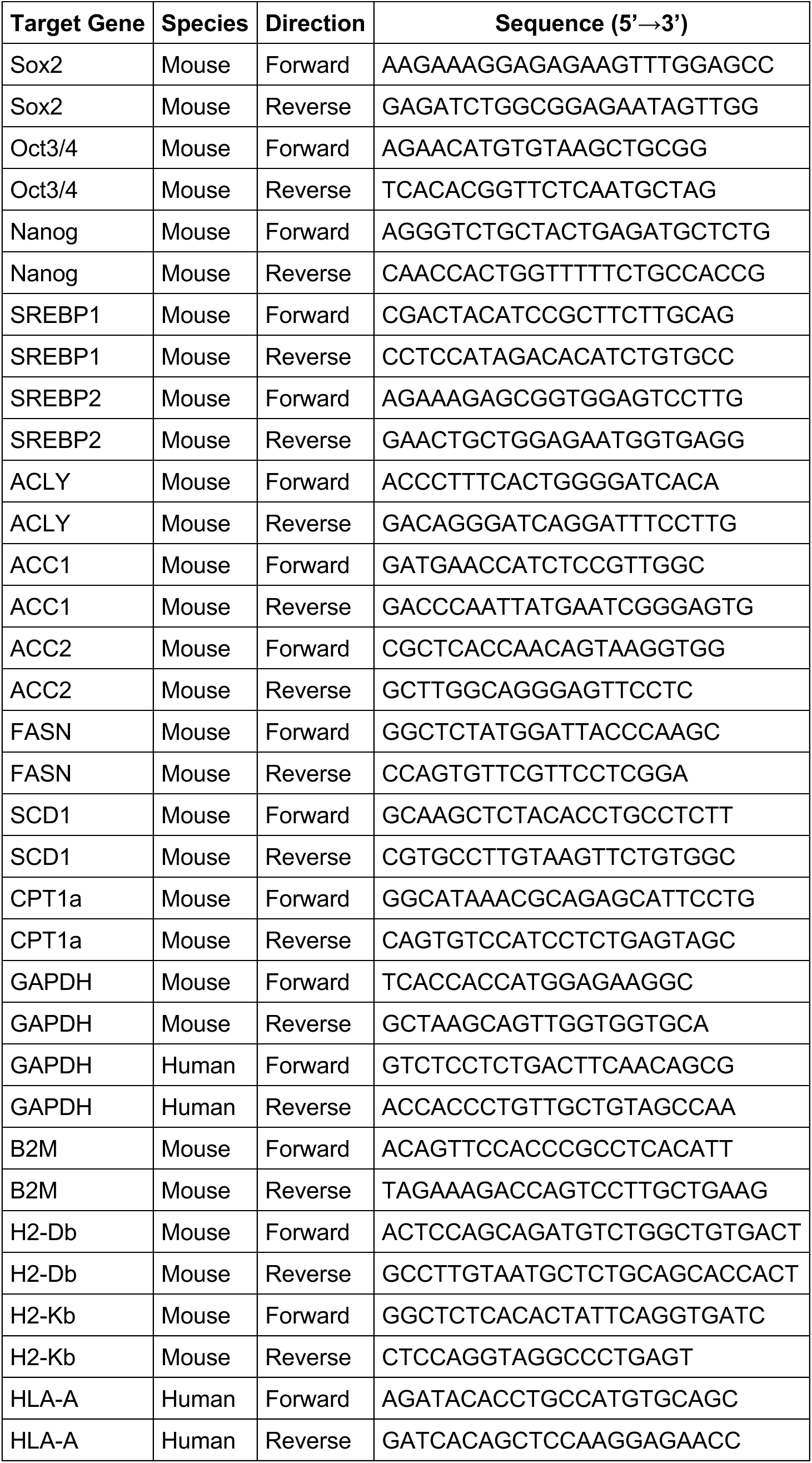
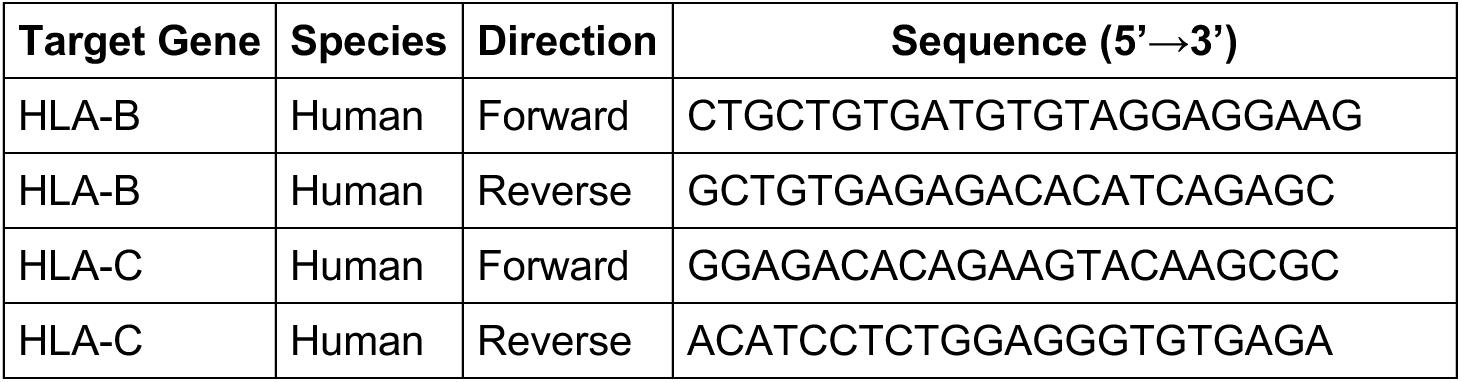
qPCR primers used in this study.

**Table S2.**
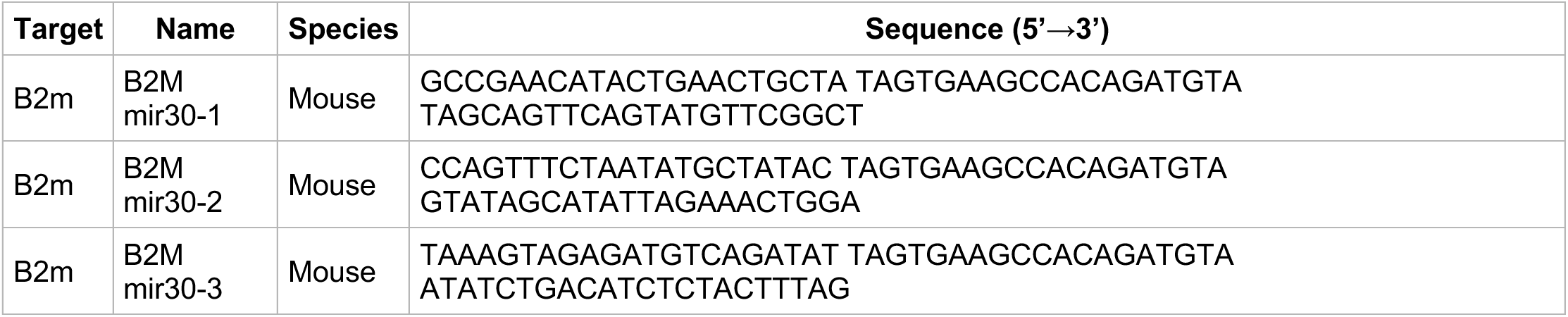
shRNA constructs.

**Table S3.**
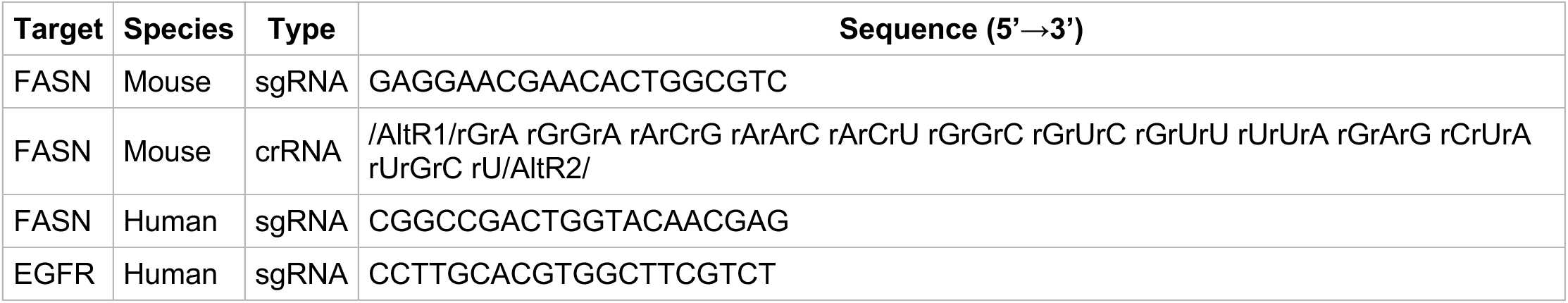
CRISPR guides.

**Table S4.**
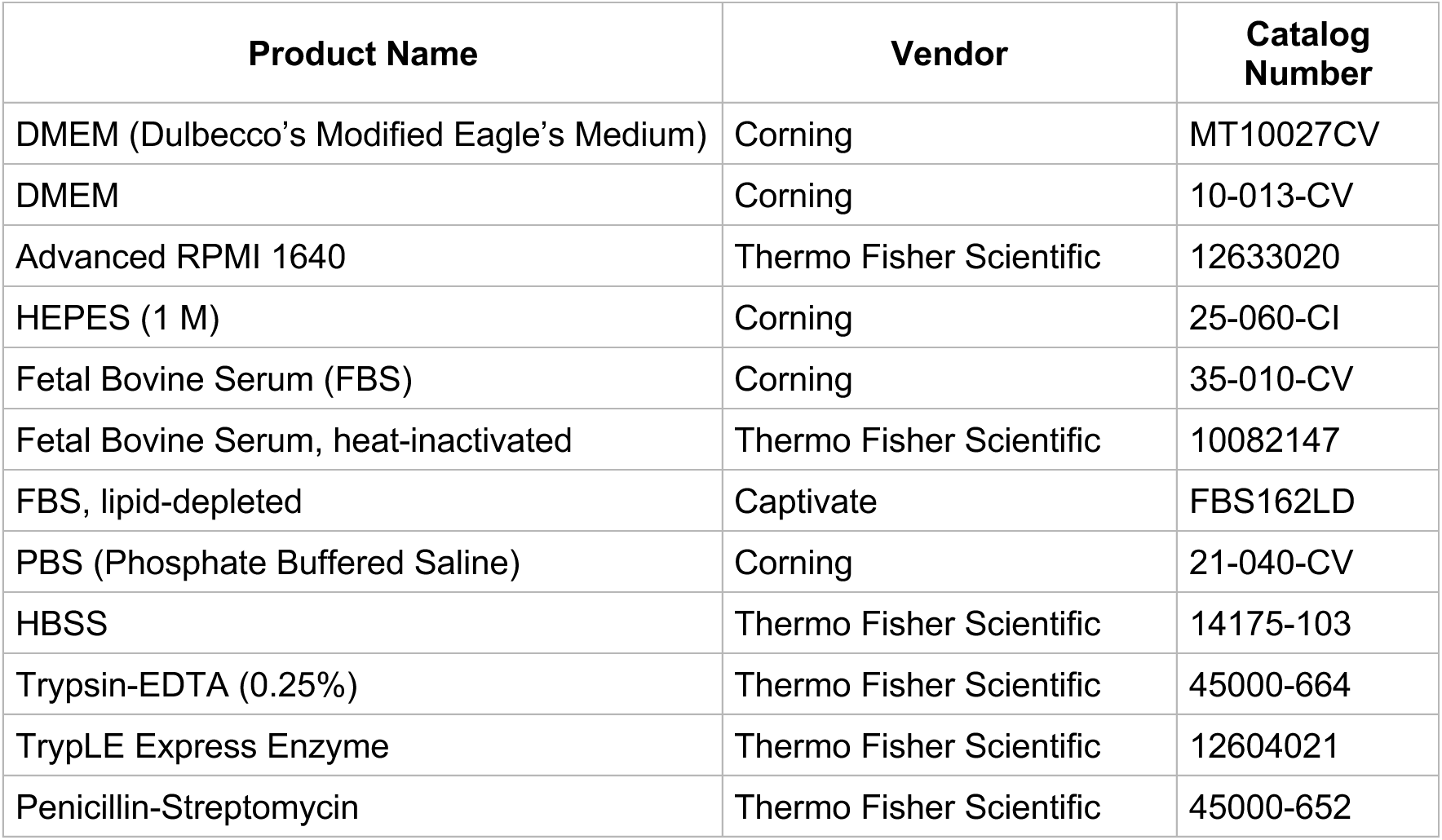
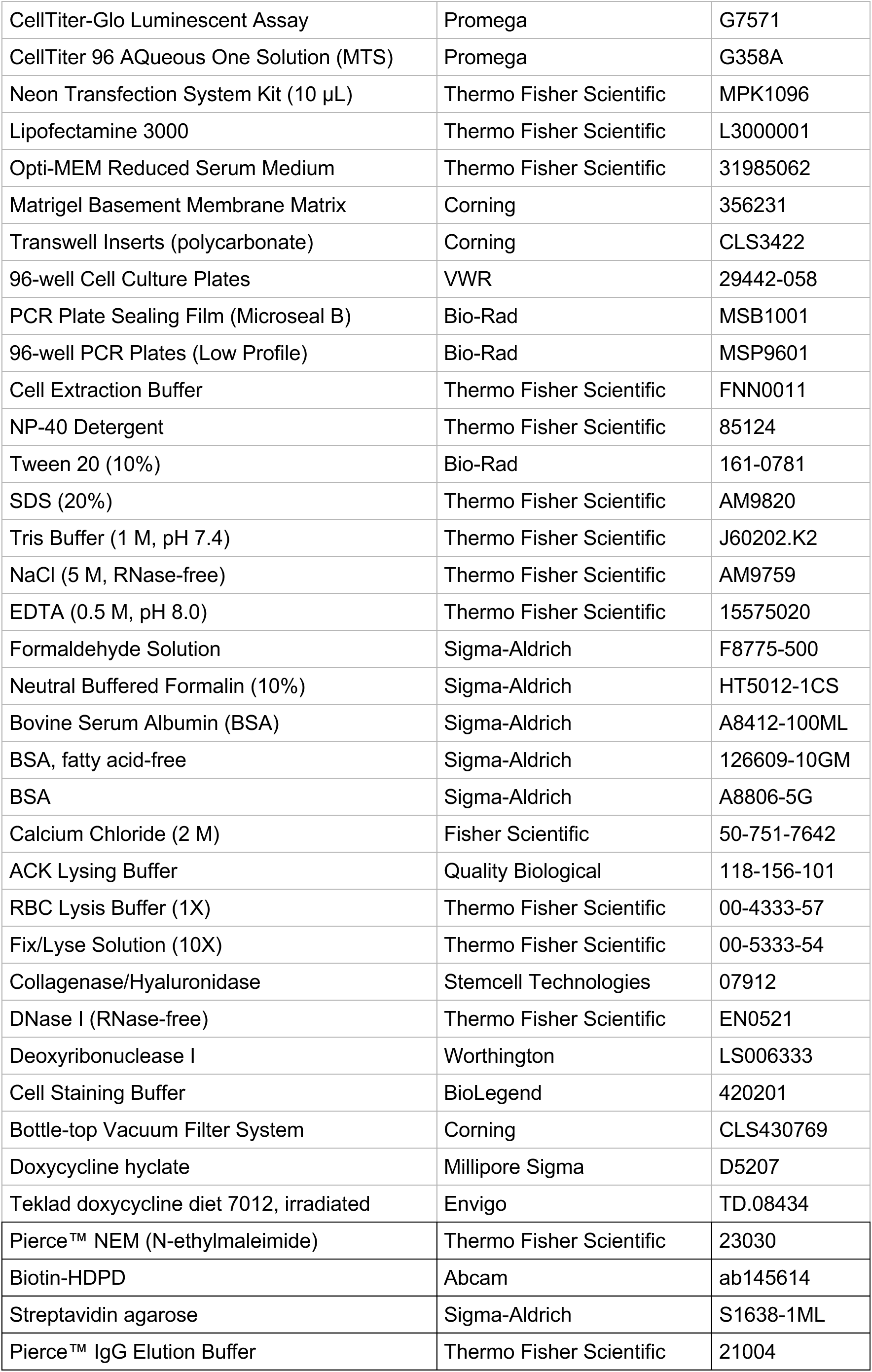
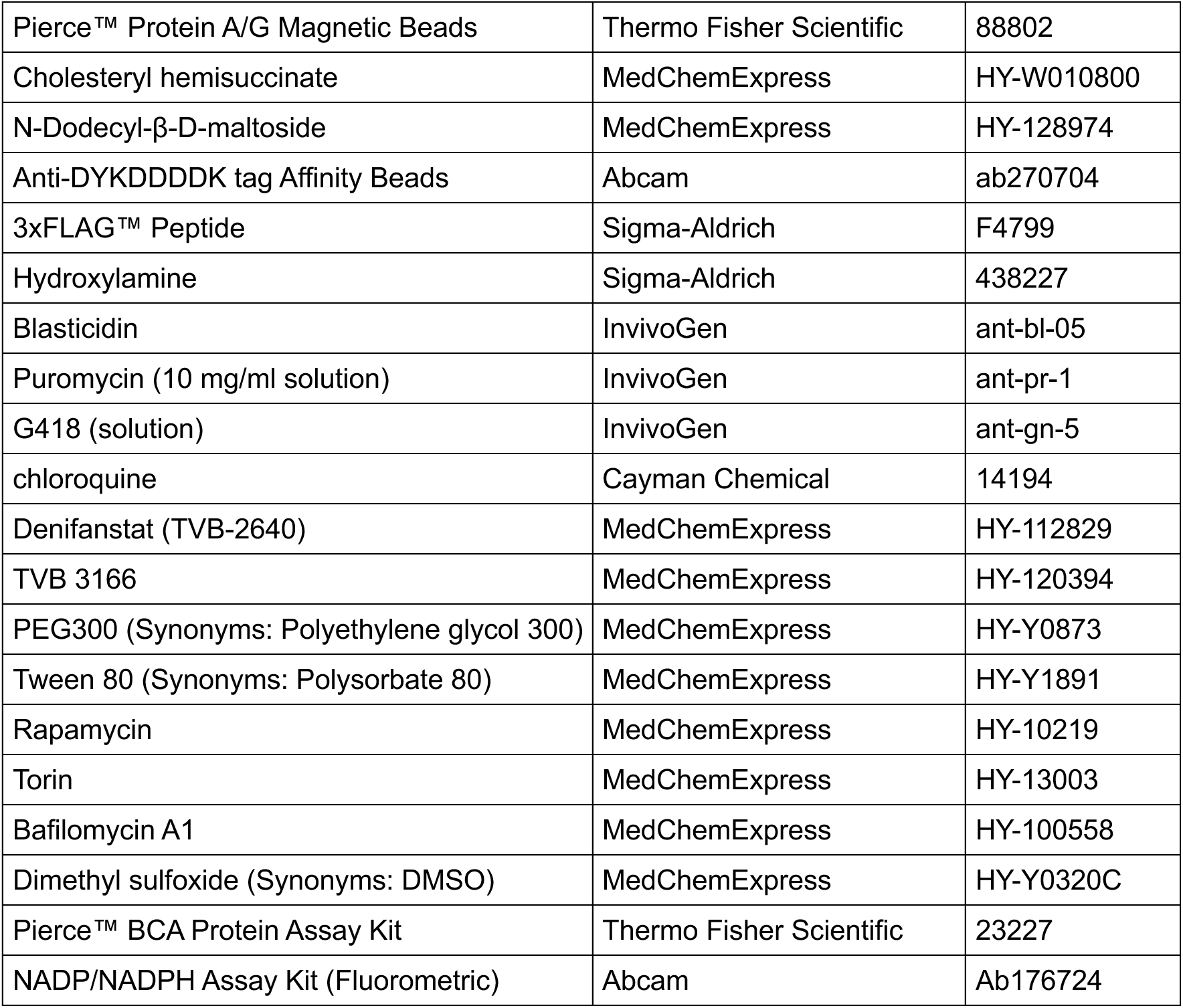
Reagents and laboratory supplies used in this study.

| Product Name | Vendor | Catalog Number |
| --- | --- | --- |
| DMEM (Dulbecco's Modified Eagle's Medium) | Corning | MT10027CV |
| DMEM | Corning | 10-013-CV |
| Advanced RPMI 1640 | Thermo Fisher Scientific | 12633020 |
| HEPES (1 M) | Corning | 25-060-CI |
| Fetal Bovine Serum (FBS) | Corning | 35-010-CV |
| Fetal Bovine Serum, heat-inactivated | Thermo Fisher Scientific | 10082147 |
| FBS, lipid-depleted | Captivate | FBS162LD |
| PBS (Phosphate Buffered Saline) | Corning | 21-040-CV |
| HBSS | Thermo Fisher Scientific | 14175-103 |
| Trypsin-EDTA (0.25%) | Thermo Fisher Scientific | 45000-664 |
| TrypLE Express Enzyme | Thermo Fisher Scientific | 12604021 |
| Penicillin-Streptomycin | Thermo Fisher Scientific | 45000-652 |

| **Product Name** | **Vendor** | **Catalog Number** |
| --- | --- | --- |
| CellTiter-Glo Luminescent Assay | Promega | G7571 |
| CellTiter 96 AQueous One Solution (MTS) | Promega | G358A |
| Neon Transfection System Kit (10 µL) | Thermo Fisher Scientific | MPK1096 |
| Lipofectamine 3000 | Thermo Fisher Scientific | L3000001 |
| Opti-MEM Reduced Serum Medium | Thermo Fisher Scientific | 31985062 |
| Matrigel Basement Membrane Matrix | Corning | 356231 |
| Transwell Inserts (polycarbonate) | Corning | CLS3422 |
| 96-well Cell Culture Plates | VWR | 29442-058 |
| PCR Plate Sealing Film (Microseal B) | Bio-Rad | MSB1001 |
| 96-well PCR Plates (Low Profile) | Bio-Rad | MSP9601 |
| Cell Extraction Buffer | Thermo Fisher Scientific | FNN0011 |
| NP-40 Detergent | Thermo Fisher Scientific | 85124 |
| Tween 20 (10%) | Bio-Rad | 161-0781 |
| SDS (20%) | Thermo Fisher Scientific | AM9820 |
| Tris Buffer (1 M, pH 7.4) | Thermo Fisher Scientific | J60202.K2 |
| NaCl (5 M, RNase-free) | Thermo Fisher Scientific | AM9759 |
| EDTA (0.5 M, pH 8.0) | Thermo Fisher Scientific | 15575020 |
| Formaldehyde Solution | Sigma-Aldrich | F8775-500 |
| Neutral Buffered Formalin (10%) | Sigma-Aldrich | HT5012-1CS |
| Bovine Serum Albumin (BSA) | Sigma-Aldrich | A8412-100ML |
| BSA, fatty acid-free | Sigma-Aldrich | 126609-10GM |
| BSA | Sigma-Aldrich | A8806-5G |
| Calcium Chloride (2 M) | Fisher Scientific | 50-751-7642 |
| ACK Lysing Buffer | Quality Biological | 118-156-101 |
| RBC Lysis Buffer (1X) | Thermo Fisher Scientific | 00-4333-57 |
| Fix/Lyse Solution (10X) | Thermo Fisher Scientific | 00-5333-54 |
| Collagenase/Hyaluronidase | Stemcell Technologies | 07912 |
| DNase I (RNase-free) | Thermo Fisher Scientific | EN0521 |
| Deoxyribonuclease I | Worthington | LS006333 |
| Cell Staining Buffer | BioLegend | 420201 |
| Bottle-top Vacuum Filter System | Corning | CLS430769 |
| Doxycycline hyclate | Millipore Sigma | D5207 |
| Teklad doxycycline diet 7012, irradiated | Envigo | TD.08434 |
| Pierce™ NEM (N-ethylmaleimide) | Thermo Fisher Scientific | 23030 |
| Biotin-HDPD | Abcam | ab145614 |
| Streptavidin agarose | Sigma-Aldrich | S1638-1ML |
| Pierce™ IgG Elution Buffer | Thermo Fisher Scientific | 21004 |
| Pierce™ Protein A/G Magnetic Beads | Thermo Fisher Scientific | 88802 |
| Cholesteryl hemisuccinate | MedChemExpress | HY-W010800 |
| N-Dodecyl-β-D-maltoside | MedChemExpress | HY-128974 |
| Anti-DYKDDDDK tag Affinity Beads | Abcam | ab270704 |
| 3xFLAG™ Peptide | Sigma-Aldrich | F4799 |
| Hydroxylamine | Sigma-Aldrich | 438227 |
| Blasticidin | InvivoGen | ant-bl-05 |
| Puromycin (10 mg/ml solution) | InvivoGen | ant-pr-1 |
| G418 (solution) | InvivoGen | ant-gn-5 |
| chloroquine | Cayman Chemical | 14194 |
| Denifanstat (TVB-2640) | MedChemExpress | HY-112829 |
| TVB 3166 | MedChemExpress | HY-120394 |
| PEG300 (Synonyms: Polyethylene glycol 300) | MedChemExpress | HY-Y0873 |
| Tween 80 (Synonyms: Polysorbate 80) | MedChemExpress | HY-Y1891 |
| Rapamycin | MedChemExpress | HY-10219 |
| Torin | MedChemExpress | HY-13003 |
| Bafilomycin A1 | MedChemExpress | HY-100558 |
| Dimethyl sulfoxide (Synonyms: DMSO) | MedChemExpress | HY-Y0320C |
| Pierce™ BCA Protein Assay Kit | Thermo Fisher Scientific | 23227 |
| NADP/NADPH Assay Kit (Fluorometric) | Abcam | Ab176724 |

**Table S5.** Flow Cytometry Reagents.

| Target / Marker | Fluorophore | Product Name | Species | Vendor | Catalog # | Dilution |
| --- | --- | --- | --- | --- | --- | --- |
| General Reagents & Buffers |  |  |  |  |  |  |
| RBC Lysis | — | 1X RBC Lysis Buffer | N/A | Thermo Fisher Scientific | 00-4333-57 | N/A |
| Staining Buffer | — | Cell Staining Buffer | N/A | Fisher Scientific | NC9742503 | N/A |
| Flow Buffer | — | Flow Cytometry Staining Buffer | N/A | Thermo Fisher | 00-4222-26 | N/A |
| Fix/Perm | — | Fixation/Permeabilization Concentrate | N/A | Thermo Fisher | 00-5123-43 | 1:10 |
| Fix/Perm | — | Fixation/Permeabilization Diluent | N/A | Thermo Fisher | 00-5223-56 | N/A |
| Permeabilization | — | Permeabilization Buffer (10X) | N/A | Thermo Fisher | 00-8333-56 | 1:10 |
| Fc Block | — | Human BD Fc Block™ | human | BD Biosciences | 564220 | 1:100 |
| Fc Block | — | TruStain FcX™ PLUS (CD16/32) | mouse | BioLegend | 156604 | 1:100 |
| Viability / Apoptosis |  |  |  |  |  |  |
| Live/Dead | 7-AAD | Viability Staining Solution | N/A | BioLegend | 420404 | 1:100 |
| Live/Dead | Zombie UV | Zombie UV | N/A | BioLegend | 423108 | 1:100 |
| DNA | DAPI | DAPI | N/A | Thermo Fisher | D1306 | 0.2 ug/mL |
| PI | — | Propidium Iodide Solution | N/A | BioLegend | 421301 | 1:100 |
| PI Kit | — | Propidium Iodide Kit | N/A | Abcam | ab139418 | N/A |
| T Cell Markers |  |  |  |  |  |  |
| CD3 | BUV395 | CD3 (17A2) | mouse | Thermo Fisher | 363-0032-82 | 1:100 |
| CD3 | RB744 | Rat Anti-Mouse CD3 | mouse | BD Biosciences | 570561 | 1:100 |
| CD4 | BUV737 | Rat Anti-Mouse CD4 | mouse | BD Horizon | 612844 | 1:100 |
| CD8a | AF488 | CD8a | mouse | BioLegend | 100723 | 1:100 |
| CD8a | BV785 | CD8a | mouse | BioLegend | 100750 | 1:100 |
| CD8a | RB780 | CD8a | mouse | BD Biosciences | 568692 | 1:100 |
| CD25 | BUV737 | CD25 (PC61.5) | mouse | Thermo Fisher | 367-0251-82 | 1:100 |
| CD25 | BV750 | CD25 | mouse | BioLegend | 102077 | 1:100 |
| CD69 | AF700 | CD69 | mouse | BioLegend | 104539 | 1:100 |
| CD69 | BUV661 | CD69 | mouse | BD Biosciences | 741478 | 1:100 |
| CD62L | BV785 | CD62L | mouse | BioLegend | 104440 | 1:100 |
| CD62L | PE-CF594 | CD62L | mouse | BD Biosciences | 562404 | 1:100 |
| CD62L | RB744 | CD62L | mouse | BD Biosciences | 570965 | 1:100 |
| CD44 | AF488 | CD44 | mouse/human | BioLegend | 103016 | 1:100 |
| CD44 | AF700 | CD44 | mouse | BioLegend | 156010 | 1:100 |
| CD44 | BV711 | CD44 | mouse/human | BioLegend | 103057 | 1:100 |
| Checkpoint / Exhaustion Markers |  |  |  |  |  |  |
| PD-1 | BV711 | CD279 (PD-1) | mouse | BioLegend | 135231 | 1:100 |
| PD-1 | BUV395 | PD-1 (J43) | mouse | Thermo Fisher | 363-9985-82 | 1:100 |
| PD-1 | PE/Dazzle 594 | PD-1 | mouse | BioLegend | 135228 | 1:100 |
| PD-1 | RB705 | PD-1 | mouse | BD Biosciences | 570566 | 1:100 |
| PD-L1 | BV711 | CD274 (PD-L1) | mouse | BioLegend | 124319 | 1:100 |
| PD-L1 | BUV737 | CD274 (MIH5) | mouse | Thermo Fisher | 367-5982-82 | 1:100 |
| LAG-3 | BV711 | CD223 (LAG-3) | mouse | BioLegend | 125243 | 1:100 |
| LAG-3 | BUV737 | CD223 | mouse | Thermo Fisher | 367-2239-42 | 1:100 |
| Tim-3 | BV785 | CD366 (Tim-3) | mouse | BioLegend | 119725 | 1:100 |
| Myeloid / Innate Markers |  |  |  |  |  |  |
| CD11b | BV605 | CD11b | mouse/human | BioLegend | 101257 | 1:100 |
| CD11c | PerCP-Cy5.5 | CD11c | mouse | BioLegend | 117328 | 1:100 |
| Ly-6C | BUV563 | Ly-6C | mouse | BD Biosciences | 755198 | 1:100 |
| Ly-6C | RB545 | Ly-6C | mouse | BD Biosciences | 569288 | 1:100 |
| Ly-6G | PerCP-Fire 806 | Ly-6G | mouse | BioLegend | 127684 | 1:100 |
| F4/80 | RB780 | F4/80 | mouse | BD Biosciences | 569223 | 1:100 |
| CD206 | AF700 | CD206 | mouse | BioLegend | 141734 | 1:100 |
| CD206 | BV711 | CD206 | mouse | BioLegend | 141727 | 1:100 |
| B / NK Cell Markers |  |  |  |  |  |  |
| B220 | BV711 | CD45R (B220) | mouse | BioLegend | 103255 | 1:100 |
| NK1.1 | BUV395 | NK1.1 | mouse | BD Horizon | 564144 | 1:100 |
| NK1.1 | BUV395 | NK1.1 (PK136) | mouse | Thermo Fisher | 363-5941-82 | 1:100 |
| Antigen Presentation / MHC |  |  |  |  |  |  |
| HLA-ABC | AF647 | HLA-A,B,C | human | BioLegend | 114612 / 311414 | 1:100 |
| HLA-ABC | PE | HLA-A,B,C | human | BioLegend | 311406 | 1:100 |
| HLA-ABC | BUV395 | HLA-ABC | human | BD Biosciences | 567859 | 1:100 |
| MHC-I | AF647 | H-2Kb/H-2Db | mouse | BioLegend | 114612 | 1:100 |
| MHC-I | BUV395 | H-2Kb/H-2Db | mouse | BD Biosciences | 745596 | 1:100 |
| SIINFEKL | AF647 | H-2Kb/SIINFEKL | mouse | BD Biosciences | 570011 | 1:100 |
| SIINFEKL | BUV395 | H-2Kb/SIINFEKL | mouse | BD Biosciences | 570093 | 1:100 |
| MHC-II | AF647 | I-A/I-E | mouse | BioLegend | 107618 | 1:100 |
| MHC-II | Pacific Blue | I-A/I-E | mouse | BioLegend | 107620 | 1:100 |
| Tumor / Epithelial Markers |  |  |  |  |  |  |
| EGFR | AF647 | EGFR (528) | mouse | Santa Cruz | sc-120 AF647 | 1:100 |
| EGFR | BV785 | EGFR | human | BioLegend | 352942 | 1:100 |
| EpCAM | BV711 | CD326 (EpCAM) | mouse | BioLegend | 118233 | 1:100 |
| CD31 | AF647 | CD31 | mouse | BioLegend | 102516 | 1:100 |
| Cytokines / Functional Markers |  |  |  |  |  |  |
| IFN-g | BUV737 | IFN-g (XMG1.2) | mouse | Thermo Fisher | 367-7311-82 | 1:100 |
| TNF-a | BV785 | TNF-a | mouse | BioLegend | 506341 | 1:100 |
| IL-2 | BV421 | IL-2 | mouse | BioLegend | 503826 | 1:100 |
| CD107a | BV711 | CD107a (LAMP-1) | mouse | BioLegend | 121631 | 1:100 |
| T-bet | BV605 | T-bet | Human/mouse | BioLegend | 644817 | 1:100 |
| Other Reagents |  |  |  |  |  |  |
| Brefeldin A | — | Brefeldin A (1000X) | N/A | BioLegend | 420601 | 1:1000 |
| Monensin | — | Monensin (1000X) | N/A | BioLegend | 420701 | 1:1000 |
| Ionomycin | — | Ionomycin (1000X) | N/A | MedChemExpress | HY-13434 | 1:1000 |
| PMA | — | Phorbol 12-myristate 13-acetate | N/A | MedChemExpress | HY-18739 | 1:1000 |
| Calibration | — | Quantum Alexa Fluor 647 MESF | N/A | Bangs Labs | 647 | N/A |
| Beads | — | Simply Cellular anti-Mouse IgG | mouse | Bangs Labs | 815 | 1:100 |
| Beads | — | Simply Cellular anti-Rat IgG | N/A | Bangs Labs | 817 | N/A |

**Table S6.** Western Blot Reagents and Antibodies.

| Target / Category | Product Name | Vendor | Catalog # | Dilution |
| --- | --- | --- | --- | --- |
| Gel Electrophoresis & Transfer Reagents |  |  |  |  |
| Gel | 4–20% Mini-PROTEAN® TGX™ Precast Gel (15-well) | Bio-Rad | 4561096 | N/A |
| Gel | 4–20% Mini-PROTEAN® TGX™ Precast Gel (10-well) | Bio-Rad | 4561094 | N/A |
| Ladder | Precision Plus Protein™ Kaleidoscope™ Standards | Bio-Rad | 1610395 | N/A |
| Membrane | Immun-Blot PVDF Membrane | Bio-Rad | 1620177 | N/A |
| Transfer | Foam Pads for Mini Trans-Blot® Cell | Bio-Rad | 1703933 | N/A |
| Transfer Buffer | 10× Tris/Glycine Blot Buffer | Bio-Rad | 1610771 | N/A |
| Running Buffer | 10× Tris/Glycine/SDS Buffer | Bio-Rad | 1610772EDU | N/A |
| Blocking & General Buffers |  |  |  |  |
| Blocking | Bovine Serum Albumin (BSA) | Sigma | A8806-5G | 5% |
| Loading Buffer | Blue Loading Buffer Pack | Cell Signaling Technology | 7722S | — |
| Wash Buffer | Tris Buffered Saline (TBS) | Santa Cruz | sc-362186 | 1× |
| Stripping | Restore™ Stripping Buffer | Thermo Fisher | PI21063 | N/A |
| Solvent | Methanol | VWR | TS42395-0025 | 20% |
| Blotting | Blotting-Grade Blocker | Bio-rad | 1706404 | 5% |
| Detection Reagents |  |  |  |  |
| Substrate | Amersham ECL Prime Detection Reagent | GE Healthcare | RPN2232 | N/A |
| Film | Development Folders | Thermo Fisher | T2258 | N/A |
| Secondary Antibodies (HRP) |  |  |  |  |
| Anti-Mouse IgG | Goat anti-Mouse IgG (H+L), HRP | Thermo Fisher | 31430 | 1:10,000 |
| Anti-Mouse IgG | Anti-Mouse IgG, HRP-linked | Cell Signaling Technology | 7076S | 1:10,000 |
| Anti-Rabbit IgG | Anti-Rabbit IgG, HRP-linked | Cell Signaling Technology | 7074S | 1:10,000 |
| Loading Controls |  |  |  |  |
| b-Actin | β-Actin (D6A8) Rabbit mAb | Cell Signaling Technology | 8457S | 1:1,000 |
| GAPDH | GAPDH (14C10) Rabbit mAb | Cell Signaling Technology | 2118S | 1:1,000 |
| α-Tubulin | α-Tubulin (11H10) Rabbit mAb | Cell Signaling Technology | 2144S | 1:1,000 |
| MHC-I |  |  |  |  |
| b2M | b-2 Microglobulin (4H5L6) | Thermo Fisher | 701250 | 1:1,000 |
| Lipid Metabolism / SREBP Axis |  |  |  |  |
| FASN | Fatty Acid Synthase Antibody | Cell Signaling Technology | 3189S | 1:1,000 |
| FASN | Fatty Acid Synthase (C20G5) Rabbit mAb | Cell Signaling Technology | 3180S | 1:1,000 |
| SREBP1 | Anti-SREBP1 | Abcam | ab28481 | 1:1,000 |
| SREBP2 | SREBP2 Polyclonal Antibody | Thermo Fisher | PA1-338 | 1:1,000 |
| Epithelial–Mesenchymal Transition (EMT) |  |  |  |  |
| E-Cadherin | E-Cadherin (24E10) Rabbit mAb | Cell Signaling Technology | 3195S | 1:1,000 |
| Vimentin | Vimentin (D21H3) Rabbit mAb | Cell Signaling Technology | 5741S | 1:1,000 |
| EGFR Signaling |  |  |  |  |
| EGFR (Total) | EGFR (D38B1) Rabbit mAb | Cell Signaling Technology | 4267S | 1:1,000 |
| EGFR (Total) | EGFR Antibody (A-10) | Santa Cruz | sc-373746 | 1:1,000 |
| EGFR (pY1068) | Phospho-EGFR (Tyr1068) (D7A5) | Cell Signaling Technology | 3777S | 1:1,000 |
| EGFR (pY1068) | Phospho-EGFR (Y1068) | Abcam | ab5644 | 1:1,000 |
| PI3K–AKT–mTOR Pathway |  |  |  |  |
| PI3K p85 (Total) | PI3K p85 Antibody | Cell Signaling Technology | 4292S | 1:1,000 |
| PI3K p85 (pTyr458) | Phospho-PI3K p85/p55 | Cell Signaling Technology | 17366S | 1:1,000 |
| Akt (Total) | Akt Antibody | Cell Signaling Technology | 9272S | 1:1,000 |
| Akt (pSer473) | Phospho-Akt (Ser473) | Cell Signaling Technology | 9271S | 1:1,000 |
| mTOR (Total) | mTOR (7C10) Rabbit mAb | Cell Signaling Technology | 2983S | 1:1,000 |
| mTOR (pSer2448) | Phospho-mTOR (Ser2448) | Cell Signaling Technology | 5536S | 1:1,000 |
| p70 S6K (Total) | p70 S6 Kinase | Cell Signaling Technology | 9202S | 1:1,000 |
| p70 S6K (pThr389) | Phospho-p70 S6 Kinase | Cell Signaling Technology | 9205S | 1:1,000 |
| MAPK Signaling |  |  |  |  |
| ERK1/2 (Total) | p44/42 MAPK (Erk1/2) | Cell Signaling Technology | 9102S | 1:1,000 |
| ERK1/2 (pThr202/Tyr204) | Phospho-ERK1/2 | Cell Signaling Technology | 9101S | 1:1,000 |
| p38 (Total) | p38 MAPK | Cell Signaling Technology | 9212S | 1:1,000 |
| p38 (pThr180/Tyr182) | Phospho-p38 MAPK | Cell Signaling Technology | 9211S | 1:1,000 |
| Energy Sensing |  |  |  |  |
| AMPKα (Total) | AMPKα Antibody | Cell Signaling Technology | 2532S | 1:1,000 |
| AMPKα (pThr172) | Phospho-AMPKα | Cell Signaling Technology | 2535S | 1:1,000 |
| Autophagy Flux |  |  |  |  |
| LC3B Polyclonal Antibody | LC3I and LC3II | Thermo Fisher | PA1-46286 | 1:1,000 |
| Tags / Miscellaneous |  |  |  |  |
| FLAG tag | Anti-DDDDK tag | Abcam | ab205606 | 1:1,000 |

**Table S7.** DNA Cloning and Molecular Biology Reagents.

| Category | Product Name | Vendor | Catalog # |
| --- | --- | --- | --- |
| Plasmid Preparation & DNA Purification |  |  |  |
| Endotoxin-free plasmid prep | NucleoBond Xtra Midi Plus EF | MACHEREY-NAGEL | 740422-5 |
| Plasmid miniprep | Monarch® Plasmid Miniprep Kit | NEB | T1010S |
| PCR cleanup | Monarch® PCR & DNA Cleanup Kit | NEB | T1030S |
| Gel extraction | Monarch® DNA Gel Extraction Kit | NEB | T1020S |
| RNA isolation | RNeasy Plus Mini Kit | Qiagen | 74134 |
| DNase treatment | DNase I | Worthington | LS006333 |
| PCR Amplification & Mutagenesis |  |  |  |
| High-fidelity PCR | Q5® Hot Start High-Fidelity Master Mix | NEB | M0494S |
| High-fidelity PCR | iProof HF Master Mix | Bio-Rad | 1725310 |
| Site-directed mutagenesis | Q5® Site-Directed Mutagenesis Kit | NEB | E0552S |
| Additive | DMSO | VWR | 97063-136 |
| DNA Assembly & Ligation |  |  |  |
| DNA assembly | NEBuilder® HiFi DNA Assembly Kit | NEB | E5520S |
| Ligation kit | Quick Ligation™ Kit | NEB | M2200S |
| DNA ligase | T4 DNA Ligase | NEB | M0202S |
| Dephosphorylation | Quick CIP | NEB | M0525S |
| Phosphorylation | T4 Polynucleotide Kinase | NEB | M0201S |
| Restriction Enzymes |  |  |  |
| Type IIS enzyme | BsmBI-v2 | NEB | R0739S |
| Restriction enzyme | SpeI-HF | NEB | R3133S |
| Restriction enzyme | XbaI | NEB | R0145S |
| Bacterial Transformation & Culture |  |  |  |
| Competent cells | NEB® 5-alpha Competent *E. coli* | NEB | C2987H |
| Competent cells | NEB® Stable Competent *E. coli* | NEB | C3040H |
| Competent cells | One Shot™ Stbl3™ *E. coli* | Thermo Fisher | C737303 |
| Selection antibiotic | Ampicillin sodium salt | Thermo Fisher | 611770250 |
| Selection antibiotic | Chloramphenicol | MilliporeSigma | C0378-25G |
| Buffers & Core Reagents |  |  |  |
| Water | Nuclease-Free Water | Invitrogen | AM9937 |
| Buffer | Tris (1.0 M, pH 7.4) | Thermo Fisher | J60202.K2 |
| Salt | NaCl (5 M, RNase-free) | Thermo Fisher | AM9759 |
| Detergent | NP-40 Surfact-Amps™ | Thermo Fisher | 85124 |
| Detergent | SDS (20% solution) | Thermo Fisher | AM9820 |
| cDNA Synthesis & qPCR |  |  |  |
| Reverse transcription | qScript cDNA SuperMix | Quanta Biosciences | 95048-500 |
| qPCR master mix | SsoAdvanced™ SYBR® Green Supermix | Bio-Rad | 1725274 |
| DNA stain | SYBR™ Safe DNA Gel Stain | Thermo Fisher | S33102 |
| General Laboratory Reagents |  |  |  |
| Solvent | 2-Propanol | VWR | BT216010-4X1L |
| Solvent | Ethanol (absolute) | MilliporeSigma | E7023-6X500ML |

**Table S8.**
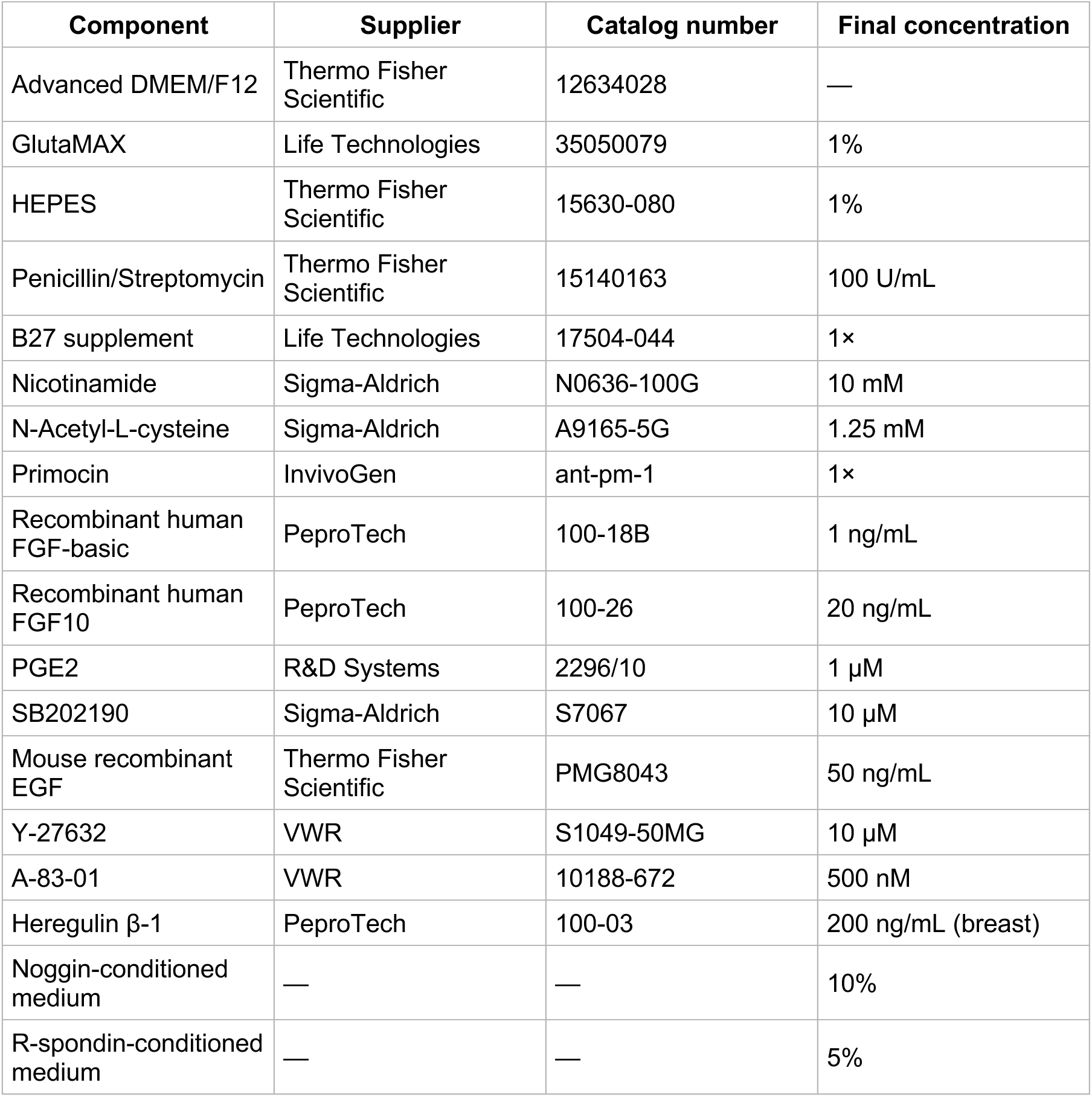
PDTO medium composition.

| Component | Supplier | Catalog number | Final concentration |
| --- | --- | --- | --- |
| Advanced DMEM/F12 | Thermo Fisher Scientific | 12634028 | — |
| GlutaMAX | Life Technologies | 35050079 | 1% |
| HEPES | Thermo Fisher Scientific | 15630-080 | 1% |
| Penicillin/Streptomycin | Thermo Fisher Scientific | 15140163 | 100 U/mL |
| B27 supplement | Life Technologies | 17504-044 | 1× |
| Nicotinamide | Sigma-Aldrich | N0636-100G | 10 mM |
| N-Acetyl-L-cysteine | Sigma-Aldrich | A9165-5G | 1.25 mM |
| Primocin | InvivoGen | ant-pm-1 | 1× |
| Recombinant human FGF-basic | PeproTech | 100-18B | 1 ng/mL |
| Recombinant human FGF10 | PeproTech | 100-26 | 20 ng/mL |
| PGE2 | R&D Systems | 2296/10 | 1 µM |
| SB202190 | Sigma-Aldrich | S7067 | 10 µM |
| Mouse recombinant EGF | Thermo Fisher Scientific | PMG8043 | 50 ng/mL |
| Y-27632 | VWR | S1049-50MG | 10 µM |
| A-83-01 | VWR | 10188-672 | 500 nM |
| Heregulin β-1 | PeproTech | 100-03 | 200 ng/mL (breast) |
| Noggin-conditioned medium | — | — | 10% |
| R-spondin-conditioned medium | — | — | 5% |

